# Spatiotemporally flexible subnetworks reveal the quasi-cyclic nature of integration and segregation in the human brain

**DOI:** 10.1101/2021.06.09.447672

**Authors:** Marika Strindberg, Peter Fransson, Joana Cabral, Ulrika Ådén

## Abstract

Though the organization of functional brain networks is modular at its core, modularity does not capture the full range of dynamic interactions between individual brain areas nor at the level of subnetworks. In this paper we present a hierarchical model that represents both flexible and modular aspects of intrinsic brain organization across time by constructing spatiotemporally flexible subnetworks. We also demonstrate that segregation and integration are complementary and simultaneous events. The method is based on combining the instantaneous phase synchrony analysis (IPSA) framework with community detection to identify a small, yet representative set of subnetwork components at the finest level of spatial granularity. At the next level, subnetwork components are combined into spatiotemporally flexibly subnetworks where temporal lag in the recruitment of areas within subnetworks is captured. Since individual brain areas are permitted to be part of multiple interleaved subnetworks, both modularity as well as more flexible tendencies of connectivity are accommodated for in the model. Importantly, we show that assignment of subnetworks to the same community (integration) corresponds to positive phase coherence within and between subnetworks, while assignment to different communities (segregation) corresponds to negative phase coherence or orthogonality. Together with disintegration, i.e. the breakdown of internal coupling within subnetwork components, orthogonality facilitates reorganization between subnetworks. In addition, we show that the duration of periods of integration is a function of the coupling strength within subnetworks and subnetwork components which indicates an underlying metastable dynamical regime. Based on the main tendencies for either integration or segregation, subnetworks are further clustered into larger meta-networks that are shown to correspond to combinations of core resting-state networks. We also demonstrate that subnetworks and meta-networks are coarse graining strategies that captures the quasi-cyclic recurrence of global patterns of integration and segregation in the brain. Finally, the method allows us to estimate in broad terms the spectrum of flexible and/or modular tendencies for individual brain areas.

## 1 Introduction

It is well established that spontaneous brain activity exhibits highly reproducible modular topological patterns of co-activation. These static representations of co-activity are known as intrinsic or resting-state networks (RSNs) and they reflect the strongest modular tendencies of connectivity over time in resting-state fMRI (Biswal et al., 1995; Kiviniemi et al., 2000; Raichle et al., 1996). However, during shorter time intervals, the core RSNs do not fully account for the dynamic interplay between individual brain regions and RSNs. (Allen et al., 2014; Betzel et al., 2016a, 2014; Chang and Glover, 2010; Chen et al., 2015; Deco et al., 2011; Leonardi et al., 2014; Majeed et al., 2011; Ponce-Alvarez et al., 2015; Zalesky et al., 2014). Rather, most brain areas are likely to participate in multiple subnetwork constellations across different modules with varied frequency. A salient example is the cingulate cortex. It consist of highly connected hub areas (Hagmann et al., 2008) that each has been shown to interact with multiple subnetworks during shorter time periods (Allen et al., 2014; Chang and Glover, 2010; Hutchison et al., 2013b; Karahanoʇlu and Van De Ville, 2015). Zones of instabilities, i.e. areas with high variability in connectivity preferences across time, is another example (Allen et al., 2014). While both flexibility and modularity of brain regions describe fundamental aspects of functional brain organization, they are somewhat contradictory. Moreover, the method used for defining subnetworks will inevitably influence which aspect of functional brain organization that will be emphasized (Keilholz et al., 2017). For example, in static parcellation schemes, overlap in space is fully avoided whereas for spatial ICA, the degree of spatial overlap is minimized (Beckmann and Smith, 2004; Calhoun et al., 2001; Kiviniemi et al., 2003). However, these approaches to define subnetworks have in common a tendency to introduce a bias towards networks characterized by a high degree of modularity.

Other approaches have been developed that allow spatially overlapping subnetworks while maximizing temporal independence (Smith et al., 2012). The resulting spatially partly overlapping temporal modes (TMFs) did not only reflect the variable connectivity profiles of different areas. It also highlighted the anticorrelated nature of simultaneous subnetwork activity, which has been previously shown using seed-based correlation analysis (Fransson, 2005). Even though Smith et al. (2012) at the time expressed some caution regarding the significance, later work has corroborated the ubiquitous nature of anti-correlations in temporally overlapping components (Karahanoʇlu and Van De Ville, 2015). Despite this progress in the understanding of time-resolved organization of intrinsic functional brain activity, much is still unknown. For example, few studies have investigated the time point by time point change in organization of functional connectivity without some initial averaging strategy across time points that limits the temporal resolution (Allen et al., 2014). Two methods that do not suffer from this limitation are IPSA (instantaneous phase synchrony analysis) (Glerean et al., 2016) and iCAP (innovation-driven co-activation patterns) (Karahanoʇlu and Van De Ville, 2015). While the latter uses the total activation (TA) – framework (Karahanoğlu et al., 2013) on the unfiltered BOLD-signal to derive spatiotemporally partly overlapping patterns of co-activations, the former is dependent on eliminating riding waves in the signal in order to derive meaningful instantaneous phase information. This is commonly achieved by applying a narrow bandpass filter. However, it does not only come at the cost of inducing potentially meaningful loss of variability in the signal. It is also still very much an open question which frequency domains in the BOLD-signal that are coupled to neuronal activity (Niazy et al., 2011). Another important question that remains unanswered is how to best accommodate modularity and flexibility simultaneously in the time-resolved domain as outlined above. This problem is closely related to the concepts of integration and segregation that are central to dynamic brain organization. Segregation refers to a state where connectivity within distinct modules is strong while links between modules are weak. Integration refers to the opposite situation where connectivity across modules are equally strong or stronger than within modules (Shine and Poldrack, 2018; Sporns, 2013). However, this definition usually leaves little room for the event when a module temporarily ceases to be coherent enough to be referred to as a unit. It is also still unclear how transitions between integration and segregation take places.

One approach to brain organization that in theory can harbor the complementary and contradictory aspects of modularity and flexibility is that of modeling the brain as a dynamical system. In this model framework, all areas are continuously changing relationships with each other along some gradient. The gradients of coupling strengths between areas are determined by the underlying anatomical wiring and the current configuration of the system (Deco et al., 2011). The coupling strengths correspond to the edges in a network. In addition, the concept of meta-stability has the explanatory power to describe the forces that drive the internal dynamics. In meta-stability, areas quasi-synchronize transiently to form temporary functional units where stronger coupling results in longer duration of co-operation (Kelso et al., 2013; Scott Kelso, 2012; Tognoli and Kelso, 2014). If no configuration acts as a fixed point, the dynamics is instead constrained by the gradual movement between a small number of quasi-stable principal configurations (Cavanna et al., 2018; Tognoli and Kelso, 2014; Varela et al., 2001). These so-called quasi-attractors are abstract generalizations of the strongest common coupling tendencies. The tension between the quasi-attractors is what gives momentum to the dynamics.

In this work we propose a novel method inspired by the global view of dynamical systems to derive spatiotemporally flexible subnetworks from BOLD fMRI resting-state data. At its core, our model is based on idea of metastable couplings between brain regions. In contrast to a full description of a dynamical system, our method does not consider every brain area at every time point but rather focus on the strongest coupled constellations of brain areas across different spatial granularities to study the temporal dynamics of network integration and segregation. Importantly, our hierarchical spatiotemporal approach reduces the full dynamical system into a subset of interleaved subnetworks that capture network activity both within and across borders of modularity. Therefore, both of the fundamentally different tendencies of brain areas, that is, to be either strongly and repeatedly connected to a small number of other areas (commonly referred to by the concept of modularity) or, to be briefly and flexibly connected to a large set of areas (i.e. flexibility), are thereby in part accommodated for by our model. Of note, the suggested method captures the time point to time point change in functional connectivity as well as the segregated nature of functional brain organization. Furthermore, our analysis suggests the presence of quasi-attractors that shape the quasi cyclic nature of global connectivity in the brain. It also suggests that integration and segregation co-exist along the temporal domain in the brain at different levels of spatial granularity.

## 2 Material and methods

### 2.1 Data and acquisition

From the HCP 100 unrelated Young Adult subject release www.humanconnectome.org/study/hcp-young-adult, sixty-one individuals who met the criterium of low motion in both the left-to-right (rfMRI_REST1_LR) and right-to-left (rfMRI_REST1_RL) rs-fMRI acquisitions were included (Smith et al., 2013; Van Essen et al., 2012). Low motion was defined as a maximum peak in FDRMS (framewise displacement root mean square) time series < 1 mm together with a mean FDRMS < 0.1 mm. Only the rfMIR_REST1_LR dataset was used in the subsequent analyses. Each 4D EPI dataset consisted of 1200 volumes with TR = 720 msec. The total acquisition time corresponded to 864 sec (approximately 14,5 min). Other MR acquisition parameters and further details can be found in the publications from the HCP - consortium (Uğurbil et al., 2013). The collection of the data was approved by the Washington University institutional review board and is made publicly available (https://db.humanconnectome.org).

### 2.2 Preprocessing and parcellation

The dataset had been extensively preprocessed using the FIX-pipeline to remove scanner related artefacts as well as artefacts related to motion and physiological noise. Details of the preprocessing steps can be found in the HCP-publications (Smith et al., 2013). No further pre-processing of the data was conducted. For cortical parcellation the Shaefer 200 brain area parcellation (7 Networks) was used (Schaefer et al., 2018). Additionally, the subcortical portion of the Human Brainnetome Atlas (Fan et al., 2016) was used to parcellate the thalamus, basal ganglia, hippocampus and amygdala. In total, the parcellation scheme used comprised 236 brain areas. The parcellation of the amygdala and hippocampus were assigned to the “limbic network” in the Schaefer parcellation scheme while the thalamus and basal ganglia were treated as separate networks. Hence, our analysis therefore considered nine separate core resting-state networks.

### 2.3 Overview of the method to compute spatiotemporally flexible subnetworks

The aim of the hierarchical method presented below was to account for both modular and flexible tendencies of connectivity in the brain, as well as capture changes in integration and segregation across time. It was based on combining the IPSA (Instantaneous phase synchrony analysis) framework (Glerean et al., 2016) with the Louvain community detection algorithm (Blondel et al., 2008). This allowed community assignment to be associated with coupling strength in terms of phase coherence. Instead of choosing a narrow band-pass filter, the empirical mode decomposition (EMD) was used to identify the intrinsic mode function (IMF) that was best fitted to the classical frequency range [0.01-0.1Hz] of spontaneous BOLD-signal fluctuations (Huang et al., 1998; Niazy et al., 2011). Here, the concepts of integration, segregation and disintegration between network building blocks were defined based on phase coherence relationships within and between communities. Our method combined spatially and temporally interleaved building blocks into flexible subnetworks. The core assumption was that areas that were frequently assigned to the same community had a strong phase coupling across time. This assumption was also subsequently tested and verified. Though hierarchical in nature, our model was built from the bottom up without any a-priori assumptions about assignment of brain areas to specific networks. The proposed model represented the most salient intrinsic relationships in the brain in terms of spatiotemporally flexible and interleaved subnetworks at four levels of spatial granularity: combinations of two areas (level 1), subnetwork components (SNCs, level 2), subnetworks (SNs, level 3) and meta-networks (MN, level 4). First, the strongest pairwise relationships from the point of view of each brain area were calculated in terms of the highest frequency of assignment to the same community (q_int_). The pairs were then used as seeds in an iterative search to identify larger SNCs (up to a total of eight areas). SNCs were subsequently grouped into spatiotemporally flexible SNs that were, by definition, always internally coherent, i.e. assigned to the same community. Since a large number of SNs were found to be spatially partly overlapping, SNs were furthered clustered into meta-networks (MN). The MNs were thus a way to summarize spatiotemporally highly interleaved SNs into larger networks that for most (but not all) time points were internally coherent, i.e. the constituting SNs were assigned to the same community. To test how well the procedure of gradually building larger networks units (SNCs, SNs and MNs) performed as a hierarchical strategy to capture the global pattern of time resolved amplitude fluctuations, we constructed time resolved state vectors. These state vectors represented the time-resolved “global state” of the brain in terms of the combined amplitude fluctuations from all SNs and MNs respectively at each time point. The degree of global state recurrence as a function of time was estimated by correlating state vectors at different time points. Recurrence was then compared with estimates based on brain areas independently of network membership. Finally, we applied our method to estimate the relative degree of flexibility versus modularity of individual brain areas. A schematic overview of the key steps that are performed in our method to derive spatiotemporally flexible networks is provided in Figure 1.

**Figure 1.**
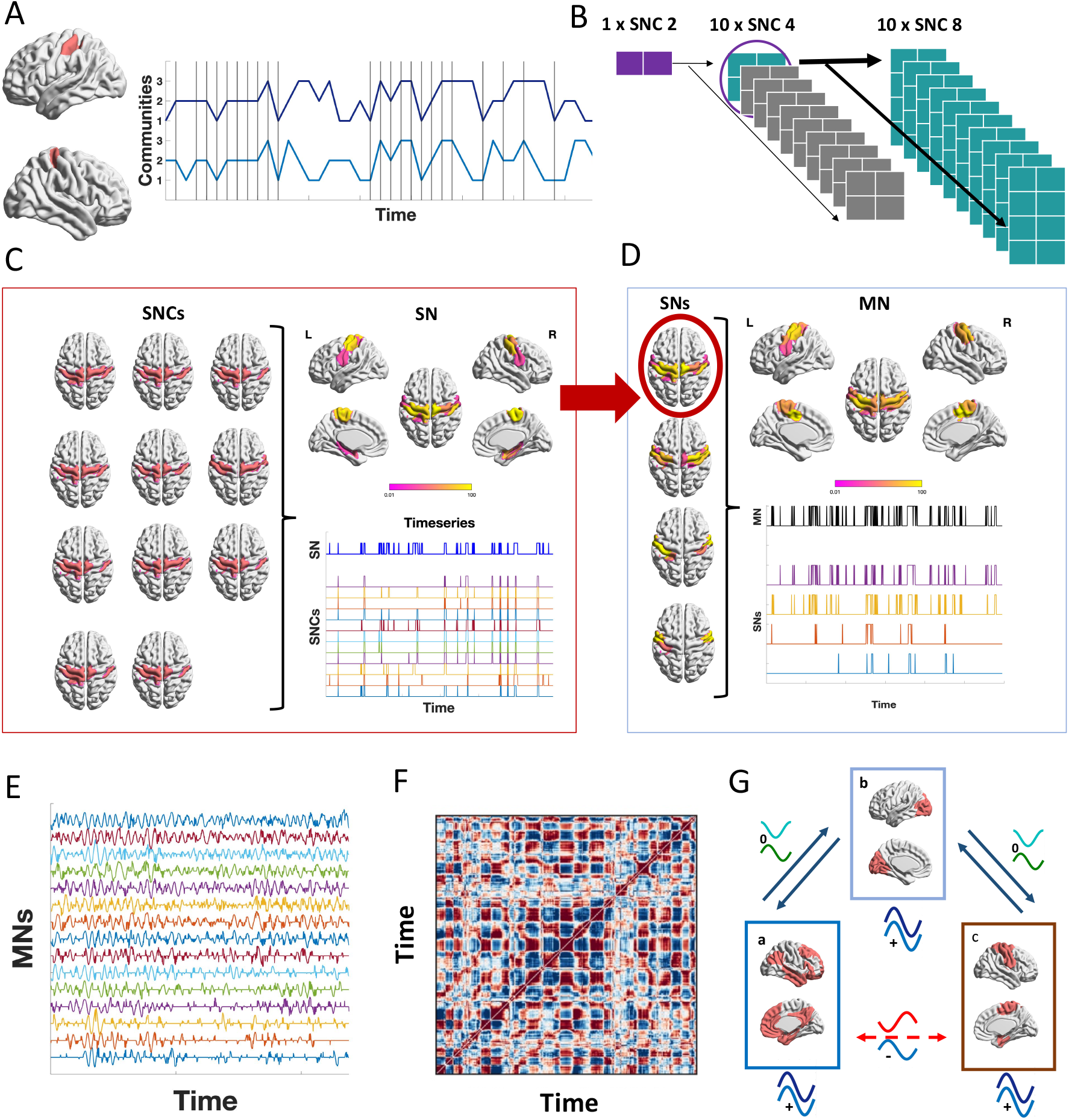
Schematic figure that summarizes the key steps for the proposed model. A) The calculation of the parameter q_int_ is illustrated for the case of a pair of brain areas in the left and right motor cortex. Vertical lines denote the time points when the areas are assigned to the same community. The parameter q_int_ for the two areas is then computed as the number of times they are assigned to the same community divided by the total number of time points in the fMRI time series. B) Based on the highest values of q_int_, seeds pairs are selected for each area and expanded in an iterative search to identify SNCs (subnetwork components) with an increasingly larger size. At each iteration (i.e. increase in size of SNCs) the ten constellations of brain areas with the highest q_int_ are selected and included as seeds in the next iteration. The search is halted when the SNCs reaches a size of eight areas. C) An example of how 11 SNCs together form a SN (subnetwork) located in the motor cortex. D) An example of how SNs are clustered together to represent a MN (meta-network). E) Schematic representation of state vectors (the amplitude values of the MNs) that are the combined representation of time resolved amplitude fluctuations in all MNs (the same type of vector can be constructed from SNs). F) Recurrence plot representing the correlation between all state vectors. G) Schematic illustration of integration (positive phase coherence) and segregation (orthogonal or negative phase coherence) between SNs in different communities. While the SNs in each community a, b and c are integrated (internal positive phase coherence) they are segregated vis-à-vis each other. SNs in a and c are the most segregated by virtue of their negative phase coherence. In order for these networks to integrate they would have to make a transition through intermediary states of orthogonal segregation (b) illustrated by dark blue double lines. The red dotted line between a and c signify that SNs cannot make a direct transition between them.

### 2.4 IPSA and EMD of resting-state fMRI time series

To date there is no golden standard for the analysis of time-resolved functional connectivity for reviews, see Hutchison et al., 2013a; Keilholz et al., 2017; Preti et al., 2017. Here we chose to employ the Instantaneous phase synchrony analysis (IPSA), which has recently gained increased interest in the literature (Cabral et al., 2017; Glerean et al., 2012; Pedersen et al., 2018; Ponce-Alvarez et al., 2015). The instantaneous phase of a signal can be obtained after applying the Hilbert transform on the real signal *s* by converting to its analytical expression in polar coordinates:

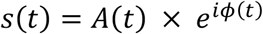

where *t* is time, *A* is the instantaneous amplitude (envelope) and the instantaneous phase. However, in order for the instantaneous phase to be interpretable, the signal should not contain riding waves (Huang et al., 1998; Niazy et al., 2011). One way to accomplish this is to apply a narrow bandpass filter were the frequency interval of interest must be specified. An alternative is to use the Empirical Mode Decomposition (EMD), the core innovation of the Hilbert-Huang transformation (Huang et al., 1998). The EMD is a sifting algorithm that decomposes a signal into a finite number of oscillatory components called intrinsic mode functions (IMFs). No a priori assumptions about frequency intervals, stationarity or linearity have to be made. The IMFs are designed to have approximately locally symmetrical waveforms (local mean of upper and lower envelopes equals zero) without riding waves which makes them well suited for subsequent Hilbert transformation for extraction of instantaneous phase and frequency information. The EMD-framework has previously been used on BOLD data (Niazy et al., 2011). IMFs have also been shown to have a high similarity with band-pass filtered signals at corresponding frequency intervals when analyzed for specific RSNs (Niazy et al., 2011). Here, the EMD-algorithm as implemented in MATLAB was used. The first IMF always captures the highest frequencies and subsequent IMFs encapsulates lower frequencies with some overlap. The EMD-decomposition of the BOLD fMRI time series yielded a minimum of six IMFs for all brain areas. The power spectra for all six IMFs are shown in Suppl. Figure S1. An example of IMF BOLD signal intensity time-series is provided in Suppl. Figure S2. The third IMF that had 99% of its power in the range of (0.003,0.1Hz) was chosen for further analysis. Of note, it was also the dominating IMF within the “classical” BOLD band (0.01-0.1Hz) (Biswal et al., 1995) accounting for 40% of the total power. The IMF time series were Hilbert transformed to yield instantaneous phase information. Due to end artefacts arising from the Hilbert transformation, five volumes at the beginning and end of the time series were removed resulting in 1190 time points for further analysis (Huang et al., 1998). At each time point the phase relationships between all brain areas were calculated as the cosine of the phase difference. This resulted in the instantaneous phase coherence matrix dPC (dynamic phase coherence) (Cabral et al., 2017; Glerean et al., 2012) where the relationship between brain areas *n* and *m* was calculated according to:

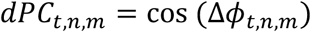

where Δ*ϕ* is the instantaneous difference in phase between area *n* and *m* at time *t*. Importantly, dPC captures the whole spectrum of positive, negative and orthogonal relationships between brain areas. Positive values hence reflect positive phase coherence, where 1 represents perfect in-phase alignment. Similarly, negative values correspond to negative phase coherence where -1 is perfect anti-phase alignment. A value of 0 reflects a complete decoupling, i.e. orthogonality. The distribution of dPC values for a single subject is given in Suppl. Figure S3. This measure is analogous to the correlation measure that also spans the [-1,1] range. Therefore, the IPSA framework has the power to capture changing connectivity through cycles of activation and deactivation of the BOLD-signal.

### 2.5 Defining subnetwork components (SNCs) and subnetworks (SNs)

The first step to construct the second smallest unit of subnetworks, the SNCs, was to find a measure that adequately captured the strongest time-resolved phase coherence relationships between brain areas across time. Our proposed novel measure (i.e. q_int_) to achieve this aim is described in section 2.5.1. Based on the definition of the q_int_ parameter, we define integration, segregation and disintegration between brain areas in section 2.5.2. Section 2.5.3 briefly outlines the most important consideration for choosing the initial brain area seeds. For details on the full algorithm that identifies SNCs, please see Supplementary Text. Section 2.5.4 provides a description of how SNs were constructed from SNCs.

#### 2.5.1 Assignment of brain areas into communities and estimation of the relative frequency of assignment to the same community (qint)

To identify substructures in complex data, community detection algorithms are commonly used (Newman, 2006; Rubinov and Sporns, 2011; Sporns and Betzel, 2016). In this context, with the time-resolved dPC-matrices as a foundation, we reasoned that a frequent (across time) assignment of brain areas to the same community would constitute a measure of a strong underlying relationship in terms of phase coherence. The parameter q_int_ was therefore defined to be the relative frequency (computed across all time points of the resting-state fMRI data acquisition) for which brain areas were assigned to the same community. Each phase-coherence matrix was hence divided into communities. Since phase coherence relationships can be both positive and negative (dPC range [-1,1]), the Louvain modularity algorithm was chosen because it accepts adjacency matrices with both positive and negative edges (Blondel et al., 2008; Rubinov and Sporns, 2010). The distribution of dPCs in the present dataset had two bimodal peaks close to -1 and 1 respectively (and a third one close to 0, see also Suppl. Figure S3). This distribution of the dPC values mandated an equal weighting of positive and negative dPC relationships. For this reason the “negative symmetric” option was used as implemented in the Louvain algorithm in the Brain Connectivity toolbox (Rubinov and Sporns, 2010). The information captured in the key parameter q_int_ can be illustrated by a simple example using two brain areas sampled at five points in time. Since the Louvain community algorithm, when applied to dPC data, divided all areas into 2 or 3 communities, the community vectors for the two areas in our example could be set to [1 1 2 1 3] and [1 1 1 3 3], respectively. In this example, q_int_ is equal to 0.6 since the two areas are assigned to the same community in three out of five points in time. More generally, for community vectors X and Y, we write

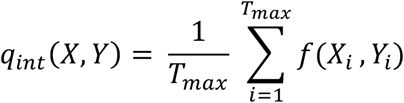

where *f*(*X_i_*, *Y_i_*) = 1 *if X_i_* = *Y_i_* and *f*(*X_i_*, *Y_i_*) = 0 *if X_i_* ≠ *Y_i_*. *T_max_* is the total number of time points in the BOLD signal intensity time-series.

#### 2.5.2 Defining integration, segregation & disintegration

Since the assignment of brain areas to communities described above was based on the time-resolved phase coherence matrices, a reasonable assumption would be that areas assigned to the same community at any given time point would, in general, have a positive phase relationship vis-à-vis each other. In the same way, areas assigned to different communities at the same time point would be either approximately orthogonal or have a negative phase coherence relationship. Based on these assumptions, *integration* was defined to be the simultaneous assignment of brain areas to the same community. Similarly, segregation was defined as the relationship that described areas assigned to different communities. Segregated areas were therefore assumed to have either orthogonal or a negative phase coherence relationship. For all levels in our hierarchical model (pairwise areas, subnetwork components (SNCs), subnetworks (SNs) and meta-networks (MNs)), the assumptions that integration equaled positive phase coherence and segregation equaled orthogonality or negative phase coherence were tested and validated (this is described in detail in section 2.7). It is important to note that with these definitions, integration and segregation are not mutually exclusive events in the temporal domain. Two ensembles can be integrated at the same time but in separate communities and therefore segregated relative to each other. This means that integration and segregation can take place simultaneously. In addition, the term *disintegration* was introduced to describe points in time when a SNC (second smallest network building block, i.e. level 2) was not integrated (the same principle holds for pairs of areas, i.e. level 1). At those time points, the areas that constitute the SNC will be assigned to different communities due to, on average, orthogonal or negative phase coherence between the areas. If all of the SNCs that together form a SN were to be disintegrated it would mean that the corresponding SN also would be disintegrated at that point in time. The definitions of integration, segregation and disintegration are schematically depicted in Suppl. Figure S4.

#### 2.5.3 Definition and identification of subnetwork components (SNCs)

The next step was to define subnetwork components (SNCs) based on the information of the frequency of assignment of brain areas to the same community (q_int_). The aim was to represent both modular and flexible tendencies among areas and to find building blocks that together captured the spatiotemporal variability at the next level of granularity (i.e. subnetworks, SNs). By modularity we here refer to a preference to repeatedly synchronize with the same set of areas while flexibility represents the tendency to synchronize with different sets of areas at different time points. At this level of spatial granularity, modularity would correspond to SNCs with high values of the q_int_ parameter, while more flexible yet potentially functionally specific relationships would be represented by SNCs with a lower value of q_int_. Since areas vary in their tendencies towards modularity or flexibility, it was important that the strongest functional relationship from the point of view *of each area* were represented from the start in order for the stronger modular tendencies not to override the weaker tendency of flexibility. Therefore, SNCs were derived in an iterative, stepwise fashion starting with pairs of areas such that all areas in the parcellation scheme were represented. For each area, only the pair with highest q_int_ value was included as an initial seed (see also Suppl. Figure S5). The size of the SNCs were grown in steps of two until the final size of eight areas was reached (Figure 1B). Each SNC had a subject specific time-course of community assignment that was used to calculate the relative frequency (q_int_). For details on the algorithm to construct SNCs and our motivation for parameter choices, see Supplementary Text. See also Suppl. Figure S6 that shows the distribution of q_int_ values for randomly constructed SNCs of different sizes together with SNCs derived by our algorithm.

#### 2.5.4 Construction of subnetworks (SNs)

Previous work has shown that spatiotemporal variability in network configuration is, together with modularity, an inherent aspect of brain organization. In this step, the aim was to capture both principles in spatiotemporally flexible subnetworks (SN) where the configuration of SNs were allowed to vary over time. To achieve this, the SNCs derived in the previous step were used as flexible and partly overlapping building blocks that were combined into SNs. In the same way as for SNCs, each SN had a subject specific time-course of community assignment and time-resolved spatial configuration that resulted from combining all the time courses and areas of the SNCs into a common time-resolved representation of the SN. Thus, a SN was considered integrated if at least one of its constituting SNCs were integrated at that particular point in time (see also Suppl. Figure S7). That the spatial configuration of SNs in terms of brain areas that are included varies over time implies that some areas will often or always be part of a specific subnetwork while other areas will more rarely make a contribution. This type of probabilistic representation could in theory allow all areas with some small probability to be part of all subnetworks or ultimately create one large undifferentiated network. Therefore, in order to penalize the undesired scenario where all areas would be included in all SNs, a criterion of internal coherence of SNs was used to guide the combination of SNCs into SNs. Internal coherence meant that none of the SNCs that together formed an SN could at any time point be segregated vis-á-vis each other (integrated simultaneously in different communities). An example of how SNCs were combined to yield a SN is given in Figure 1C. For further details on the construction of SNs see Supplementary Text.

### 2.6 Definition of Meta-networks (MNs)

The SNs derived in the previous step resulted in several spatially overlapping representations. We therefore sought to clarify the relationships between the SNs in terms of integration and segregation. The aim was, if possible, to aggregate SNs into larger meta-networks (MNs) that together would reflect the main patterns of integration and segregation in the brain. Multiple spatiotemporally interleaved SNs would together constitute an MN if the MN most of the time could adequately summarize the collective behavior of the SNs in a single, integrated unit. To establish suitable criteria to achieve this aim, we first calculated the degree of integration (i.e. frequency of assignment to the same community) between all pairs of SNs relative to the total number of time points of temporal overlap (i.e. when SNs were integrated simultaneously) between all SNs. This measure of integration, denoted q_tov_, only differed from q_int_ in that it was calculated relative to total time points of overlap instead of all time points in the scan. The calculation of q_tov_ was first done separately for each subject before the mean was taken across all subjects. Then, the pairwise frequency of segregation between SNs relative to temporal overlap, i.e. the complement to integration 1-q_tov_, was used as a distance measure for hierarchical clustering of SNs into MNs. For this purpose, the *linkage* and *clust* algorithms as implemented in MATLAB were utilized. All solutions of clustering in terms of number of clusters from two up to the total number of SNs minus one were evaluated. We defined two criteria for selecting a suitable number of clusters to work with. The first criteria focused on the mean degree of integration between all SNs that were clustered together into a MN. Here, we opted to set the threshold to 95 %, meaning that all SNs included in a MN were, on average, integrated in the same community at least 95 percent of the time of temporal overlap. Second, we used the minimum degree of integration between SNs (threshold set to 80 percent of temporal overlap). An example of how SNs were combined to form a meta-network (MN) is given in Figure 1D. Further details regarding the thresholds for forming MNs are given in the Supplementary Text.

### 2.7 Phase coherence

In this section we describe the steps taken to test the assumption we made previously (see section 2.5.2) that assignment to the same community (i.e. when a community algorithm is applied on a phase coherence matrix) corresponded to a positive phase coherence relationship. This was tested at three levels of network granularity (i.e. SNCs, SNs, and MNs). In the first section (2.7.1) that concerns the phase coherence within SNCs, we also investigated our assumption that meta-stability is the foundation for dynamic alterations in brain connectivity. This assumption was tested by calculating phase coherence for SNCs prior to and during the time points of integration (i.e. assignment to the same community). Additionally, we also tested the assumption that the phase coherence within a community is not equally strong for all areas. For this purpose, we randomly assigned brain areas to SNCs. Finally, in this section we also calculated the average phase coherence across all SNCs at time points of integration. In the second section (2.7.2), we evaluate the relationship between community assignment and phase coherence at the coarser spatial scales of subnetworks (SNs) and Meta-Networks (MNs).

#### 2.7.1 Assessment of phase coherence within subnetwork components (SNCs)

Within a meta-stable framework, a SNC would transiently quasi-synchronize where the duration of synchronization would depend on the strength of the phase coupling (dPC) (Tognoli and Kelso, 2014). Therefore, assignments of brain areas to the same community should cause a transient, but significant rise in dPC within the SNCs. For this purpose, a sample of 500 randomly selected SNCs (from the top two thirds of the distribution) was used. First, the mean dPC was computed between the brain areas constituting each of the 500 samples at time points prior to and during assignment to the same community. Time points were defined relative to the moment of integration t_0_ such that time points before t_0_ had negative values (corresponding to disintegration) and time points at t_0_ and after had positive values (corresponding to integration). This calculation was done separately for each relative time point *t*, where *t* = -18, .. ,17. The phase values for each time point were averaged across instances of integration, subjects and SNCs. In more detail, if *Y* denotes the ensemble of all SNCs, *M* is the number of brain areas in the SNC, *S* the set of all subjects, *U* is the total number of instances of integration in the *y*:th SNC (*y* ∈ *Y*) in subject *s* (*s* ∈ *S*), then for each relative time point *t*, the mean phase coherence within SNCs, 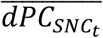 can be calculated as:

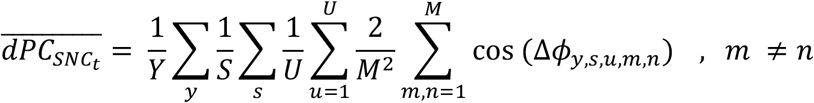

where Δ*ϕ_m,n_* is the instantaneous phase angle between area *m* and *n*. Moreover, we tested for a potential positive relationship between the duration of integration and peak strength in dPC. Finally, the strength of the phase coherence of the SNCs was compared to that of 443 randomly combined components of the same size (size of SNCs = 8 brain areas). Additionally, we also calculated the average phase coherence across all SNCs at time points of integration.

#### 2.7.2 Assessment of phase coherence within and between subnetworks (SNs) and meta-networks (MNs)

Next, we tested the assumption that integration implied positive phase coherence also held at the level of SNs and MNs. In an analogous manner to the case of SNCs described above, we first computed the mean phase-coherence (dPC) within each SN at each time point of integration. This was achieved by first calculating the mean phase of the areas for each SNC that represented the SN at each time point. Subsequently, the cosine of the difference in phase-angle between all pairs of SNCs was calculated and its mean was computed. More formally, if *Z* denotes the ensemble of all the SNs and *G* is the number of SNCs in *z* (*z* ∈ *Z*) that are integrated at time *t,* then the mean phase-coherence within SNs, 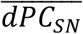 can be calculated as:

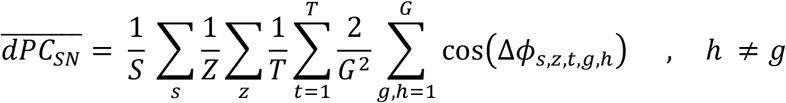

where *H* = *G* and *T* is the total number of time points of integration for each SN.

In a subsequent step, the similar procedure was repeated for calculating phase coherence between SNs integrated in the same community. Notably, the computation of differences in phase between SNs was done separately for SNs that belonged to the same MN and between SNs that belonged to different MNs. The mean phase-angle for each SN at each time point was calculated based on the mean of the phase-angle of each SNC included in the SN. More formally, let *C* be the number of unique communities at time *t* and z (*z* ∈ *Z*) the set *of* SNs integrated at time *t* in community c (*c* ∈ *C*). Moreover, let *L* be the number of different MNs in c and let z_s_ (*z*_2_ ∈ *Z*) denote the subsets of *Z* that consists of SNs from the same MN. Also, let z_d_ (*z_d_* ∈ *Z*) be the subsets of *Z* containing combinations of SNs from different MNs (from the perspective of each MN). In the first case, i.e. for SNs belonging to the same MN, the average phase coherence when integrated in same community can be calculated as:

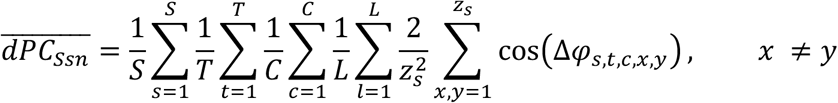

Similarly, average phase coherence between SNs from different MNs can be calculated according to:

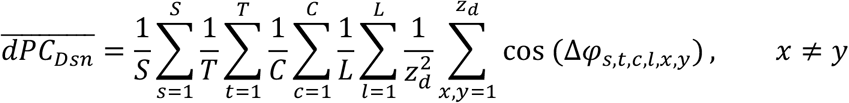

Our assumption that assignment to different communities at the level of SNs was associated with negative phase coherence or orthogonality was subsequently tested in a similar way.

### 2.8 Change in global state

Functional activation in the brain is intrinsically highly organized and also in constant change. The change in brain activity is bounded by its core organization such that some patterns of co-activation occurs frequently, i.e. the RSNs, while others are vastly unlikely to appear. The perhaps simplest way to look at the main patterns of intrinsic co-activation is by pairwise correlation of brain area time-series. Thus, the information in the 2D space (N) x time (T) matrix, where N is the number of areas in the parcellation and T is the number of time points, is summarized in a spatial N x N matrix. In a similar way, one can appreciate how the global brain configuration, in terms of time-resolved BOLD signal amplitudes, changes across time. Instead of doing pairwise correlation of brain areas time-series, the correlation is done over the temporal dimension resulting in a T x T matrix (i.e. a recurrence matrix). The vector at each time point contains the time-resolved amplitude for each of the N areas. After demeaning, positive amplitudes can be interpreted as relative activation and negative amplitudes as relative deactivation. Thus, by combining the joint information of activations and deactivations of all areas at time *t*, we achieve a simple way to represent the time-resolved state of intrinsic organization of the whole brain at any given point in time. To be clear, by “state” we only refer to the configuration of activation and deactivation of all the areas in the parcellation considered simultaneously at one point in time.

The SNs and MNs derived in previous steps were, in essence, coarse-grained higher order representations of the phase information contained in the time-resolved co-activations and co-deactivations between individual brain areas. Importantly, the networks defined at the coarser spatial resolutions (SNs and MNs) would only have sufficient contrast if they adequately captured the integration and segregation at the lower levels of spatial granulation (areas and SNCs). We therefore wanted to test if our coarse-graining strategy adequately captured the raw, time-resolved patterns of whole-brain recurrences in amplitude fluctuations observed in the BOLD IMF3 time-series. This test was done by constructing time-resolved state vectors based on SNs and MNs and compare their patterns of recurrence to that of the state vectors based on individual brain areas as described above. First, each BOLD IMF3 time-series was demeaned and normalized to the absolute value of its maximum amplitude (positive or negative). The purpose of this procedure was to prevent that areas with large amplitude fluctuations contribute disproportionally to the average amplitude of the SNs and MNs. The time-resolved vectors were computed as the mean amplitude (BOLD IMF3 signal intensity time-courses) of each of the SNs and MNs. Following the reasoning outlined above, global brain state-vectors were subsequently constructed, yielding a T x T recurrence matrix for each subject at each level of spatial granularity (brain areas, SNs and MNs). For each subject, the degree of similarity between all three types of recurrence matrices (brain areas, SNs, and MNs) was evaluated using the Mantel test (Spearman’s rho) with 1000 permutations. Subsequently, the mean was the taken across subjects.

### 2.9 Assessment of flexibility and modularity at the levels of individual brain areas

A major challenge for methods that tries to capture core aspects of functional connectivity in the brain is the potential heterogeneity of connectivity profiles in terms of divergent tendencies towards either flexibility or modularity, or both, among brain areas. Most methods are likely to be more or less skewed towards modularity rather than flexibility since the former is easier to capture. Here, we applied our method to estimate heterogeneity in connectivity (based on properties of SNCs) at the level of individual brain areas. The degree of heterogeneity for individual brain areas in terms of their relative flexibility, modularity or both was estimated by computing two parameters. First, we calculated the degree of temporal representation (p_int_) which was defined as the relative frequency of participation of an area in any SNC across the scan time (i.e. the relative frequency with which the area was represented across all time points by the model). The theoretical range for p_int_ is 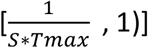 where T_max_ is the total number of time points and S is the number of subjects. A p_int_ value of 1 implies that an area is represented by the model at all points in time in all subjects. In contrast, the lowest theoretically possible value for p_int_,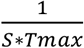, reflects the situation where a brain area is part of only one SNC which is integrated only once in a single subject. Second, we computed the spatial diversity for each brain area, p_div_ which is defined as the proportion of the total number of areas that each particular area shares an SNC with. The theoretical range for p_div_ is 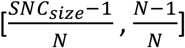 where N is the number of areas in the parcellation and SNC_size_ is the number of areas in each SNC (SNC_size_ = 8).

## 3 Results

### 3.1 Subnetwork components - SNCs

In a preparatory step to identify SNCs, the pairwise relationships between all brain areas were analyzed in terms of the relative frequency with which they were assigned to the same community (q_int_). These results are presented in the Supplementary Text and Figures S8. At each step of the iterative process to find the most frequent and representative eight area SNCs, we found a substantial amount of convergence across the hierarchical levels of iterations such that a portion of SNCs of size n (n= 2, 4, 6 and 8 brain areas) merged into the same SNC at the next level (n+2). Convergence at level n was defined as one minus the proportion between the unique number of SNCs and all SNCs resulting from the seeds at level n - 2. This implies that SNCs that were individually specific and recognizable at a lower complexity level (i.e. smaller SNCs) merged together at the next level of complexity (spatially larger SNCs). The degree of merging increased substantially as the size parameter n increased: From n = 2 to n = 4 by 17%, from n = 4 to n = 6 by 52%, and from n = 6 to n = 8 by 66%. As a result, the iterative procedure yielded 31976 unique SNCs of size n = 8. This number of SNCs corresponded to only 14% of the total number of SNCs that would have been formed if no convergence/merging were to be present in the data. The observed degree of convergence reflected the strong interconnectedness of the functional topology, in so far that the same SNCs were formed from multiple starting seed areas. In addition, this finding shows that it was sufficient to use only the top ten constellations of areas in each iteration to obtain a sparse representation of the strongest relationships between brain areas in the form of SNCs. However, for completeness, we also tested the effects on our results by increasing the expansion factor K such that more SNCs were sampled (see Supplementary Text, Figures S22 - S28 and Tables S1 - S4 for detailed comparison between the results for K = 10, 20 and 40 as well as replacing the EMD step with a narrow band-pass filter for K = 10, Supplementary Text and Figures S29 - S34). For the sake of simplicity, in the following we present only the results for K = 10. Across subjects, individual SNCs remained integrated during short time periods (q_int_ values ranged in a low regime 0.01 to 0.12 with a median of 0.06, see also Suppl. Figure S22 and Suppl. Table S1). Also, there was as strong positive relationship between q_int_ and mean phase coherence (dPC) within SNCs on a group level (r = 0.78).

### 3.2 Subnetworks - SNs

In our hierarchical model, the SNs were per definition the largest network units that were always guaranteed to be integrated into the same community. Therefore, the main condition for grouping SNCs into SNs was that all the SNCs in an SN at all instances in time of simultaneous integration should be assigned to the same community. The minimum degree of spatial overlap between the reference SNC and the other SNCs (see also Suppl. Figure S7) in the SN that maintained this condition was identified heuristically and found to be five out of eight areas. This rule resulted in a total number of 240 SNs, where the number of SNCs within SNs ranged between 1 and 2527 and each SN contained from 8 up to 63 individual brain areas (median = 20, IQR = 14). Next, the brain areas in the SNs were weighted according to their relative frequency of occurrence in the SNs. A summary of the topological maps for all SNs is given in Supplementary Video S1.

Due to the shape of the underlying distribution of the dPC values, the Louvain algorithm consistently partitioned the dPC matrix into three communities for the vast majority of time points (89%). The remaining time points (11%) were divided into two communities. At no time was there only one community. This finding highlights the notion that a certain degree of segregation was always present in the data. At any given point in time, on average 19% of all SNs (SD = 3%) were found to be integrated. For most time points, there were 2 or 3 communities that contained integrated SNs but sometimes only one (mean number of communities with integrated SNs = 2.4, SD = 0.1). The SNs integrated in different communities had (mostly) an orthogonal or negative phase-coherence (anti-phase) relationship to each other (see Figure 1G & Suppl. Figure S9). However, when the size of the sampling from the pool of SNCs was increased (setting the expanding factor K = 40, see Supplementary Text), there were always SNs integrated in at least two different communities. This result suggests that some degree of segregation is always present as a core dynamical organization principle in the brain. That multiple and often spatially overlapping SNs were simultaneously integrated in the same community highlights the interleaved relationship between subnetworks. The spatially interwoven nature of the SNs was also apparent from the topological maps given in Supplementary Video S1. On average, across subjects, 46% (SD = 5%) of the total number of brain areas were represented as part of an SN at any given time point.

As previously pointed out in the framework of coordination dynamics, strong coupling results in longer duration of integration or dwell time (Tognoli and Kelso, 2014). Analogously, this relation was also present in the SNs, where SNs with stronger phase coherence (in terms of q_int_ that in turn is related to coupling strength) had longer durations of integration (Suppl. Figure S10). However, the duration of integration also had a strong relationship to total number of SNCs in the SNs (Spearman’s correlation r = 0.81). Taken together, these observations explain to a large part the relationship between q_int_ and duration of integration at the level of SNs.

### 3.3 Meta-Networks - MNs

The meta-networks (MNs) constituted an attempt to summarize the relationships between the SNs in terms of main tendencies towards either integration or segregation in the time domain. This attempt was deemed important since several of the SNs derived in the previous step showed a large degree of topological overlap. The clustering of SNs into MNs was therefore based on the principle of maximizing the relative frequency of integration between SNs that were to be grouped into the same MN. The threshold for clustering SNs into MNs was set to assure that all SNs constituting an MN were integrated in the same community on average at least 95% of the time of temporal overlap. Additionally, we set the minimum degree of internal integration for any MN in the clustering to be at least 80% of the temporal overlap (see also Methods section 2.6 and Supplementary Figures S11 and S12). These two thresholds were chosen to assure that MNs could be regarded as integrated network units most of the time. Given these choices, the dynamics of the 240 SNs were summarized into 71 MNs (Supplementary Video S1 contains a visual presentation of all MNs as well as the SNs that each MN is made of, sorted with respect to highest q_int_ of the MNs). While most MNs were combinations of several SNs, some of the less frequent MNs contained only one SN. The ten most frequent MNs included 65% of the total number of SNs and their topological maps are shown in Figure 2. In general, the spatial topology of the MNs showed a high degree of similarity with the core RSNs: either as part of a RSN, or as a combination of two or more RSNs. Each MN was therefore named according the RSNs (as defined by the parcellation, see Methods 2.2) that contributed most weight in terms of frequency of participating areas. If other RSNs contributed to more than a third of the weight of the dominating RSN, the MN was considered to be a combination of those RSNs and named accordingly. This naming strategy highlighted the fact that areas belonging to separate classical RSNs were in fact frequently integrated. The most salient example was perhaps the DMN and FPN that contributed significantly to 12 separate MNs, of which three were amongst the top ten MNs most frequently integrated in the brain. The fact that parts of the DMN is intermittently strongly coupled to the FPN has been shown previously using iCAPs (Karahanoʇlu and Van De Ville, 2015). Strong coupling across borders between RSNs was observed already at the level of pairwise brain areas, where 26% of the initial brain area seed pairs were situated in separate RSNs. Notably, the DMN and FPN each contributed to half of all the seed brain regions that paired together areas located in different RSNs. Based on the results shown in Figure 2, we note that meta-networks MN7 - MN9 encompassed large parts of both the classical DMN and FPN resting-state networks, suggesting that they were in fact highly interleaved and integrated with each other. If we had opted to set the thresholds lower in the clustering step to form MNs, the three meta-networks would likely have merged into a single MN (potentially together with more SNs). While the degree of integration within MNs was optimized when the clustering thresholds were set high, it should be noted that some MNs were not always internally coherent and sometimes did split into different communities (i.e. segregate). Hence, a lower cluster threshold would result in fewer and internally less coherent MNs. The opposite effect would be seen with higher thresholds.

**Figure 2.**
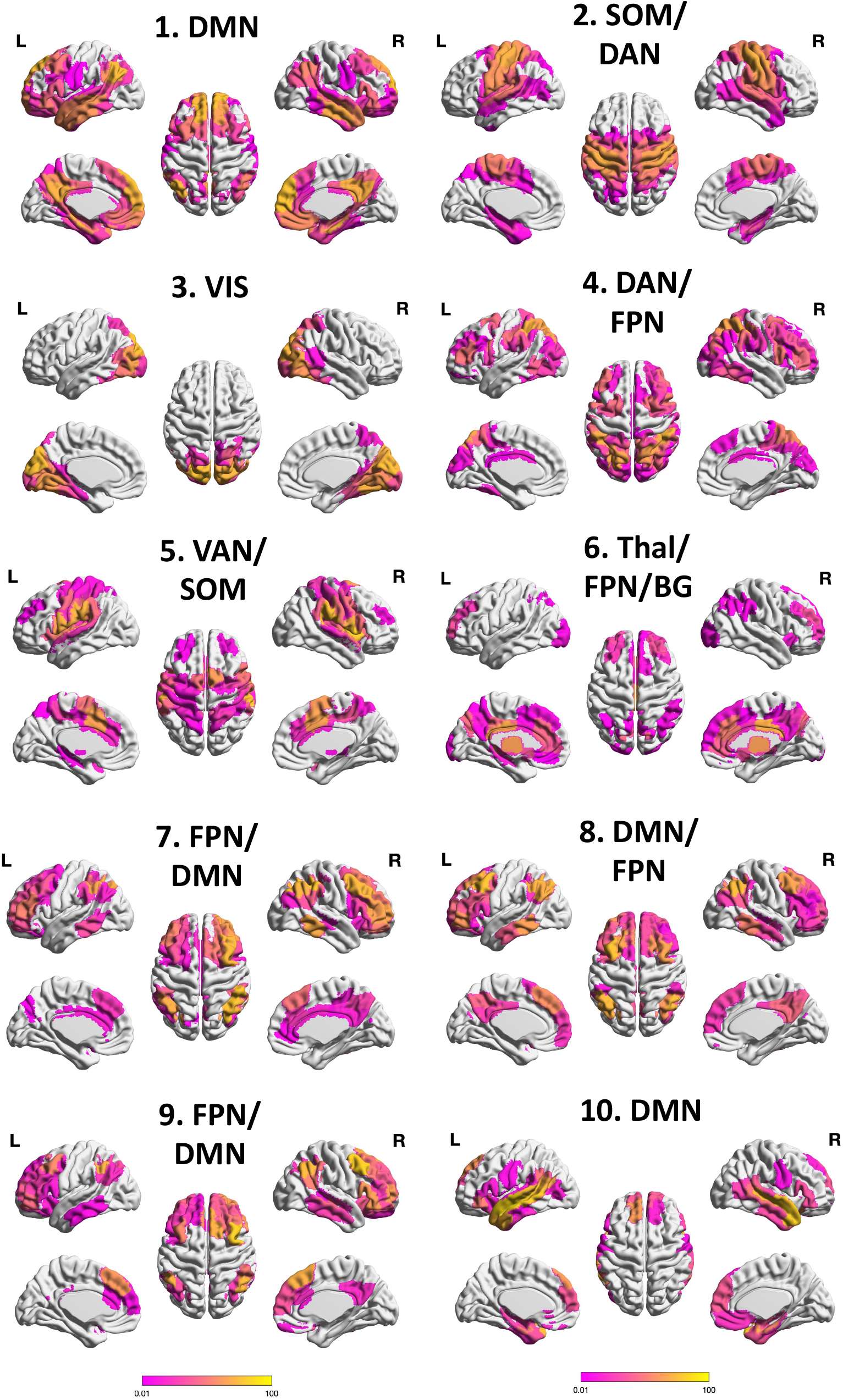
Topology of the top ten meta-networks (MNs) The areas in the maps were color coded according to the relative frequency of participation in the MN. Thus, most MNs had a core of very frequently participating areas (marked in yellow) and a wider periphery (marked in pink to purple) of less frequently participating areas. Bright yellow corresponds to q_int_ = 1, bright purple q_int_ = 0.01. Networks were given the name of the RSN that contributed most in terms of weighted areas to the MN. If other networks contributed more than a third of the weight of the dominating RSN, the MN was considered a combination of those RSNs and named accordingly. The RSNs were defined by the Shaefer 7 parcellation together with subcortical networks from the Brainnetome parcellation. The mean q_int_ values for the MNs across subjects were as follows **(1) DMN**: q_int_ = 0.94 (SD = 0.02). **(2) SOM/DAN**: q_int_ = 0.94 (SD = 0.03). **(3) VIS**: q_int_ = 0.83 (SD = 0.06). **(4) DAN/FPN**: q_int_ = 0.80 (SD = 0.05). **(5) VAN/SOM:** q_int_ = 0.75 (SD = 0.02). **(6) Thal/FPN/BG** q_int_= 0.72 (SD = 0.10). **(7) FPN/DMN** q_int_= 0.68 (SD = 0.06). **(8) DMN/FPN** q_int_ = 0.64 (SD = 0.06). **(9) FPN/DMN** q_int_ = 0.47 (SD = 0.07). **(10) DMN** q_int_ = 0.45 (SD = 0.1).

### 3.4 Phase coherence

#### 3.4.1 Phase coherence within subnetwork components (SNCs)

To test the hypothesis that assignment to the same community brought significant increase in phase coherence within SNCs, we computed the mean phase coherence (dPC) between the brain areas constituting each of the 500 sample SNCs at time points prior to and during assignment to the same community (Figure 3). Time points were defined relative to the closest subsequent time point of integration t_0_ (Figure 3C) such that time points before t_0_ had negative values (corresponding to disintegration) and time points at t_0_ or beyond had positive values (corresponding to integration). The dPC value for each time point was calculated separately where values close to t_0_ were based on many samples while values further away were based on less samples (see Methods 2.7.1 and Figure 3C and Suppl. Figure S13). Additionally, we computed the mean q_int_ in the same way for brain areas that were part of randomly generated SNCs of size eight. In our sample of SNCs, the mean q_int_ was more than 20 times larger (mean q_int_ = 0.07, SD = 0.01, range [0.04, 0.11), than for random components (mean q_int_ = 0.003, SD = 0.02, range [0.0006, 0.02]). For both SNCs and random components, duration of integration (i.e. assignment to the same community) was a function of q_int_ such that higher q_int_ values implied longer durations (Spearman’s r = 0.97)(see also Supplemenary Figure S14). As shown in Figure 3, the mean *dPC* within SNCs with long intervals between instances of integration started to increase on average eight seconds prior to the assignment to the same community. On average, at the time point of assignment to the same community (denoted by vertical bars at t = 0 in Figures 3A and 3B), *dPC* was equal to 0.72 (SD = 0.03) for SNCs and *dPC* = 0.64 (SD = 0.04) for randomly generated components. Importantly, SNCs had on average a significantly stronger (p = 0.001, effect size = 1.58) peak phase coherence (mean dPC = 0.81, SD = 0.02) than random components (mean dPC = 0.66, SD = 0.04) (Supplementary Figure S15). Overall, the degree of phase coherence was weaker and its duration was shorter in the sample of random components compared to SNCs. Coupling within SNCs in the disintegrated state (time points prior to integration) was also significantly stronger (i.e. during the time-period t = -13 to t = -1 in Figure 3A and B) (permutation t-test p < 0.001). We also calculated the mean phase coherence at time points of integration for all SNCs (n = 31976) which was found to be dPC = 0.75 (SD = 0.05). Taken together, these results support the hypothesis that meta-stability was governing the dynamics of SNC integration, segregation and disintegration. They also suggest that integration was a gradual process where some brain areas in the SNCs initiated the integration while others were recruited with a time lag. While the integration of the SNCs presumably played a major role in driving the global dynamics, in contrast, the relatively weak integration of most random components was likely mostly a biproduct of this process.

**Figure 3.**
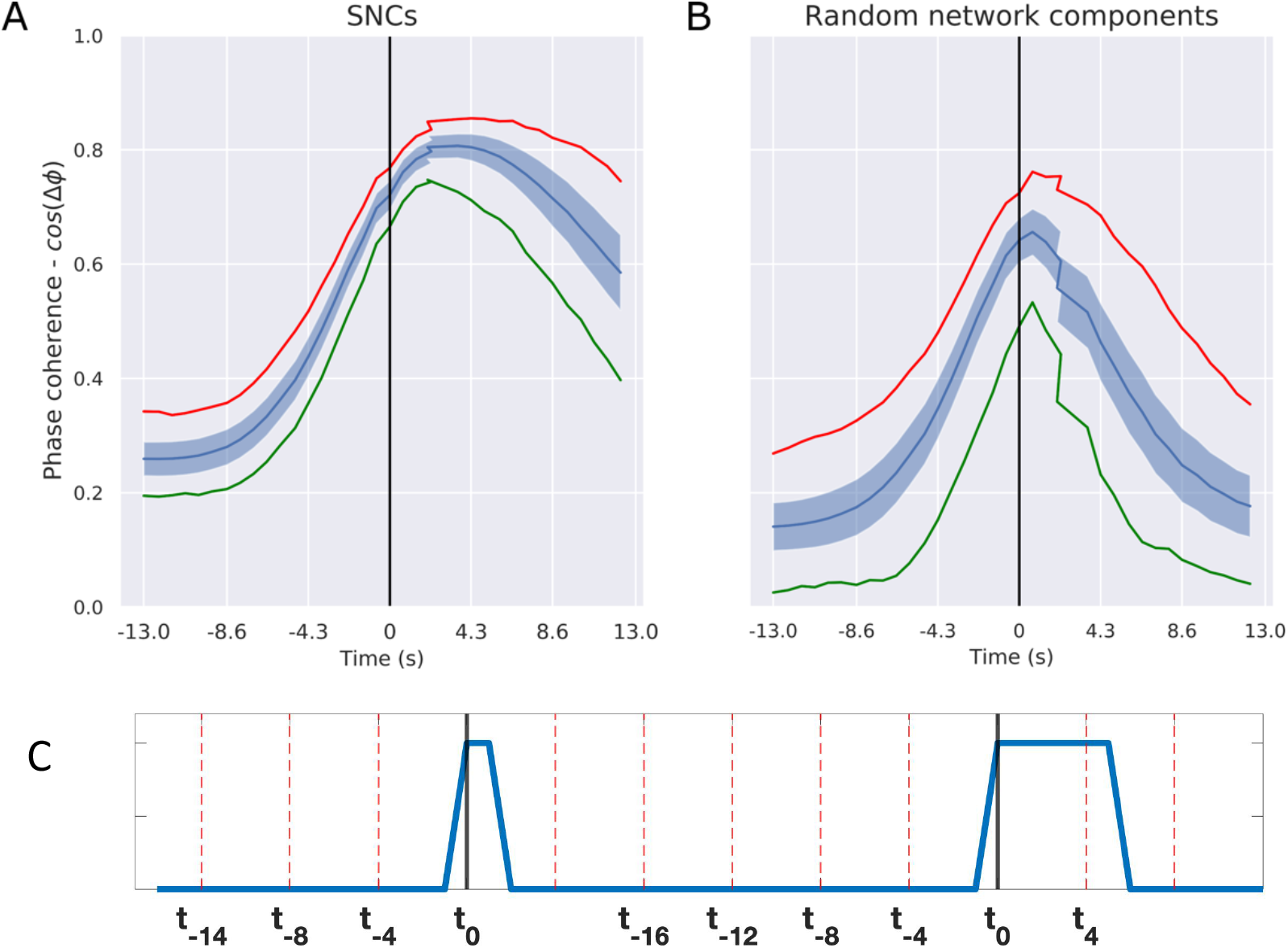
Degree of phase coherence (dPC) prior to and after the time point of integration for a sample of SNCs (A) and randomly generated components (B) The process of integration within SNCs was gradual and started before the point of integration at t = 0 (i.e. when all brain areas inside the SNC were residing in the same community). The schematic example shown in panel (C) shows the basis for calculating dPC values in a truncated and binarized SNC-time series (blue) in one subject. This schematic example contains two instances of integration separated in time by a period of disintegration. In each case, the first time point of integration is marked as t_o_ (corresponding to the vertical bar in positioned at t = 0 in panel A and B). The duration of integration for the first instance is two time units (t_0_ and t_1_) whereas the second epoch spans six time units (t_0_ and t_5_). The maximum duration of integration and disintegration included in the calculations were 18 time points (t_0_ to t_17_ for integration and t_-18_ to t_-1_ for disintegration). Here a time point corresponded to the time-resolution in the acquisition, i.e. 0.72 sec. This is reflected in the graphs of average phase-coherence per time point depicted in panels A and B but converted to seconds. Since many SNC had shorter duration of integration than 18 time points, the dPC values close to t_0_ were based on a larger sample than the time points reflecting long durations. The same was true for the intervals between integrations where both short and long intervals contributed to the negative time points just before t_0_. See also Suppl. Figure S13 for a sample density plot.

#### 3.4.2 Strength of integration within and between subnetworks (SNs)

First, we examined the average phase coherence within SNs, that is, between the SNCs constituting each SN at each point in time. As expected, the mean phase coherence within SNs was high, dPC = 0.96 (SD = 0.02). Next, the mean phase coherence between SNs residing in the same MN was compared to the phase coherence between SNs from different MNs that were integrated in the same community. SNs belonging to the same MN had a significantly stronger (p < 0.001, permutation test) mean phase coherence dPC = 0.94 (SD = 0.003), compared to SNs originating from different MNs dPC = 0.85 (SD = 0.007) (see Suppl. Figure S16). This finding suggests that the strength in phase coherence between SNs that were incorporated into the same MN was generally stronger compared to the degree of phase coherence between SNs belonging to different MNs at time points when they were integrated in the same community. The mean phase coherence between segregated SNs across subject (i.e. integrated simultaneously but in separate communities) was - 0.18 (SD = 0.06) (see also Suppl. Figure S9). Of note, the mean phase coherence within SNs (dPC = 0.96) and between SNs belonging to the same or different MNs (dPC = 0.94 and dPC = 0.85, respectively) at time points of integration compared to the mean phase coherence for SNCs (dPC = 0.75, see section 3.4.1) seems to suggest that the phase coherence both within and between SNs was stronger than within individual SNCs. However, a direct comparison is misleading since this apparent paradox stems from differences in how the calculations were performed. The calculation of phase coherence within SNCs was based on the mean phase of the SNCs while the calculations between SNs were based on the mean phase of the SNs. If instead all individual relationships between the areas within and between SNs had been considered (as it is done in the case of within SNCs phase coherence), the results would fall below that of the SNCs.

### 3.5. Recurrence in global connectivity

To test how well SNs and MNs together captured fluctuations in activity in the brain (the amplitude changes above and below mean of areas, independent of network), SN and MN derived time-resolved state vectors were correlated across time and compared to area-based vectors (as described in Methods section 2.8). The results are provided in Figure 4, which for increased clarity, shows a zoomed-in view (300 out of 1200 time points) in two different subjects as well as the mean across subjects (the recurrence matrices for the full experimental time are provided in Suppl. Figure S17). The state vector changes shown in the form of recurrence matrices in Figure 4 reveals how similar patterns of amplitude fluctuations were repeated with irregular periodicity across time. State vectors for all three levels (areas, SNs and MNs) revealed a quasi-cyclic pattern of recurrences that was more pronounced for SNs and MNs than for areas. Notably, time epochs of anti-correlations were present as well as epochs showing orthogonal relationships between state vectors. We further note that the SN and MN based recurrence matrices, which included periods of correlated, anti-correlated and orthogonal relations, spanned almost the full range of correlation values [-1,1]. The difference in terms of strength of the average correlation (absolute values) was significantly higher for SNs (middle column, r = 0.14) and MNs (right column, r = 0.21) compared to brain areas (left column, r = 0.11) whereas the correlation from time point to time point (diagonal) was stronger for the brain areas (due to the autocorrelation inherent in the time series). The similarities between the recurrence matrices at different levels of granularity (within subjects) was evaluated using a Mantel test (Spearman’s r). In all subjects, similarities between matrices were high (MN versus SN: r = 0.81 (SD = 0.03), MN versus brain areas: r = 0.57 (SD = 0.06), SN versus brain areas: r = 0.65 (SD = 0.05)). These results indicate that our hierarchical strategy adequately captured the segregation and integration between brain areas at different levels of granularity, which is a result that strengthens the validity of our method. If the network merging steps had been carried out over brain areas that did not have a positive phase coherence relationship, averaging amplitudes within SNs and MNs would have resulted in values close to zero. This in turn would have resulted in low contrast in terms of random fluctuations in the recurrence matrices unrelated to the baseline seen based on independent areas.

**Figure 4.**
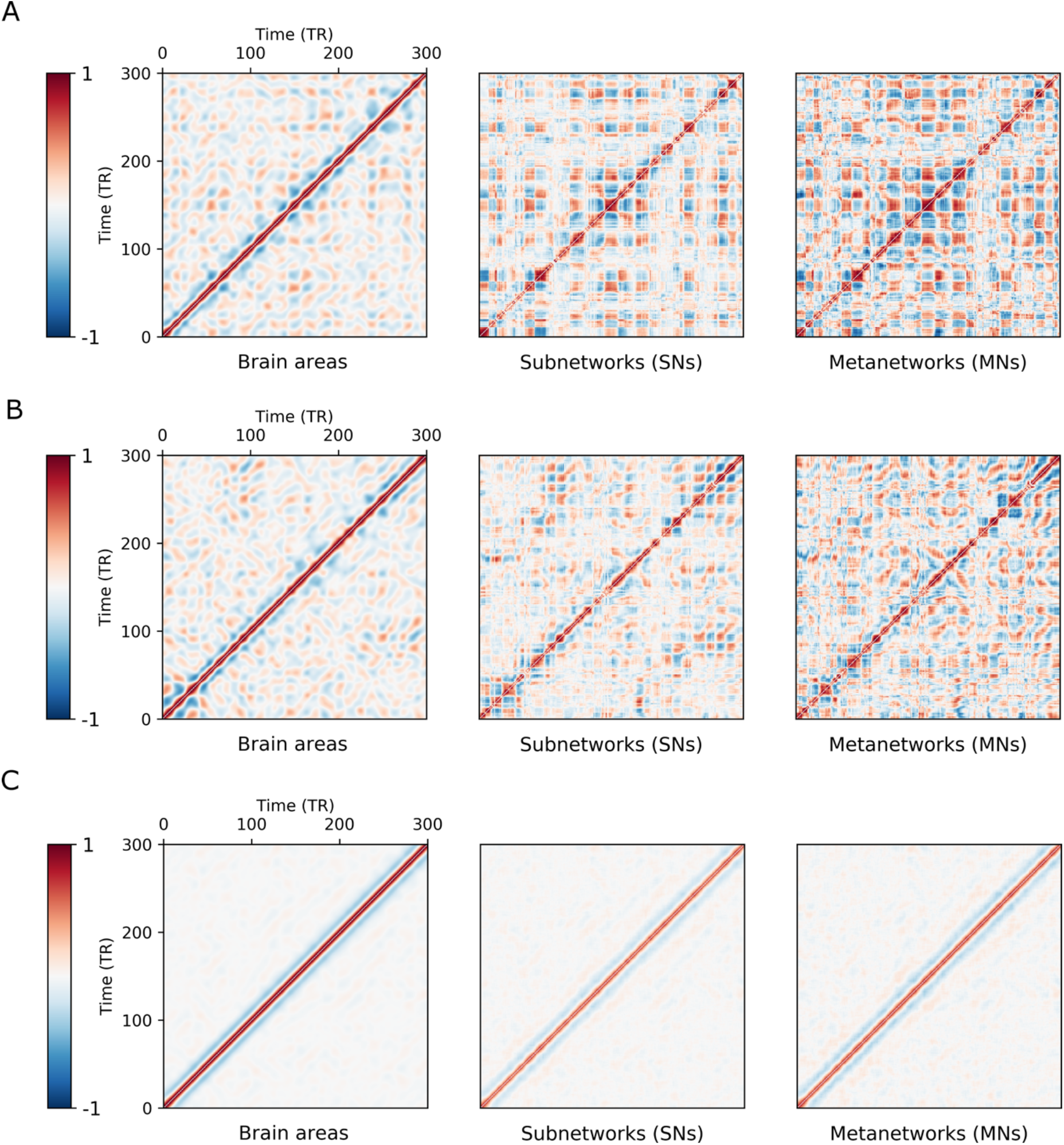
Quasi-cyclic recurrence of state vectors for areas time series, SNs and MNs (BOLD IMF) Example of recurrence plots of time-resolved state vectors across time for two different subjects (panels A and B) and the mean across all subjects (panel C). For increased clarity, each plot is a zoomed-in version focusing on the first 300 time points (out of a total of 1200 time points). For graphs that shows the full temporal range, see Suppl. Figure S17. For a version based on narrow-band pass filtered signals, see Suppl. Figure S34.

### 3.3. Flexible and modular tendencies at the levels of individual brain areas

To provide a comprehensive and informative picture of tendencies towards flexibility and/or modularity for individual brain areas, we plotted the temporal representation (p_int_) (x-axis) against spatial diversity (p_div_) (y-axis). The two-dimensional space spanned by these two parameters could broadly be divided into four categories as follows: (1) “flexible” brain areas in the lower left quadrant (below the mean p_int_ and below the mean p_div_), (2) “hyper-flexible” brain areas in upper left quadrant (below the mean p_int_ and above mean p_div_), (3) “modular/flexible” i.e. brain areas with mainly modular but also flexible tendencies in the upper right quadrant (above the mean p_int_ and above mean p_div_), and finally, (4) brain areas with exclusively “modular” tendencies in the lower right quadrant (above the mean p_int_ and below the mean p_div_). In this coarse categorization, the definition of flexibility/modularity was based on the logic that areas with low p_int_ (relative to the group mean) had a weak preference for forming specific constellations in time, reflected in a low q_int_ for all the SNCs in which they were part of. Rather, they would integrate briefly with a large set of areas forming SNCs with low values of q_int_, many of which would not be represented by the model. In contrast, areas with a very strong tendency towards modularity would, in principle, be marked by a high temporal representation (p_int_) and low value for spatial diversity (p_div_). However, for strongly modular networks consisting of many areas, p_div_ could be relatively large. For areas in the third category that were mainly modular but also flexible, the high p_int_ value would mostly be contributed by the SNCs representing modular tendencies. In contrast the high p_div_ in this group would to a large extent be a contribution from spatially diverse SNCs that has a much lower frequency than the main modular connections. “Hyper-flexibility” (4) would categorize areas that were represented only by spatially diverse SNCs that all had low q_int_.

The results for the estimated relative degree of spatial diversity (p_div_) of connectivity and frequency of temporal representation (p_int_) are shown in Figure 5 and were strongly correlated (r = 0.68, See also Suppl. Figures S18 & S19 for whole brain weighted spatial maps of p_int_ and p_div_). Brain areas that were frequently represented in time (high p_int_) also tended to be part of a larger number of spatially diverse SNCs (high p_div_). This group was referred to as “modular / flexible” (Figure 5, upper right quadrant). The high p_int_ values in this particular group were mostly a contribution from the high q_int_ values obtained from the SNCs that represented their main modular tendencies. However, this group of areas also had rather flexible connections occurring with lower frequency of integration, which contributed to the spatial diversity parameter p_div_. For example, core areas of the DMN as well as many areas residing in the DAN, FPN, VAN and SOM networks were assigned to the “modular/flexible” group. However, there was a significant dispersion along the both axes within most RSNs. On the “opposite” side of the two-dimensional space spanned by the p_div_ and p_int_ parameters was the “flexible” category (Figure 5, lower left quadrant). Interestingly, all brain areas located in the basal ganglia together with large parts of the thalamus and the limbic network fell into this category of strong flexibility. Additionally, a large portion of the visual network (VIS) was classified as relatively modular (lower right quadrant in Figure 5). In fact, these brain areas comprised a distinct band perpendicular to the cytoarchitectonic organization of V1 and V2 with bilaterally symmetrical p_int_ values (see also Suppl. Figure S20). Despite the fact that 32 per cent of the total number of all SNCs contained at least one subcortical or limbic area, these areas scored low both in temporal representation (p_int_) and spatial diversity (p_div_). It is notable that SNCs that contained any number of subcortical and/or limbic regions, either exclusively or in combination with cortical areas, had mean q_int_ values that on average were only half of what were obtained for the SNCs consisting of cortical areas only (q_int_ [subcortical and limbic areas] = 0.3 (SD = 0.01) versus q_int_ [cortical areas] = 0.07 (SD = 0.02)). Additionally, the mean phase coherence for this subgroup of SNCs was significantly lower than for SNCs that contained cortical areas only: dPC [subcortical and limbic] = 0.69 (SD = 0.05) vs dPC [cortical only] = 0.77 (SD = 0.02), see also Suppl. Figure S21).

**Figure 5.**
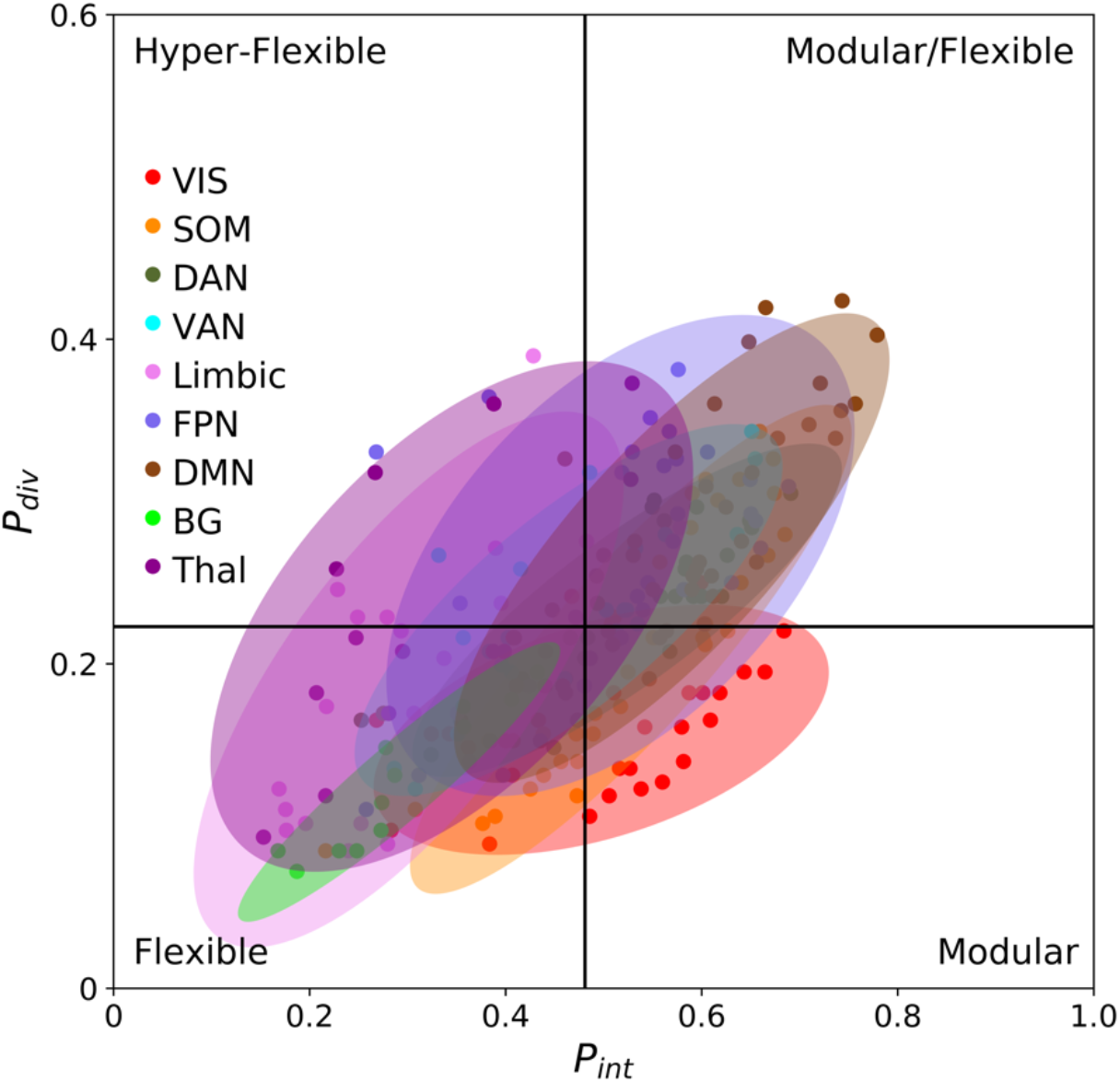
Graph that show the relative flexibility or modularity of individual brain areas grouped according to membership of core RSNs. The x-axis shows the relative frequency with which each area is represented by the model in the time domain as part of any SNC (p_int_). The y-axis represents spatial diversity (p_div_), that is the proportion of the total number of areas that each area shared an SNC with. The vertical line corresponds to mean p_int_ for the whole set of areas and the horizontal line to mean p_div_. There was a strong correlation between p_int_ and p_div_ (r = 0.68). Each ellipse represents a core RSN and is centered at the mean p_int_ and p_div_ for the areas belonging to the RSN where the height and width corresponds to two standard deviations from the mean. The tilt of the ellipses is set by the covariance. Some RSNs had outliers seen as dots outside the ellipse, notably the DMN, FPN and Limbic networks. All but the BG network spanned more than one category. The VIS (visual) stands out by its comparatively low covariance between p_int_ and p_div_.

From these results, we infer that brain areas that fell into the categories “flexible” and “highly flexible” synchronized for short durations with a wider constellation of other areas. Their time-resolved connectivity patterns were therefore less well represented by the eight-area sized SNCs used here. Smaller sized SNC would be more appropriate for this subgroup of brain areas and this issue needs to be further addressed in future studies. Importantly, the flexible/modular categorization offered here is by no means absolute but should rather be viewed as a measure of tendencies of flexibility and modularity for different brain areas relative to each other. Also, the values of the dispersion along the two axes will increase when more SNCs are sampled (see also Suppl. Figure S28).

## 4 Discussion

In this work a novel method has been described that captured both the flexible and modular as well as integrated and segregated nature of functional brain connectivity in the time-resolved domain. The starting point was to investigate the properties of smaller ensembles of cortical and subcortical brain areas (SNCs) which exhibited the strongest positive phase coherence across time. Subsequently, these ensembles were combined into spatiotemporally flexible SNs that in a final step were clustered into MNs based on their main tendencies for either integration (same community assignment and positive phase coherence) or segregation (assignment to different communities and negative or orthogonal phase coherence). To a large extent, MNs were combinations of known core RSNs. In contrast to winner-takes-it-all parcellation schemes, our results show that when individual brain areas were allowed to a priori take part in more than a single subnetwork unit (SNs), variability in connectivity patterns emerged. Importantly, modularity was clearly observed as a core organizational principle in the brain, here expressed as a high degree of segregation between specific SNs and MNs. For the MNs, segregation was most pronounced between DMN and VAN/SOM and between the DMN and SOM/DAN (Suppl. Figure S11). These two segregated pairs of MNs were integrated in some version most of the time but assigned to separate communities as an expression of their negative (or orthogonal) phase coherence vis-à-vis each other. Importantly, the terms integration and segregation were adapted to more precisely define the range of relational properties within and between subnetworks captured by the model. For example, in a graph theoretical framework, integration designates a state where inter-modular connectivity is equally strong or stronger than within separate modules (where a module can refer to a network or submodule within a network, Shine and Poldrack, 2018; Sporns, 2013). Segregation describes the opposite situation where connectivity strength within modules is significantly stronger than between modules. These definitions contain an implicit assumption that a network (or module) is held together as a unit in some form at all times, either as segregated submodules or as part of an integrated whole. It therefore does not account for the case when the general strength in connectivity within modules is very weak or non-existent. In contrast, the definition of integration and segregation employed here accommodates the possibility that a subnetwork (SN) can temporarily dissolve entirely. This was termed disintegration and happened if none of the SNs smallest components (the SNCs) had a sufficiently strong internal phase coherence such that all of its eight areas were assigned to the same community. As shown in Figure 3, assignment to the same community happened on average when internal phase coherence was approximately less than 44° (dPC = 0.72). Therefore, using the definitions presented here, integration was not only opposed to segregation but also to disintegration. A network unit was considered integrated if any of its components were in phase, i.e. the brain areas of the component were all assigned to the same community. Segregation referred to the relationship between networks that were integrated simultaneously but in different communities. Importantly it also quantified the strength of the segregation in terms of degree of phase coherence that was shown to be (for most time points) either approximately orthogonal or negative (see also Suppl. Figure S9). Of note, we only sought to consider the strongest coupled areas in each community (which was a purposefully chosen strategy to reduce contributions from noise) which arguably was those belonging to a SNCs. Therefore, the term segregation as used here refers to the same concept as in graph theory, i.e. the internal coherence within segregated entities are significantly stronger than the coupling between them. In this context, integration refers to coupling both within and between networks that was sufficiently strong to cause assignment of the individual network components to the same community. Thus, our definition and implementation of integration/segregation can be understood in terms of a more fine-grained conceptualization of the conventional separation between them. The more nuanced view endorsed here lies in the fact that the suggested method quantifies the degree of segregation in terms of negative and orthogonal phase coherence relationships and also includes the notion of disintegration. Moreover, our method suggests a more nuanced view of the previously shown mutually strong exclusiveness of integration and segregation in the temporal domain (Shine et al., 2016). It should be noted that our results did not contradict that the brain seems to fluctuate between periods of high and low global integration (Shine et al., 2016). However, our model and results provide a conceptually richer repertoire of brain dynamics in so far that integration was never observed to be entirely global but always co-existed with some amount of segregation present. Moreover, it is likely that some degree of segregation is required within the meta-stability framework to uphold the internal tension that drives the dynamics. Also, we would argue that some level of disintegration and certainly orthogonality between network units are necessary intermediaries for dynamical network reconfigurations across time. In the framework presented here, where subnetworks were flexibly built by smaller components, each with their own time-course, disintegration was prevalent in all SNCs. This fact was reflected in globally low q_int_ values for SNCs. The most frequently integrated SNCs were de facto disintegrated, on average, at 88 out of every 100 time points. Disintegration is also likely to play an important role in periods of low modularity that is not an effect of strong global integration. For example, it was recently shown that intervals of low modularity corresponds to a decrease in dissociation between networks that are otherwise mostly segregated such as the DMN and task positive networks (TPN) (Fukushima et al., 2018). It was also found that periods of high modularity were similar to each other while periods of low modularity were much more variable, where the latter was a reflection of heterogenous interactions between networks. If we view these results in light of the present framework, the variability seen in periods of low modularity would presumably correspond to different stages of transitions from states of full segregation (negative phase coherence) to relatively transient states of integration (positive phase coherence) between submodules of otherwise highly segregated networks. Since such remodulations would be gradual, they would include intermediary time points characterized by partial disintegration within the modular networks in order to facilitate the transitions, as well as a gradual decrease in segregation (from negative to orthogonal phase coherence) between the modules (see also Figure 1G). Depending on the global state, different submodules would drive such reconfigurations at different time points which would further add to the variability.

Given the complexity of spatial gradients for network affinities in the brain (Buckner and Krienen, 2013), the existence of a unique method to represent the full complexity of changing connectivity between brain areas in a concise and unbiased manner seems highly unlikely. Our bottom-up approach was to represent time-resolved neural circuitry in terms of small units which could be combined into a set of highly interleaved spatiotemporally flexible subnetworks. However, while it was rather straightforward to build SNs in which the internal coherence structure was preserved, the subsequent step of aggregating the dynamics of SNs into MNs was made more complicated due to variable interactions between SNs that was hard (or impossible) to neatly delineate. However, it is certainly possible that there are alternatives to the hierarchical clustering algorithm used here to obtain MNs that might provide better solutions. Alas, most clustering algorithms need some kind of thresholding strategy to handle the problem of determining the number of clusters present in the data.

Of note, at the coarsest scale of granularity, our method provided 71 specific meta-networks. This might seem like a large number and it is perhaps prudent to ask the question whether brain dynamics can be reduced to even further at this level? However, if we try to summarize the SNs into fewer MNs by lowering the clustering thresholds of the mean and minimum degree of integration, not only would it result in MNs that describes internally less coherent entities, but their topographical maps would in many instances be rendered less distinct. For example, the VIS network at lower thresholds tended to merge with the SOM, DAN and VAN networks. Also, it should be noted that the relative frequency of integration for most MNs, beyond those ranked among the top twenty, was relatively small (q_int_ ≤ 0.2) (Suppl. Figure S25 and Suppl. Video 1). Taken together, our results further emphasize the notion that delineation of time-resolved connectivity into separate distinct networks such as the classical RSNs does not capture the inherent gradients that are expressed as flexible reconfigurations across borders of relatively strong modularity. The p_int_ and p_div_ measures that graded areas on the flexible/modular axes suggest that not only did RSNs differ on these measures but so did different areas within the RSNs (Figure 5). Importantly, the thalamus, basal ganglia and “limbic” areas were characterized by a high degree of flexibility. The SNCs they belonged to were significantly less frequently integrated than the SNCs that included cortical areas only. At least some parts of the subcortical nuclei are known to have widely distributed reciprocal connections with the cortex (Sherman, 2007; Sherman et al., 2014). The apparent short-lived and diverse cortical-subcortical synchronization is in line with a modulatory role in cortical network dynamics (Bell and Shine, 2016; Jones, 2002; Lumer et al., 1997).

An important step in our method was the choice to use the EMD algorithm to extract intrinsic mode function (IMF) instead of using a narrowly band-pass filtered signal. The IMFs are oscillating modes of a signal derived with a heuristic sifting method that retains variability within the signal while eliminating riding waves to permit instantaneous phase information to be interpretable (See methods 2.4). In this study the third IMF was used since it contained the majority of the power within the traditional 0.01-0.1 Hz frequency band used in resting state fMRI studies. For completeness, we also applied our method to resting-state fMRI data that were filtered using a narrow band-pass filter 0.03-0.06 Hz that corresponded to most of the power of the chosen IMF. While the topological maps were very similar, the most salient difference was the sharp increase in coupling strength between areas following band-pass filtering. The stronger coupling between brain areas was apparent in terms of three times higher mean q_int_ for the SNCs compared to those derived from the IMF and an almost as large rise in mean duration of SNs (7.4 sec vs 2.9 sec). It was also reflected in a general increase in the strength of the average correlation (absolute values) within the recurrence matrices (Supplementary Text and Suppl. Figure S34). Additionally, core areas of the topological maps of the MNs had higher weights reflecting that they were more often part of the network (compare the MN topological maps shown in Figure 2 with the maps shown Suppl. Figure S33). In case of using the EMD algorithm and the third IMF, a similar behavior was observed only when the expansion factor K was increased from 10 to 40 (see Suppl. Text and Suppl. Figures S26 and S27). With a narrow band-pass filtered signal, the frequency range under investigation is clearly specified. Due to the heuristic nature of the EMD, each IMF contains mix of frequencies that is not defined a priori and also can vary across areas within subjects and between subjects. Hence, the IMFs are better adapted to capture variability (both frequency and amplitude modulations) within the signal (Huang et al., 1998; Niazy et al., 2011). Whether this variability is potentially biologically meaningful or rather should be considered an artefact of the EMD algorithm can be debated. As for the present method, the variability in the IMFs affected the overall level of phase coherence between brain areas but not the overall behavior of the method.

Meta-stability as an organizing principle for dynamic interactions has been identified at many levels of brain activity at both faster timescales (EEG, MEG) than the BOLD-signal as well as at slower time-scales, i.e. coordination dynamics (Bressler and Kelso, 2016). In our work, the principles of meta-stability appeared at multiple levels of spatial granularity, from SNCs to change in global connectivity. At the second most fined-grained level (SNCs), meta-stability was observed when investigating the time point to time point changes in brain connectivity directly without averaging across distinct time points. Importantly, a gradual increase in phase coherence of SNCs was observed prior to the time point of assignment to the same community (Figure 3). The progressive increase in phase coherence was a result of the gradual recruitment of brain areas constituting the SNCs. In line with the meta-stability framework, we found a strong relationship between q_int_ (i.e. the degree of assignment to the same community) and duration of integration (Suppl. Figure S14), as well as between duration of integration and peak phase coherence (Suppl. Figure S15). It is important to note that integration occurs during positive as well as negative amplitude fluctuation in the BOLD IMF-signal. In fact, SNCs, SNs and MNs stayed integrated through complete or partial cycles of fluctuation. These findings shine a new light on how to interpret brain dynamics in the context of task-based fMRI where only the relative amplitude of activation/de-activation is conventionally considered to be of interest. From the results presented here, it is plausible to consider the notion that periods of integration between ensembles (SNCs, SNs, MNs) are of equal or greater importance than the relative increase/decrease in BOLD activation, which captures only part of the time period of coherence for a given subnetwork. When contemplating this aspect of the results, it is important to recall that it is the third IMF from the EMD decomposition of the BOLD signal and not the raw time series that has been used here. Thus, the mentioning of amplitude fluctuations (activation and deactivation) is therefore relative to the mean of the third IMF BOLD time series. Moreover, network integration occurs in a shifting context of whole-brain configuration where the recruitment of brain areas take place with some degree of temporal lag. The functional relevance of integration of specific networks is likely to depend on the global state of brain dynamics of which it is a part of. Possibly, the actual role of an integrated network might also be sensitive to the temporal sequence of recruitment of its areas (Mitra et al., 2015).

At a global level, meta-stability was inferred from the apparent presence of quasi-attractors that shaped the aperiodic pattern of recurrence (Figure 4). The change between adjacent time points was small, guided by the presence of strong correlations. Observed over time, patterns emerged where one period of highly correlated state vectors was followed by either an uncorrelated or anti-correlated configuration. This behavior likely corresponds to a quasi-cyclic activation and deactivation of the major RSNs as first shown by Fransson (2005) interspersed with periods of reconfigurations between (sub)networks. Here we did not in detail outline the principal configurations of the quasi-attractors that presumably drive this dynamic (Suppl. Figure S11 provides some indication that one principal configuration is likely to entail integration between DMN and FPN that are segregated with respect to SOM/VAN/DAN). However, we believe it is important to further explore the apparent quasi-cyclic behavior in global activity in more detail, not least in relation to fluctuating periods of high and low modularity (Betzel et al., 2016b; Fukushima et al., 2018; Shine et al., 2016) and efficiency (Zalesky et al., 2014) as described in the existing literature. One reason for this is that (quasi-)cyclicity is a phenomenon abundant in nature. In the brain it is seen at all levels of investigation from molecular, cellular and electrophysiological recordings to studies of behavior and cognition (Buzsaki, 2006). Importantly, cyclicity has been shown to be a major force behind formation of modular organization in nature (Damicelli et al., 2019; Hebb et al., 1950; Holland, 1998). Taken together, cyclicity and modularity have been proposed as the prerequisites for any system to form long-term memory, one of the essential features of many mammalian brains (Holland, 1998). Therefore, the intrinsic quasi-cyclic nature of integration and segregation on the BOLD level is likely a core feature of functional brain organization. As such it is a topic that needs to be further investigated. This does not only pertain to technical advancements in terms of greater spatial and temporal resolution and development of methods that can handle the increased computational burden. It also puts emphasis on the need for a deeper theoretical exploration and advancement to enhance our conceptual understanding of how brain processes creates our internal world as well as how it sustains an interface with the external reality (Raichle, 2010).

Relatedly, the results of this study also point to the need for a more nuanced understanding of the cyclicity of intrinsic connectivity in the context of fMRI-experiments where task trials are commonly timed at specific intervals. There is an abundance of studies in the fMRI-literature that attributes multiple and diverse sensorimotor and cognitive functions to specific networks (http://Neurosynth.org). However, our results and the conclusions thereof lead us to consider alternative perspectives on task-related fMRI. Here, caution is warranted since it cannot be ruled out that some of the previously shown attributions between specific tasks and brain networks have inadvertently been biased by the timing of the task-based fMRI experiment relative to the intrinsic cycles of activity of the subnetworks in question. Thus, it is conceivable that in certain scenarios, a specific brain function is inferred from an experimental task design imaging experiment when in reality the contributions of the subnetwork to brain function might be much more general and even unrelated to the experiment at hand. In fact, the awareness of the problem of timing and interpretation of task activation in relation to intrinsic activity is not new (Huk et al., 2018; Papo, 2013). More than a decade ago it was shown that that the timing of trials in relation to the intrinsic dynamics contribute to the variability seen in fMRI task studies (Fox et al., 2007). It has also been shown that connectivity in task activated areas is not uniform through the duration of trials (Di and Biswal, 2019). Importantly, task activation seems to a large extent to be a mere modification of the intrinsic ongoing activity (Cole et al., 2014). This might indicate that we are not so much responding to the world around us per se as continuously sustaining and aligning an internal version of it adapted to predict and meet our future needs (Clark, 2013; Northoff and Huang, 2017; Raichle, 2010). Therefore, a better understanding of the intrinsic dynamics of the brain is necessary. This would include further elucidating the intrinsic and apparent quasi-cyclic recurrence of spatiotemporal organization across multiple frequencies domains.

### Code availability statement

All code and additional information to reproduce this work or apply the method to a different dataset and/or parcellation scheme can be found at: https://github.com/MarikaStrindberg/TimeResolvedNetworks **`DOI: 10.5281/zenodo.4794699`**

## Abbreviations and key parameters used in the text

dPC: dynamic phase coherence (i.e. cosine of the phase difference)
EMD: Empirical mode decomposition
IMF: Intrinsic mode functions
IPSA: Instantaneous phase synchrony analysis
SNC: subnetwork component (second smallest unit of brain connectivity considered)
SN: subnetwork (middle level unit considered)
MN: meta-network (largest unit of brain connectivity considered)
RSN: Resting-state network
Ensemble: any functional unit i.e. area pair, SNC, SN or MN
BG: Basal Ganglia Network
DAN: Dorsal Attention Network
DMN: Default Mode Network
FPN: Fronto-Parietal Network
Limb: Limbic Network
SOM: Somato-Motor Network
Thal: Thalamic Network
VAN: Ventral Attention Network
VIS: Visual Network
P_div_: spatial diversity (proportion of all areas in the parcellation that each area shared a SNC with)
p_int_: temporal representation (the relative frequency of participation of an area as part of any SNC across the entire scan time)
q_int_: relative frequency of assignment of any ensemble to the same community relative to the whole scan time
q_tov_: relative frequency of assignment of any ensemble to the same community relative to total number of time points of temporal overlap

## Acknowledgements

The computations were enabled in large part by the resources provided by Swedish National Infrastructure for Computing (SNIC) through Uppsala Multidisciplinary Center for Advanced Computational Science (UPPMAX) partially funded by the Swedish Research Council through grant agreement no. 2018-05973. The BrainNet viewer was used for visualization of networks: www.nitrc.org/projects/bnv (Xia et al., 2013). The following sources has in part founded this research: Swedish Medical Research Council (grant numbers, 2017-03043), the Swedish Brain Foundation (grant number, FO2019-0045), The Philipsson foundation, Erik and Edith Fernströms foundation, Fredrik and Ingrid Thurings foundation, Jerring Foundation, Sällskapet Barnavård, Stiftelsen Samariten, Svenska läkarsällskapet. PF was supported by the Swedish Research Council (grant no. 2016-03352) and the Swedish e-Science Research Center.

## Supplementary Text and Figures to Strindberg et al., 2021

### 1. On the stability of the Louvain algorithm

The Louvain algorithm does not generally give the exact same partitioning of a dataset between runs. To evaluate the downstream effect of different partitions we ran the Louvain algorithm 100×100 times. The seed pairs resulting from each run were used to compare partitions. Across hundred runs, the most frequently occurring seed pairs were calculated such that the run that contained the seed pairs with largest frequency across runs was selected. This procedure was repeated 100 times and the overlap between the resulting top seed pairs was 91%, i.e. 91% of pairs were always the same across 100×100 runs. Increasing the number of iterations to 200×100 did not improve the reproducibility but increased the run time significantly. Therefore, to let the Louvain algorithm run 100 times and chose the solution that contained the most commonly occurring seed pairs across runs was deemed sufficient.

### 2. Algorithm to identify SNCs

The algorithm to find gradually larger SNCs started with identifying seed pairs. The seed pairs represented the pair of highest pairwise q_int_ from the perspective of each area in the parcellation. At each iteration the relative frequency (q_int_) with which each pair was assigned to the same community as all other possible pairwise combinations of areas were calculated. Since only the most frequent constellations from the point of view of each area were of interest, an expansion factor had to be set for each iteration. An expansion factor K = 10 meant that for each seed pair, the top (i.e. highest q_int_ values) ten pairs with which the seed pair was most frequently found to be assigned to the same community with were kept (Figure 1B). The ten resulting SNCs of size n = 4 (brain areas) for each seed pair where then used as seeds in the next iterative step where the same process was repeated to find SNCs of size n = 6 etc. For a discussion on effect of variations in the K-parameter see the end of this supplement.

Admittedly, the iterative steps described here could in principle be repeated for increasing values of n to some limit *L* depending on the size of the parcellation. However, in order to establish a halting point for the algorithm, it was necessary to estimate an appropriate size (i.e. n, maximum number of areas) of the SNCs that would balance flexibility and specificity at the next coarser level of spatial granularity (SNs). In this context it is also essential to point out that the appropriate size of SNCs will also depend on the initial spatial granularity, that is, the number of brain areas defined by the brain parcellation scheme used. We reasoned that an acceptable criterium for the final size of SNCs would be where a clear separation could be observed in the distributions of the mean of q_int_ values for SNCs and randomly generated components of the same size. In theory, in an unstructured random network, the number of possible combinations of areas that at any time would integrate into any of three communities is vast. For components composed of pairs of areas, such a probability distribution would be narrowly centered around 0.33. However, since the brain is fundamentally modular in its basic organization, some combinations of areas are much more likely to appear than others. Thus, the difference between random constellations and representative components would therefore increase with increasing size of the SNCs. This effect is shown in Figure S6, where a sample of 10 000 random combinations of *n* unique areas of progressively larger size (i.e. n = 2, 4, 6, 8) were drawn from the set of all possible combinations of pairs of brain areas and compared to empirically (i.e. derived from q_int_ values). For n = 4, the distribution of SNCs had a mean of q_int_> 4 times larger than the corresponding mean q_int_ value for random components. At size n = 6, the distribution of SNCs had a mean of q_int_ that was 13 times larger than for random components. The difference at n = 8 was even larger, with a mean for q_int_ for the SNCs 55 times larger compared to the random components of the same size. Importantly there was no overlap between the two distributions at this level. Therefore, SNCs of size n = 8 were judged to have a distribution adequately separated from the chance distributions and was used as the final size of the SNCs in our analysis.

### 3. Grouping SNCs into SNs

Two guiding rules were used for grouping SNCs into SNs: rank and degree of spatial overlap (OV). Rank was simply the average (q_int_) across subjects such that the SNC with highest q_int_ had rank 1. The SNC with rank 1 was then used as the first centroid and spatial overlap was defined relative to this SNC. As a rule, SNs should at all time-points be internally coherent meaning that their constituting SNCs should, at all time-points of simultaneous integration, be assigned to the same community. The spatial overlap where this this “no-split” criteria was fulfilled was heuristically decided. For the parcellation (N = 236) and the size of the SNCs (n = 8) used here, OV was found to be five areas. Hence, all SNCs that shared at least five out of eight areas with the SNC of rank 1 was grouped into one SN and removed from the list of available SNCs. Next, the same steps were iterated for the remaining SNCs until all SNCs were merged into SNs. A schematic illustration of the merging of SNCs in to SNs are shown in Figure 1c and Suppl. Figure S7.

The choice of OV beyond the no-split criteria defined above is conceptually analogous to the choice of number of components in ICA. Increasing the constraint on overlap leads to an increased number of SNs that mostly were fragments of larger SNs. Similarly, a more relaxed constraint on the degree of spatial overlap resulted in fewer and less separated SNs but with time-points where some SN would be split in different communities.

### 4. Cluster thresholds to form MNs

For each of the cluster solutions, the two parameters u_1_ and u_2_ were calculated. The parameter u_1_ denotes the mean degree of integration (between the SNs that constitute the MN) relative to temporal overlap (q_tov_). The other parameter u_2_ denotes the minimum degree of integration (again, between SNs that constitute the MN) relative to the temporal overlap (q_tov_). For each cluster solution C, let c be the individual MNs and let X be the set of pairwise relationships between SNs in C. First, we calculate the mean integration (m_c_) within the MN as:

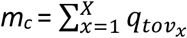

where q_tovx_ is the frequency with which SN-pair *x* is assigned to the same community relative to total time points of temporal overlap. Then, we can calculate the two parameters u_1_ and u_2_ for each cluster solution C as:

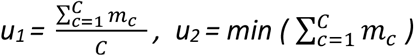

Here, we made the choice to set the cluster thresholds high so that the first cluster solution that fulfilled *u_1_* ≥ *0.95* and *u_2_ > 0.8 was chosen*.

### 5. Results for pairwise areas

In a preparatory step to identify SNCs we analyzed the pairwise relationships between all brain areas in terms of the relative frequency with which they were assigned to the same community (q_int_). The mean q_int_ = 0.39 range [0.28, 0.63] was significantly higher than would have been produced by chance (Suppl. Figure S8). There were no pairs of areas that never were assigned to the same community. The strongest coupling was seen between homologous hemispheric pairs and regional neighbors (Suppl. Figure S8). This result is well in line with what is known about the general principles of RSN topology where homologous hemispheric pairs show the strongest stability at shorter scan time (Gonzalez-Castillo et al., 2014). Of the pairs that had a q_int_ > mean q_int_ + 4 SD, all were homologous hemispheric pairs or intra-modular pairs. Of pairwise areas that had q_int_ > mean q_int_ +3 SD, the vast majority of them (93%) also belonged to the previously mentioned categories of pairs. The remaining 7% consisted of pairs from different modules (as labeled by the Shaefer 7 Networks parcellation). The majority of these inter-modular pairs (54%) connected between the DMN and FPN. The remainder were combinations of the control networks (FPN, DAN, VAN) and the somatosensory network (SOM). These patches of strong connectivity across modularity borders were also subsequently reflected in the SN and MN topology (see Results section 3.1.2 and 3.3). Conversely, the areas with the weakest strongest pairwise connectivity (q_int_ < mean q_int_ -2SD) were mostly pairs that connected between the DMN and VAN.

Finally, on the pairwise level, we also estimated to what extent the q_int_ measure captured similar affinities between brain areas as the commonly used Pearson’s correlation method. We therefore computed Pearson’s r correlation between the time-series of all brain areas and computing the mean taken across subjects. This was done both for the IMF 3 time-series as well as the “raw” BOLD time-series (prior to EMD) (see Suppl. Figure 8). Similarity between the q_int_ IMF3 matrix and the two Pearson’s correlation matrices (IMF 3 and raw BOLD respectively) was evaluated using a Mantel test implementing Spearman’s correlation. Similarity was found to be substantial between both the q_int_ IMF3 matrix and the Pearson’s r IMF3 matrix (r = 0.63, p = 0.000006), as well as between the q_int_ IMF3 matrix and the Pearson’s r raw BOLD matrix (r = 0.77, p = 0.000003).

### 6. The effect of increasing the expansion parameter K

Like any other proposed method that aims to achieve similar goals, a line had to be drawn deciding which parts of the gradients of connectivity that should be sampled and which were to be left out. At each iteration in the procedure to find SNCs, the distributions of q_int_ for all possible combinations had a right sided tail. Though the tails themselves were distinct from the main peaks of the distributions, they contained no obvious point at which a threshold could be set. The choice to use the top ten constellations (K = 10, pairs of brain areas) was therefore arbitrary. However, it was assumed that the most significant relationships just below the threshold would likely be captured in subsequent steps of the iteration. Indeed, the stark convergence of SNCs (see Results section 3.1) as the SNCs grew in size was interpreted as a confirmation that our initial intuition was correct, i.e. that several different starting seeds gave rise to the same SNC at the final stage. Another important aspect of the choice of threshold was sparsity in the representation of flexibility of SNs in terms of number of SNCs. The aim was to avoid oversampling the SNs by avoiding to represent every unique combination of brain areas of the SN as an individual SNC. There is a vast but finite number of possible SNCs associated with some frequency of integration (i.e. q_int_ > 0). Oversampling SNCs would risk causing a loss in distinction between networks and a loss of specificity from the perspective of each individual brain area, eventually leading to one big undifferentiated network.

For completeness, we explored the effect of an increasing K for the purpose of finding a range of K (K = 10, 20, 40) where subnetworks would not be over-sampled and hard to separate into distinct networks, nor under-sampled and the results susceptible to chance fluctuations in the data. To estimate the latter, the degree of convergence between the iterative steps were also evaluated. A high degree of convergence between steps would be an indication of robustness in the results. In the following is a brief comparison between expansion factors K = 10, K = 20 and K = 40 based on the same community run (Louvain).

The distributions of SNC q_int_ were compared for setting the expansion factor K = 10, K = 20 and K = 40. The shape of the distributions was found to be stable across different choices of K (see Suppl. Figure S22 and Table S1). The iterative convergence, i.e. the proportion of initial seeds that resulted in the same final SNCs, increased slightly with increased K (see Suppl. Table S2). With increasing values of K, the relative proportion of SNCs with low internal phase coherence grew larger (see Suppl. Figure S23.) The number of SNs and MNs increased with larger values of K (Suppl. Table S3). The mean duration of SNs increased from 2.9 s for K = 10 to 3.1 s for K = 40 (Suppl. Table S4). However, there was a no significant change in mean q_int_ of MNs with increased values of K (Suppl. Figure S25) despite the significant increase in number of MNs. The weights of the core areas in the topographic maps of the MNs increased substantially (see Suppl. Figures S26 and S27) with increased K since more SNCs were sampled.

### 7. Using a band-pass filtering apporach rather than the EMD and IMF procedure to extract spontanoues BOLD fMRI signal fluctations

For comparison, we also applied our method to band-pass filtered BOLD signals rather than using the IMF3 component from the EMD algorithm. A narrow band pass filter (0.03-0.06 Hz) was chosen, since it mimics the power of frequency range of the third IMF. Across all subjects and areas there was a high correspondence between the two sets of signals in terms of correlation (r = 0.68, for an example see Suppl. Figure S29). However, the signal variance was larger for IMF3 since it consists of a slightly larger mix of different frequencies. The reduction in signal variability gave rise to stronger coupling between areas which also resulted in an increase in mean q_int_ for SNCs (mean q_int_ = 0.18, SD = 0.07, range [0.05, 0.39]) compared to q_int_ values for SNCs derived from the third IMF (mean q_int_ = 0.06, SD = 0.02, range [0.01, 0.12]). It also led to an increase in the mean number of areas that were represented as part of any SNC at any given time-point (70% (SD = 5%) compared to 46% (SD = 5%) for IMF3 based calculations). Band-pass filtering also caused a significant increase in mean duration of SNs from 2.9 s in IMF3 to 7.5 s (SD = 2.1), see also Suppl. Figure S30. The number of SNCs deceased slightly (n = 28601) but the average number of communities that had integrated SNs at any given time-point remained unchanged compared to IMF3 (n = 2.4). Interestingly, the usage of band-pass filtered signal rather than the IMF3 signal led to higher P_int_ values but little change in P_div_ (Suppl. Figure S31). Correlation between the two was high for both P_int_ (r = 0.94) and P_div_ (r = 0.86). Normalized versions of the distributions are provided in Suppl. Figure S32 which shows that the relative distribution among areas was only minimally affected by the difference in signal pre-processing. The topographic maps of the top ten MNs yielded by the usage of the band-passed signal are shown in Suppl. Figure S33. The weights of the core areas in the maps are significantly higher (brighter yellow color) compared to the IMF3 derived maps for the same K = 10 (see for comparison Figure 2). Finally, the strength in the recurrence between time-resolved vectors increased (compare Figure 4 and Suppl. Figure S34). This was seen in a general increase in the strength of the average correlation (absolute values) within the recurrence plots (brain areas band-pass r = 0.20 vs brain areas IMF3 r = 0.11, SNs band-pass r 0 0.21 vs SNs IMF3 r =0.14, MNs band-pass r = 0.25 vs MNs IMF3 r = 21.

### Supplementary Figures

**Figure S1.**
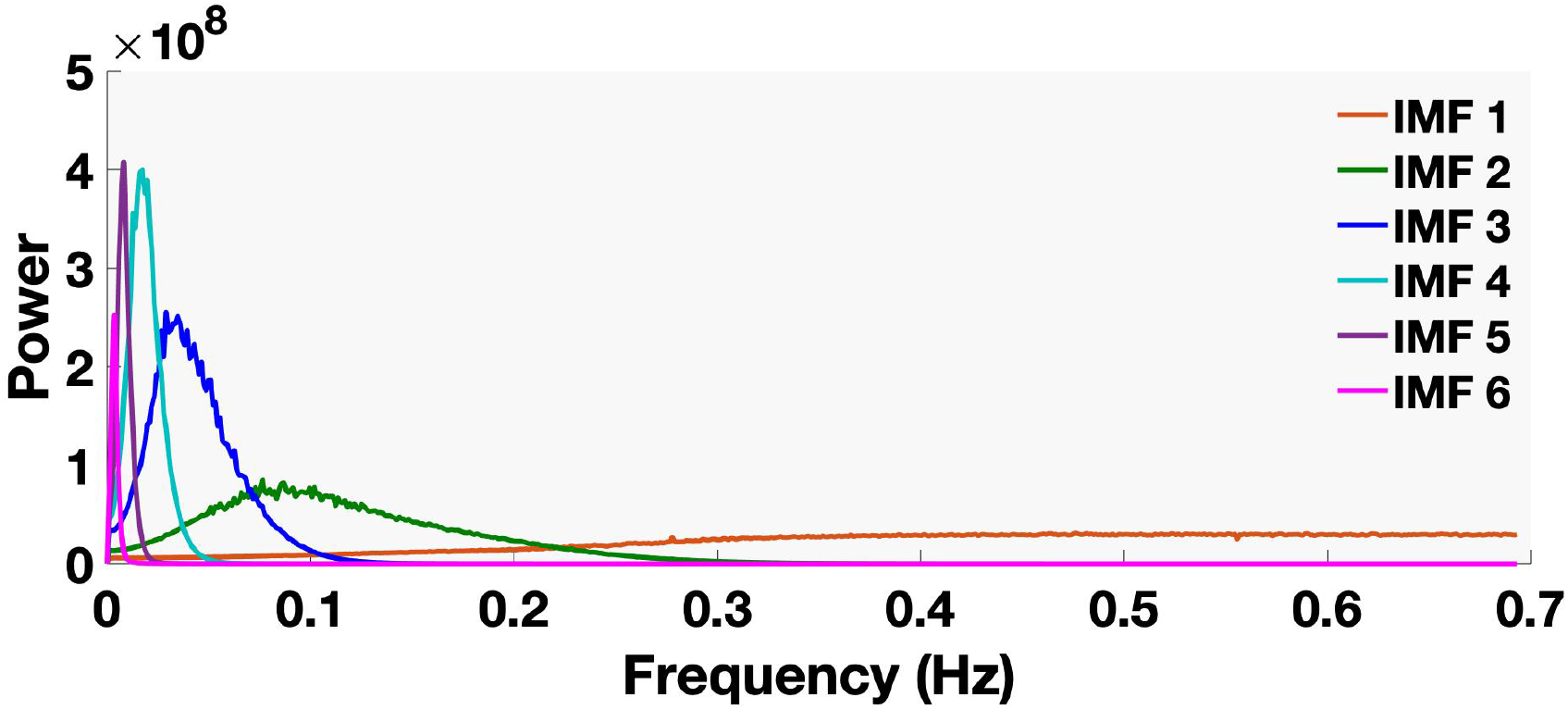
Mean power spectra for IMFs 1 to 6 averaged across all brain areas and all subjects. In the present analysis, we opted to use the IMF3 component. It was chosen since the power spectra of IMF3 closely matched the low-frequency range of spontaneous signal fluctuations that is commonly considered in fMRI resting-state studies.

**Figure S2.**
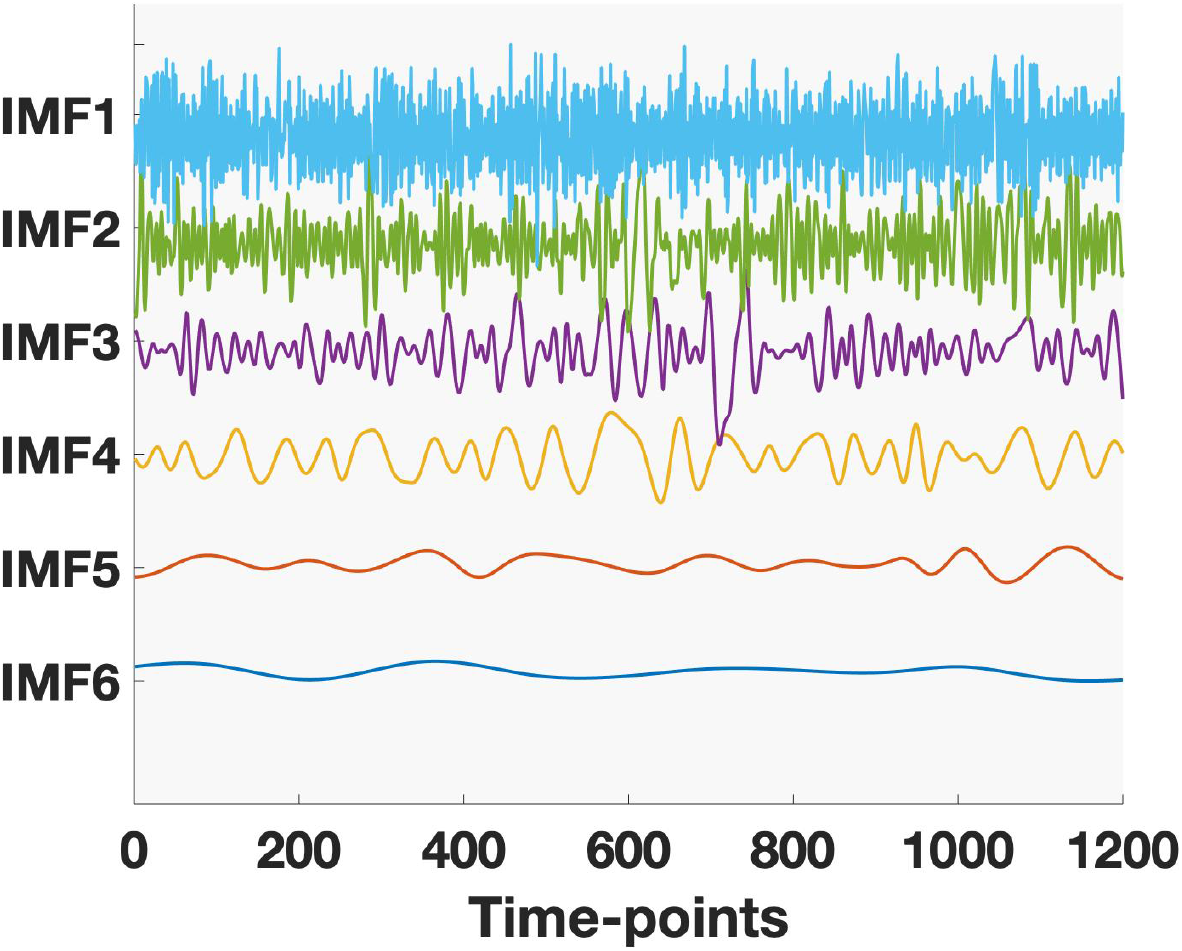
An example showing the first six BOLD IMF signal time-series derived from a single brain area in a single subject.

**Figure S3.**
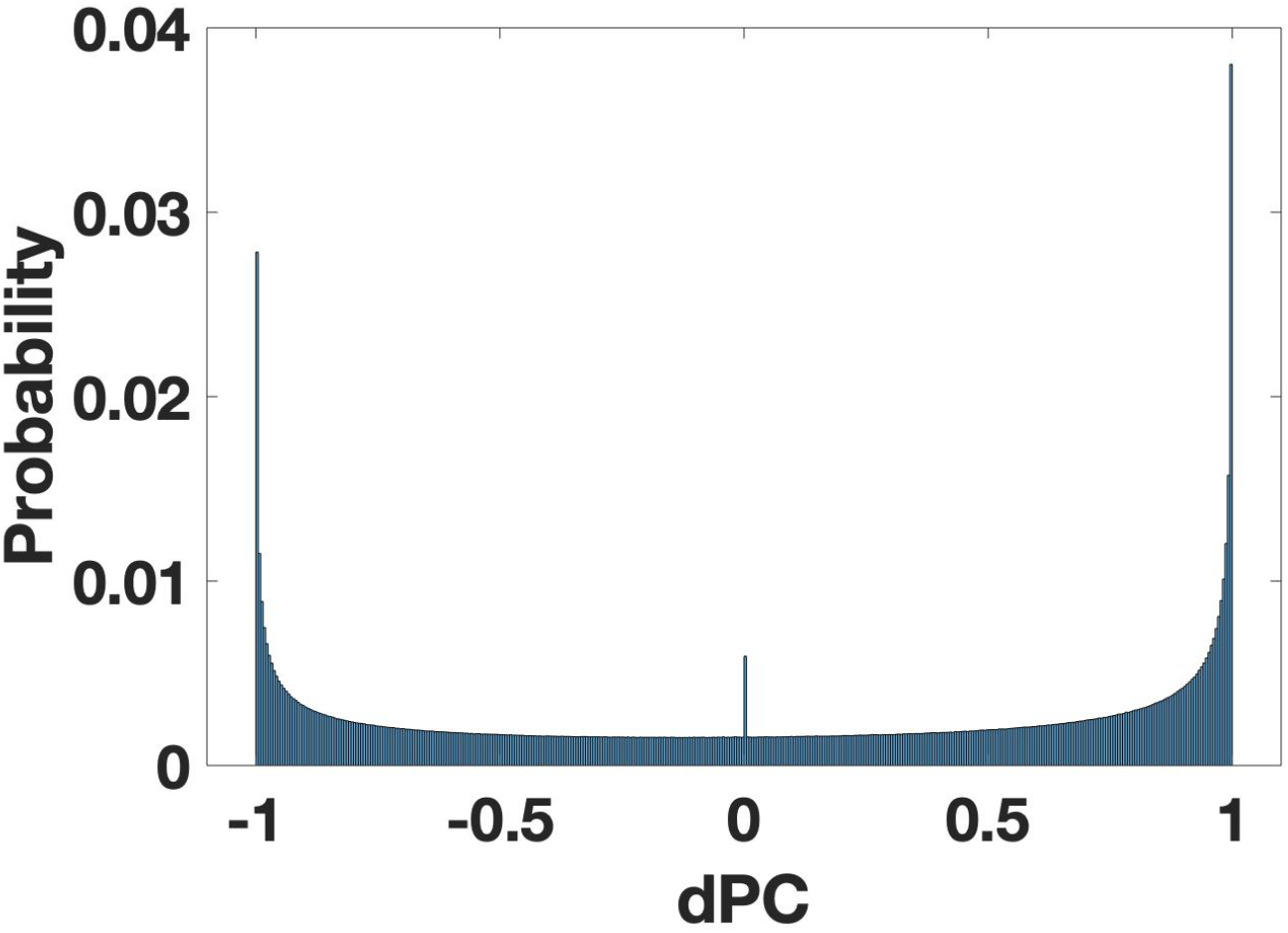
This figure shows the distribution of phase difference values (dPC) computed for one subject (all areas and all time-points). dPC represents the cosine of the phase difference (range [-1,1]). All subjects had similar distributions. Peaks in the distribution were observed at dPC = -1, 0 and 1.

**Figure S4.**
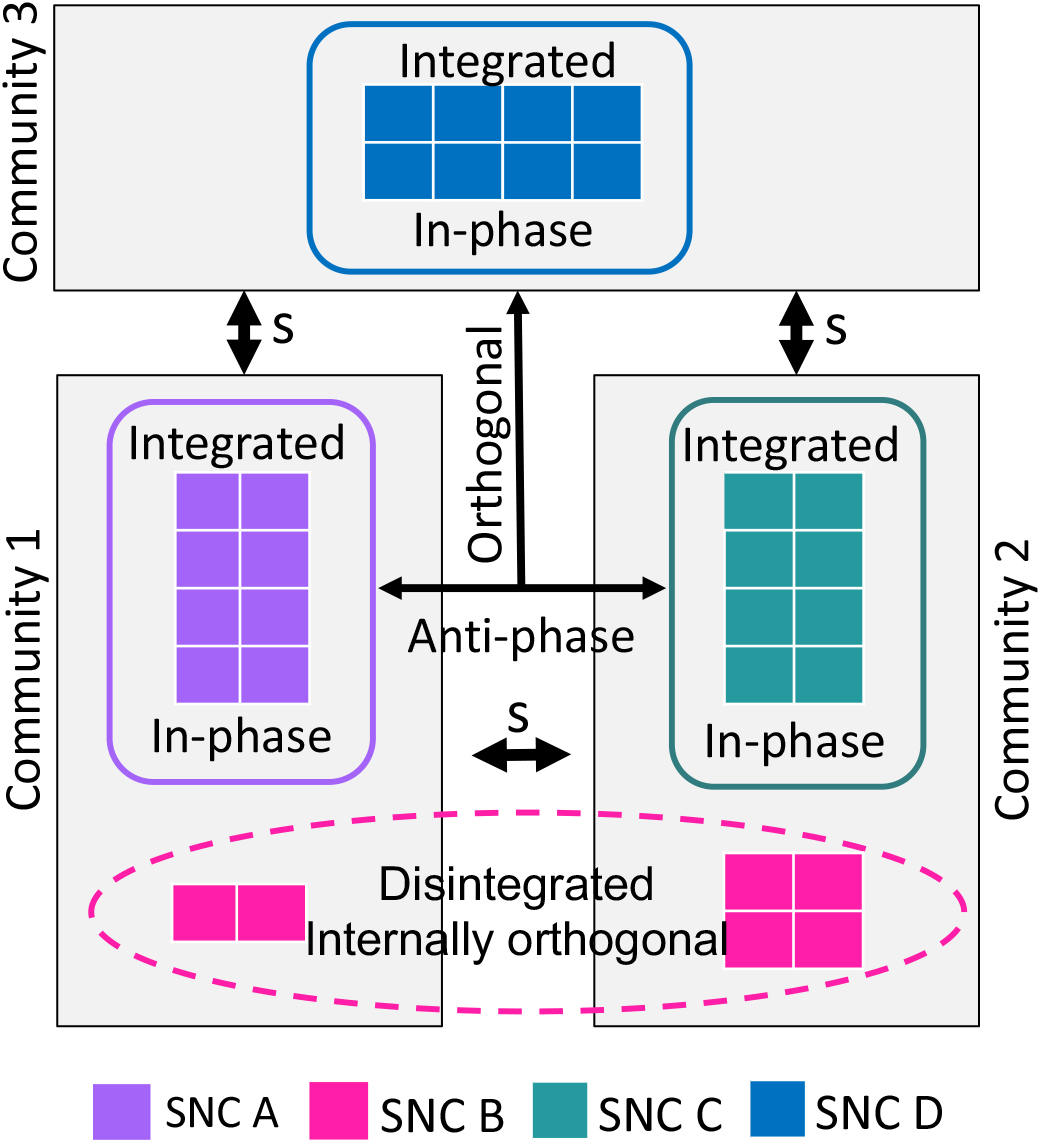
Schematic representation that delineates integration, segregation and disintegration in Subnetwork components (SNCs) at a given point in time. The same principle holds for subnetworks SNs - joint representation of a set of SNCs. Each SNC consists of eight areas. The SNCs depicted in purple (A) and green (C) and blue (D) are internally integrated (i.e. their time-series are in relative phase with each other), but assigned to different communities and are therefore segregated vis-à-vis each other (illustrated by black double arrows). On average, the time-resolved phases between segregated networks are approximately opposite (i.e. in anti-phase, e.g. A vs C) or orthogonal (i.e. A vs D and C vs D). The SNC shown in pink (SNC B) is disintegrated. The lack of internal phase coherence scatters its areas across two communities. This means that SNC B is not a coherent unit at this time-point which is reflected by a zero in its binary time course.

**Figure S5.**
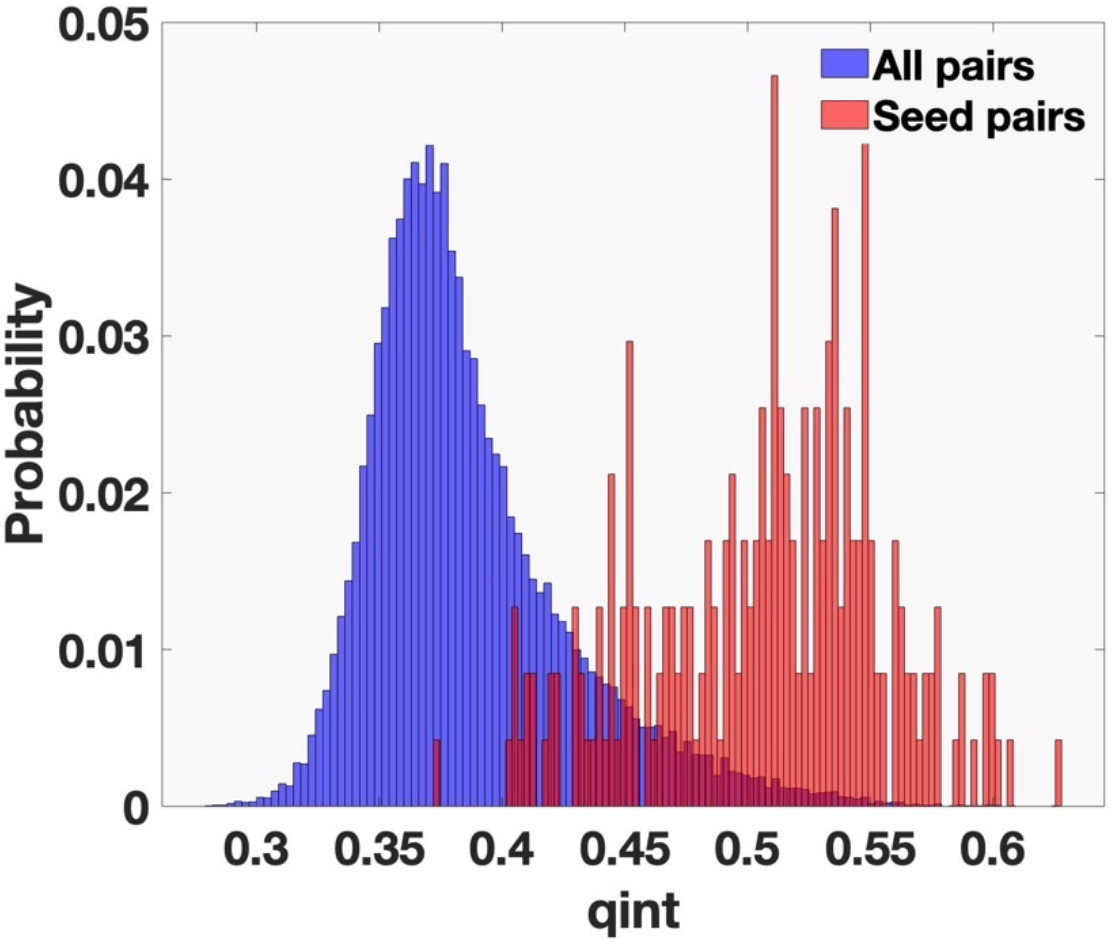
Distribution of q_int_ values for seed pairs of brain regions (red) compared to all possible pairs (blue). The distribution of q_int_ for seed pairs was approximately spread across the right tail of the corresponding distribution for all possible pairs of brain areas.

**Figure S6.**
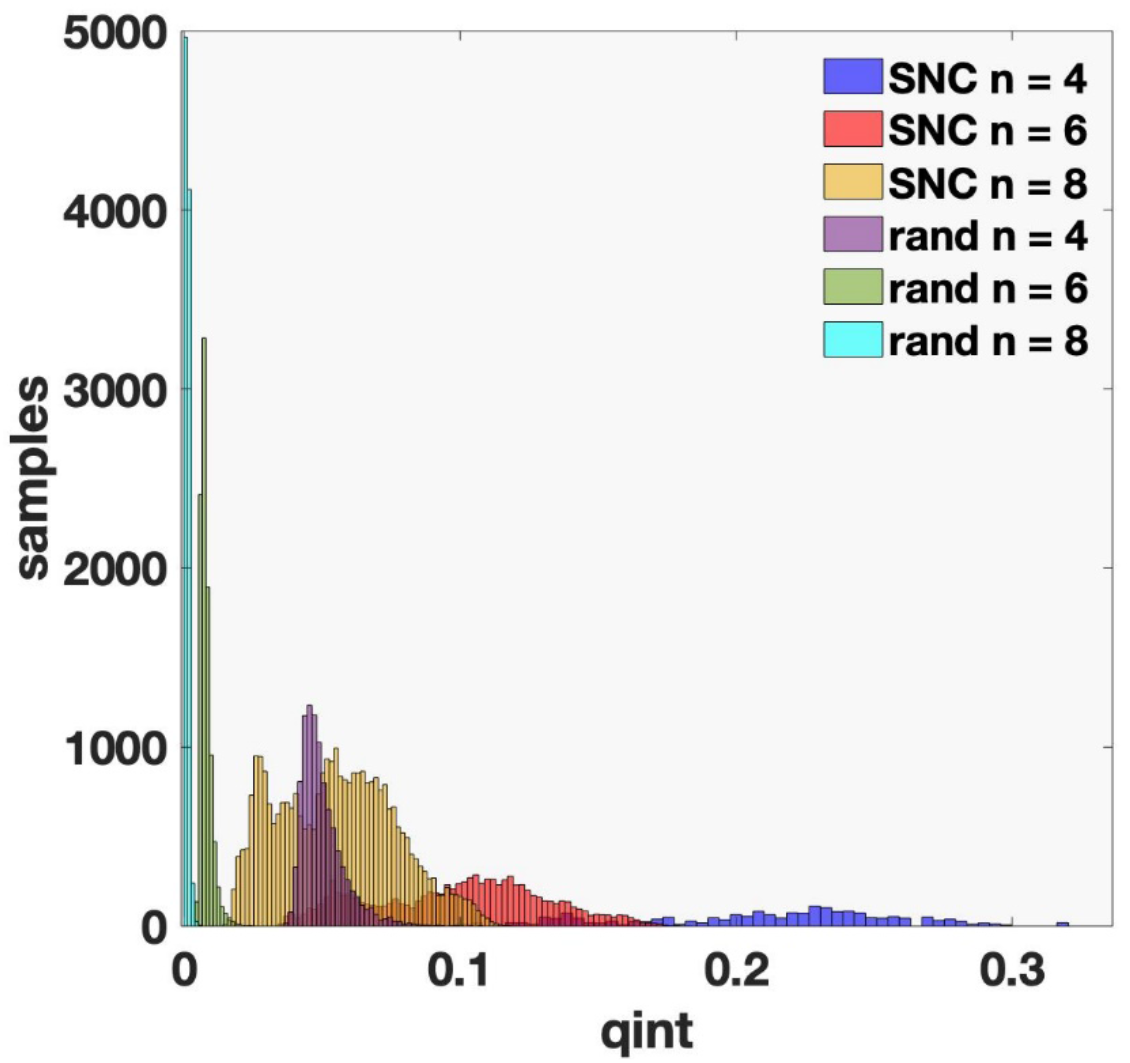
This figure shows the distribution of the parameter q_int_ (frequency of assignment to the same community) for different sizes of subnetwork components based on random combinations of brain areas (rand) as well as the SNCs derived from the selection process described in the Supplementary Text. The randomly composed components were used to estimate the background probability distributions from the empirical data. The aim was to find a size n for the empirically derived SNCs where the q_int_ distribution of the SNCs was significantly separated from the chance distribution. The estimation of distributions of q_int_ from random combinations of brain areas were constructed from ∼10 000 random samples at each size of SNC (number of brain areas (n) = 2, 4, 6 and 8). For SNC size n = 8 there was no overlap between the empirical and the probability distributions which motivated the choice to use of eight area SNCs. Random component n = 8 (turquoise): max **q_int_** = 0.010, min **q_int_** = 0.0005, mean **q_int_** = 0.001. SNCs of size = 8 (yellow): max **q_int_** = 0.119, min **q_int_** = 0.014, mean **q_int_** = 0.055.

**Figure S7.**
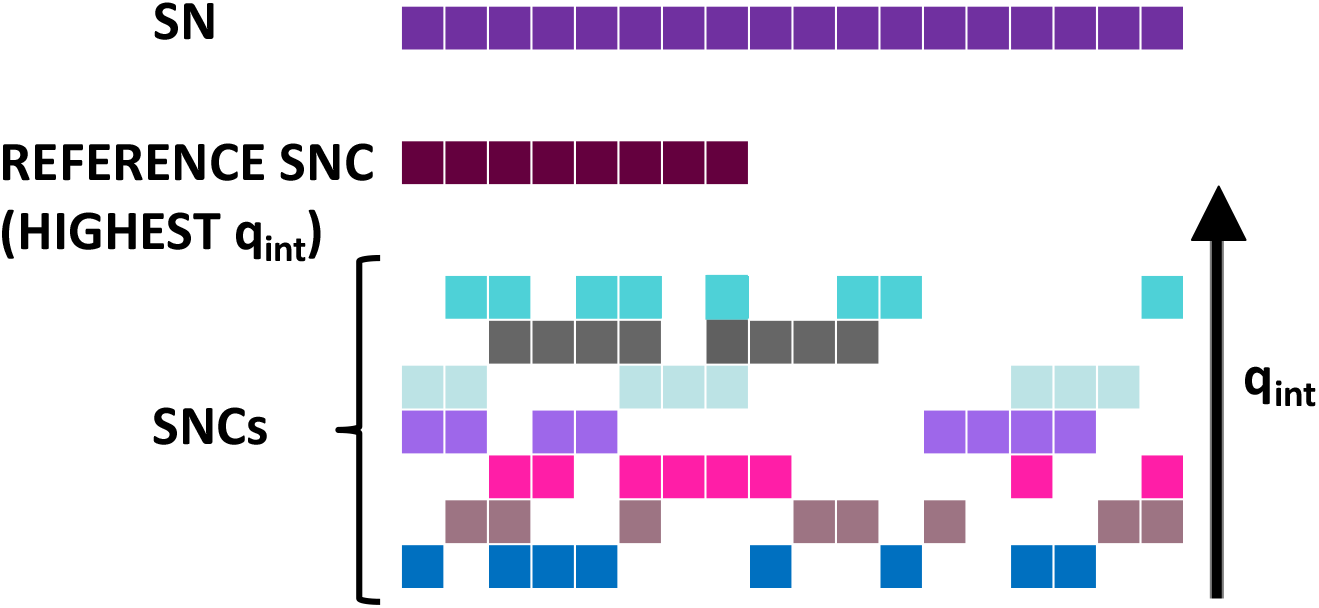
A schematic illustration of the hierarchical procedure of merging SNCs (subnetwork components) into SNs (subnetworks). This merging was based on the degree of spatial overlap with the SNC with the highest q_int_ value (reference SNC, second row in the figure that is not already part of another SN). The SNCs (depicted in the last 7 rows of the figure) that shared at least five parcels with the reference SNC were merged together into a subnetwork (SN, top row) and then removed from the set of ungrouped SNCs. Among the remaining SNCs, the one with highest q_int_ was used as new reference SNC and the process repeated to find the next SN. Each SN was associated with a time series that was the sum of the time series of the incorporated SNCs.

**Figure S8.**
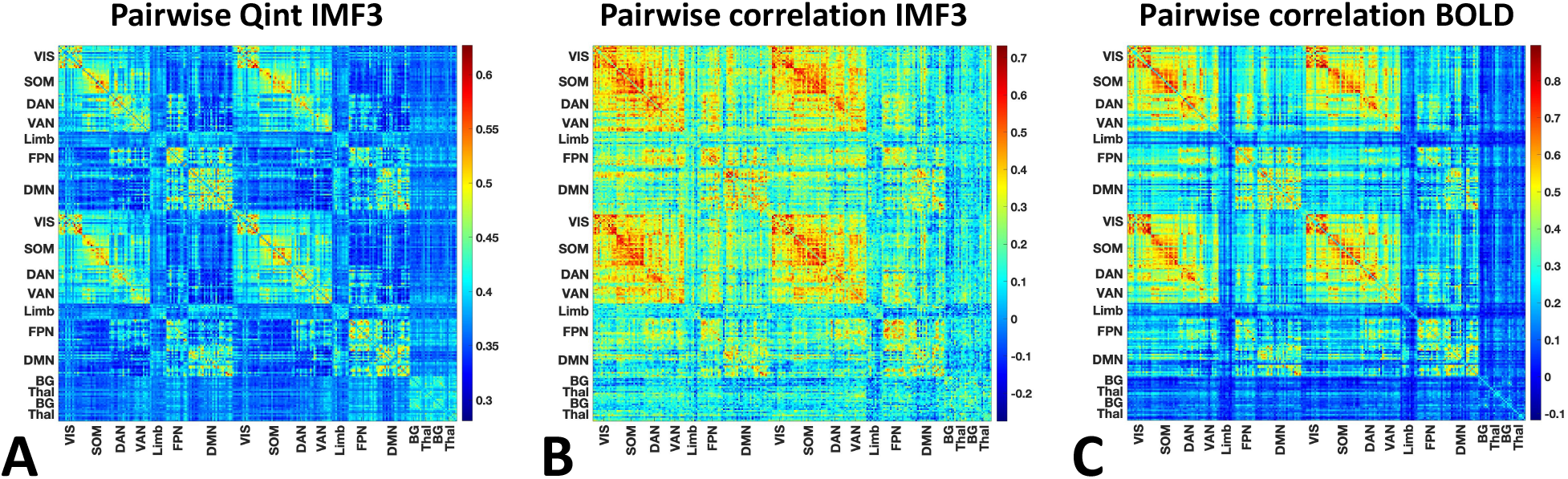
Comparison of the q_int_ measure in relation to the Pearson’s correlation for pairwise brain area relationships across subjects. (A) q_int_ matrix (mean across subjects): mean q_int_ = 0.39 (range [0.28,0.63]), (B) Pearson’s r (mean across subjects) between intrinsic mode function (BOLD IMF3), mean r = 0.21 (range [-0.26,0.73]). (C) Pearson’s r (mean across subjects) between BOLD fMRI time-series prior to the empirical mode decomposition, mean r = 0.27 (range [-0.10,0.90]). The degree of similarity between the q_int_ matrix (A) and the correlation matrixes (B and C) was quantified using a Mantel test with Spearman’s correlation. It was found to be substantial for both the IMF3 matrix (B) (r = 0.63, p = 0.000006), and the raw BOLD matrix (C) (r = 0.77, p = 0.000003).

**Figure S9.**
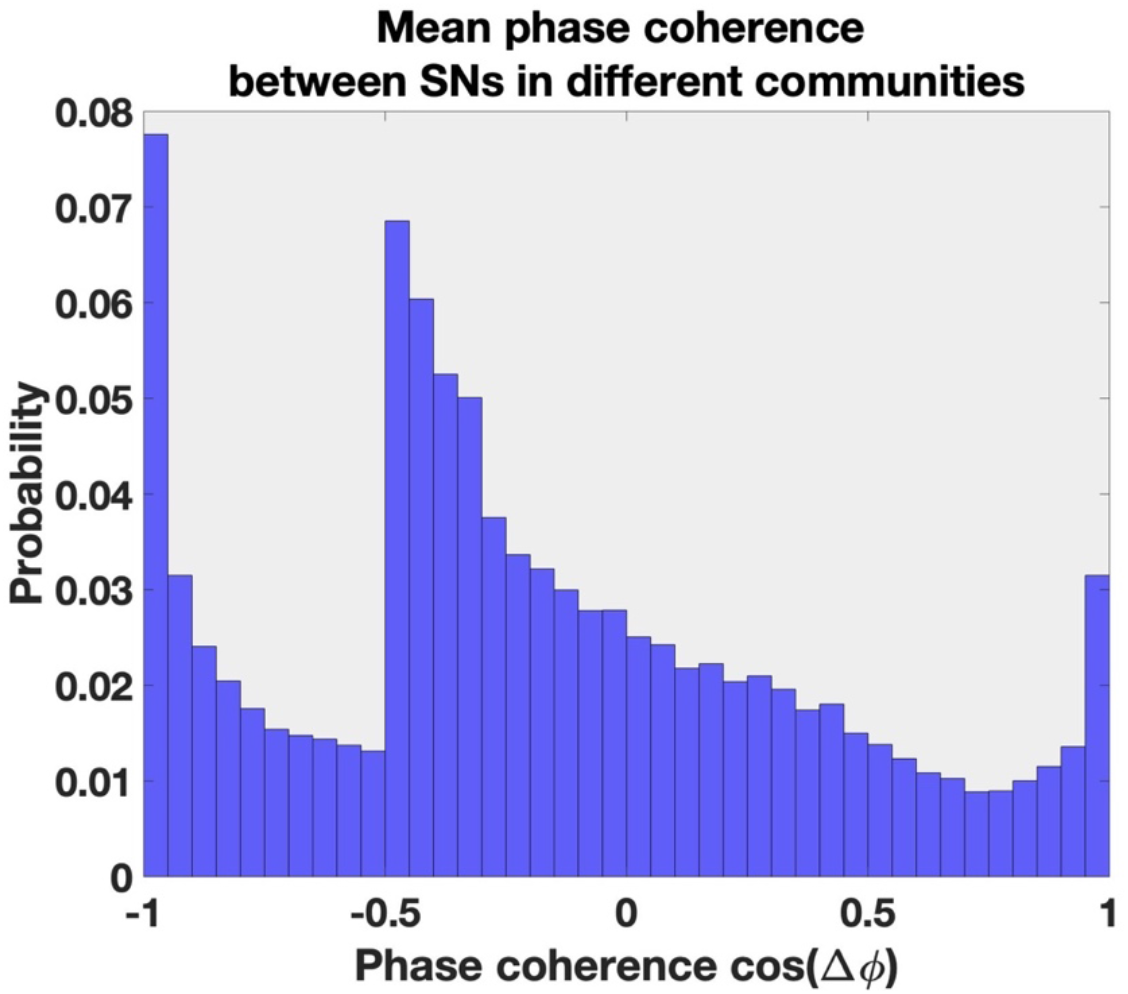
Distribution of mean phase coherence between segregated SNs, i.e. SNs assigned to different communities. If we make a simple division of the dPC values along the x-axis into three groups such that dPC values between -1 and -0.5 signify negative phase coherence, dPC values between -0.5 to 0.5 signify orthogonality and values between 0.5 and 1 is classified as positive phase coherence, we can deduce that the majority of segregated relationships between SNs can be described as either negative or orthogonal phase coherence. A minority however will be classified as positive phase coherence. Ideally these SNs should have been assignment to the same community. This finding indicates that there might exist a better algorithm for community detection of phase coherence data than the Louvain algorithm.

**Figure S10.**
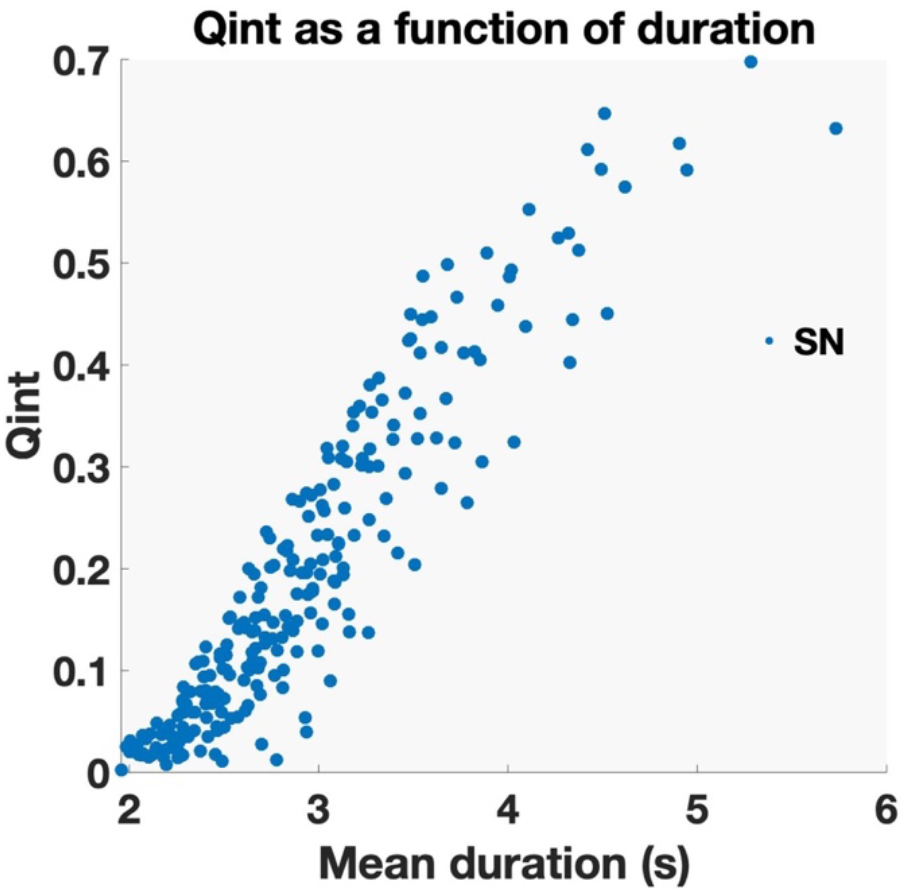
The parameter q_int_ as function of duration of integration at the level of subnetworks (SNs) The figure shows the mean duration of integration plotted against mean q_int_ for all 240 SNs. The duration of integration was strongly related to the total number of SNCs that was assigned to each SN. The average duration of integration for SNs was 2.9 s (SD = 0.7, range [2, 5.7 sec]).

**Figure S11.**
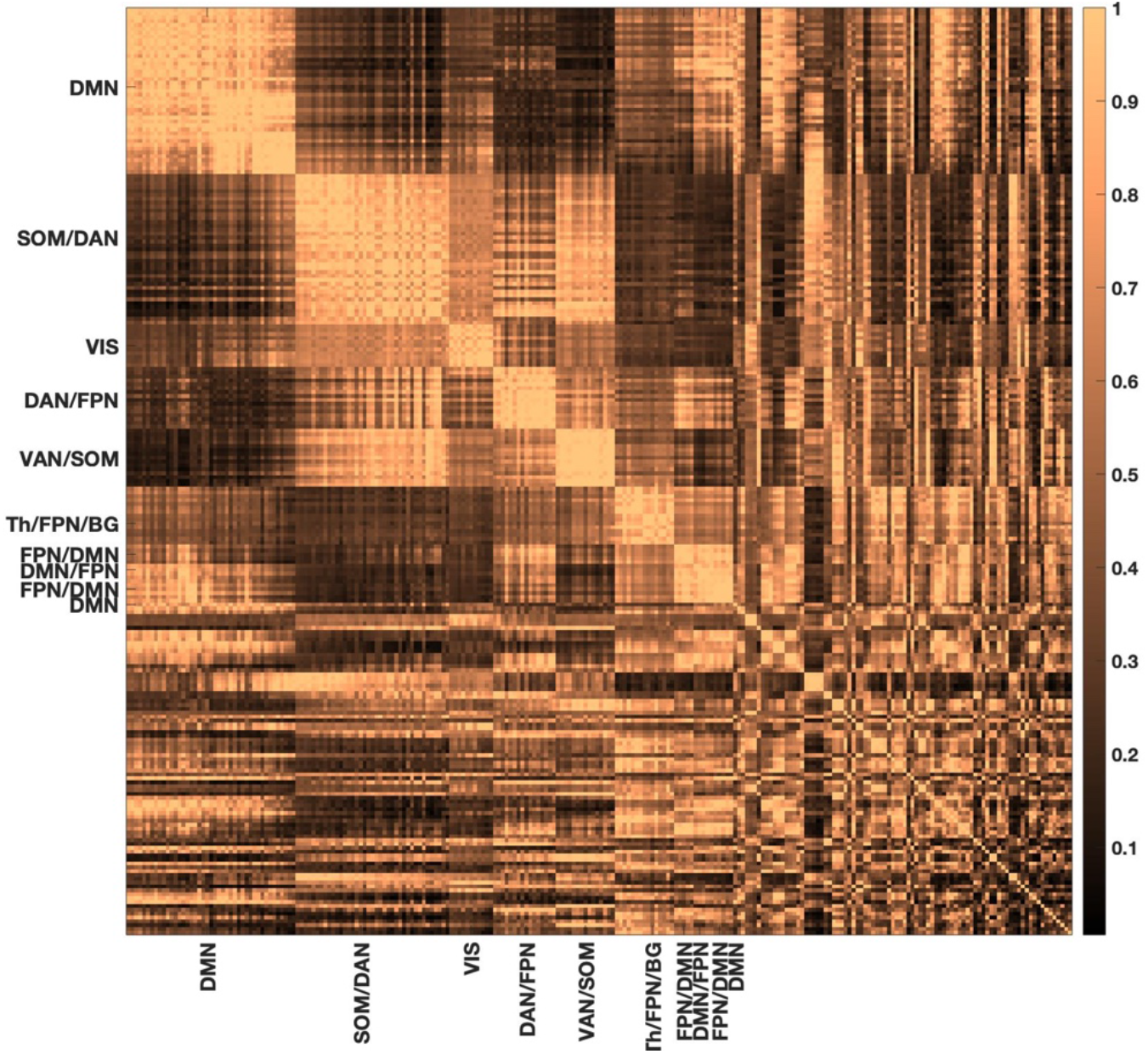
Mean degree of integration between MNs and SNs relative to time of temporal overlap q_tov_. The SNs (n = 240) were sorted by MN (n = 71), and MNs are sorted by largest q_int_ value. Within MNs, the SNs were also sorted from highest to lowest q_int_ values. Due to spatial constraints only the top 10 MNs are named (see Figure 2). Color scale: q_tov_ = 1 (bright copper, SNs were always integrated in the *same* community at times of temporal overlap). q_tov_ = 0 (black, SNs were always integrated in *different* communities at times of temporal overlap). The clearest segregation is seen between DMN and SOM/DAN as well as between DMN and VAN/SOM. Importantly, for 2% of pairwise relationships, integration was always the case. In contrast, even the most segregated SNs were sometimes integrated.

**Figure S12.**
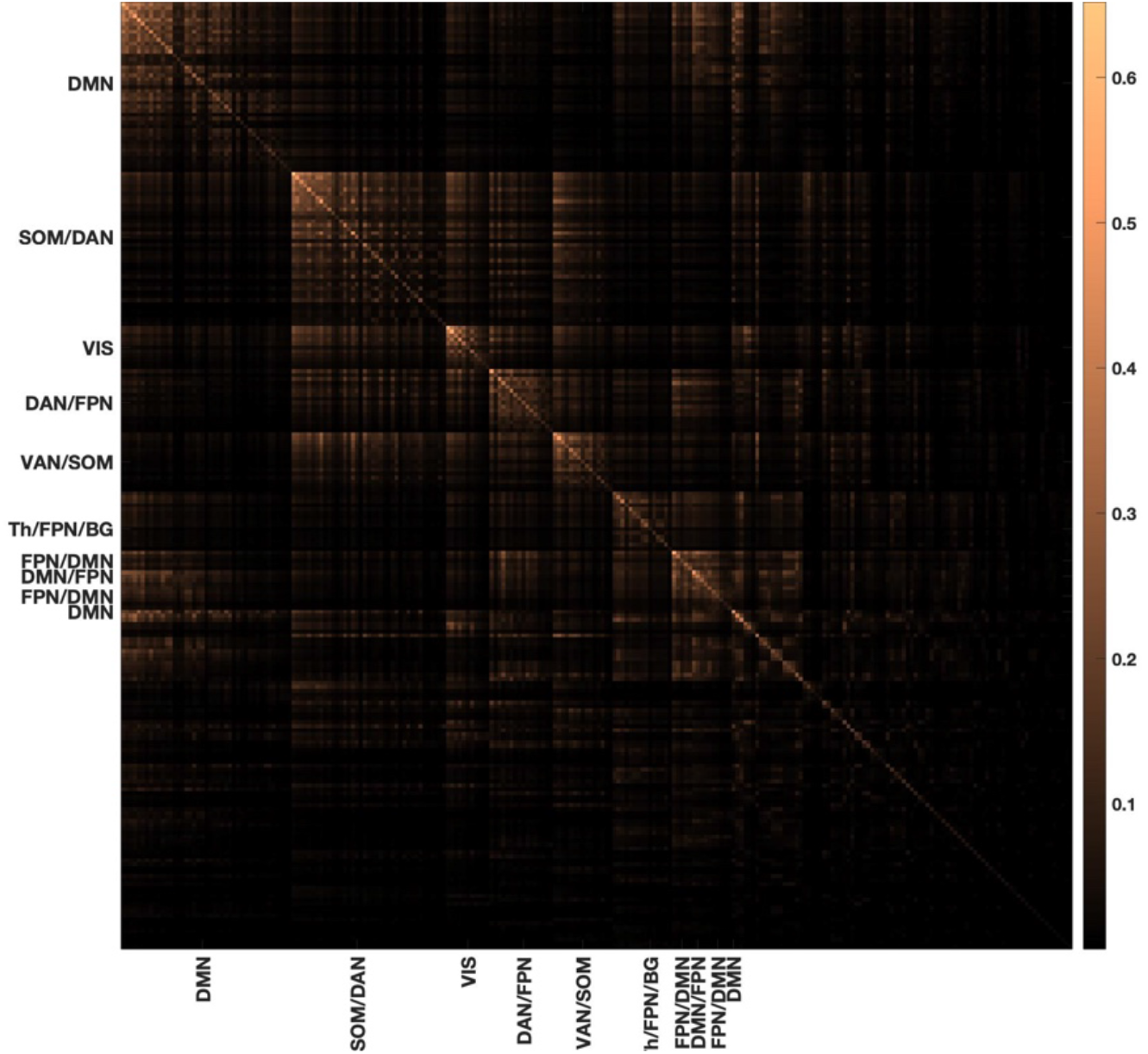
The mean degree of integration between MNs and SNs relative to the full time-span of the fMRI resting-state experiment (q_int_) The SNs (n = 240) are sorted by MN (n = 71), and MNs are sorted by the largest q_int_ value. MNs and SNs are sorted in the same way as in Figure S11.

**Figure S13.**
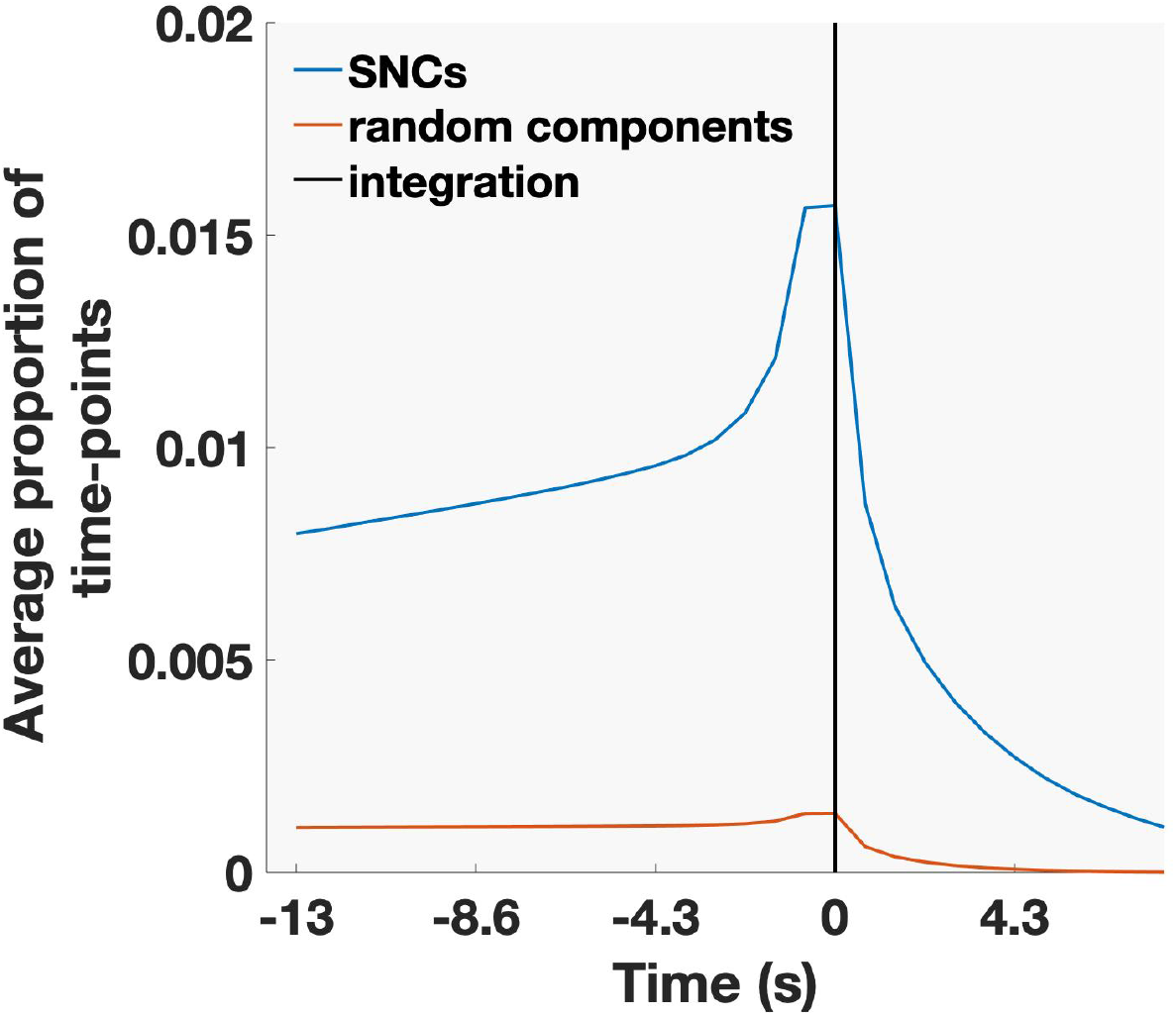
Average sample density plot for all SNCs (blue) and random components (red). The mean was taken across subjects and subnetwork components. Zero on the x-axis corresponds to the time-point of integration (black vertical line). For each time point the plot shows the average proportion of the total number of time-points that the calculation of coherence was based on. The minimum interval between instances of integration as well as duration of integration was one time-point. The highest number of samples were found directly prior to and at the time-point of integration since episodes of both long and short durations contributed to these points in time. This effect was much less pronounced for random components since both intervals between episodes of integration as well as duration of integration were more homogenous, i.e. short durations of integration separated by long intervals of disintegration. This was a consequence of the fact that random components had over all lower values of q_int_ than empirical SNCs.

**Figure S14.**
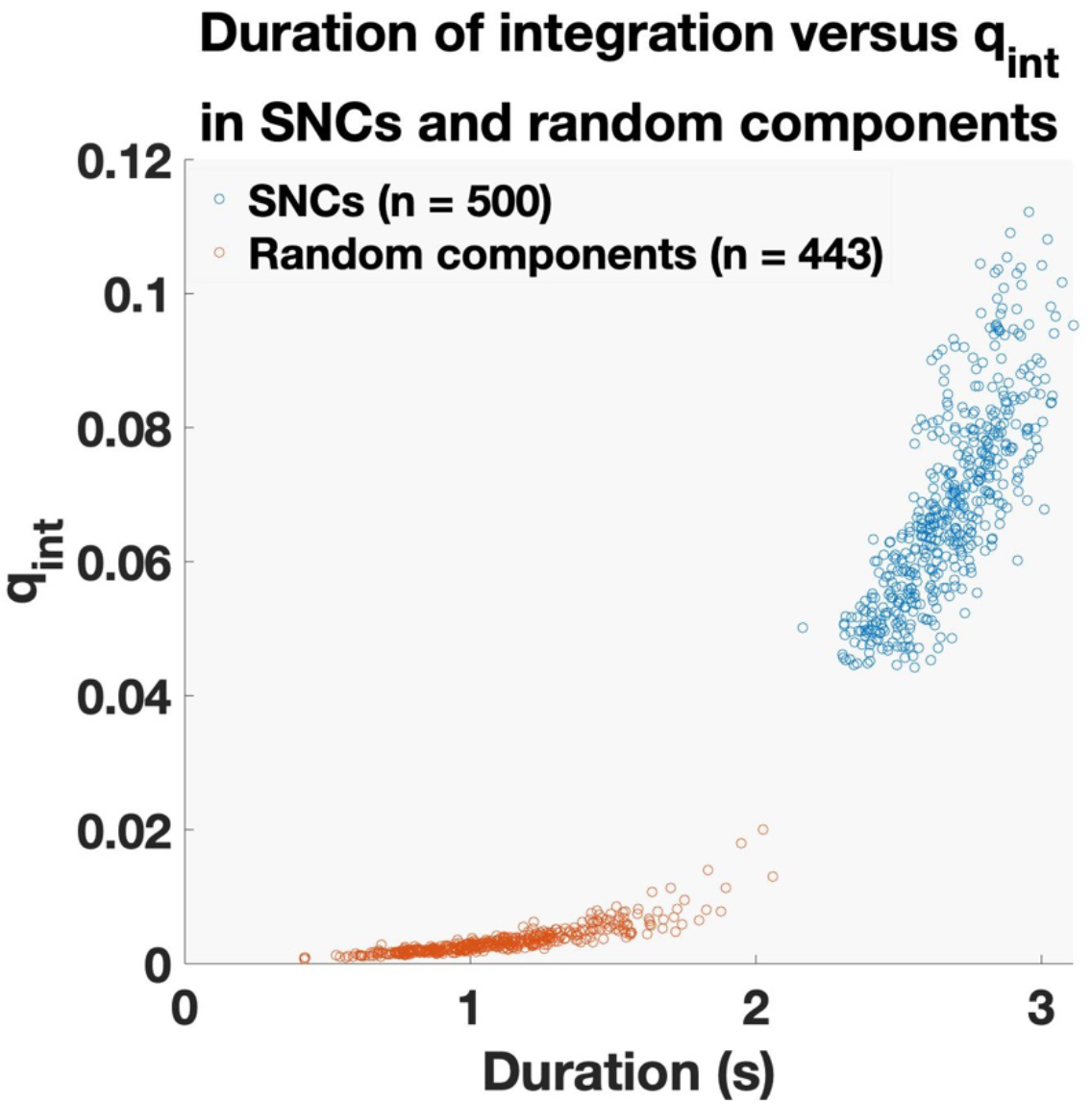
Mean duration of integration of empirically derived subnetwork components (SNCs) and random components vs relative frequency of assignment to the same community (q_int_). Blue dots signify SNCs (n = 500) and red dots represents random combination of eight areas (n = 443). SNCs had a significantly longer duration of integration compared to random components (SNC mean = 2.6 s (SD = 0.3) vs random components mean = 1.1s (SD = 0.3)). The SNCs were drawn from the top two thirds of the full distribution of the SNCs resulting in the relatively sharp lower bound of the distribution (blue).

**Figure S15.**
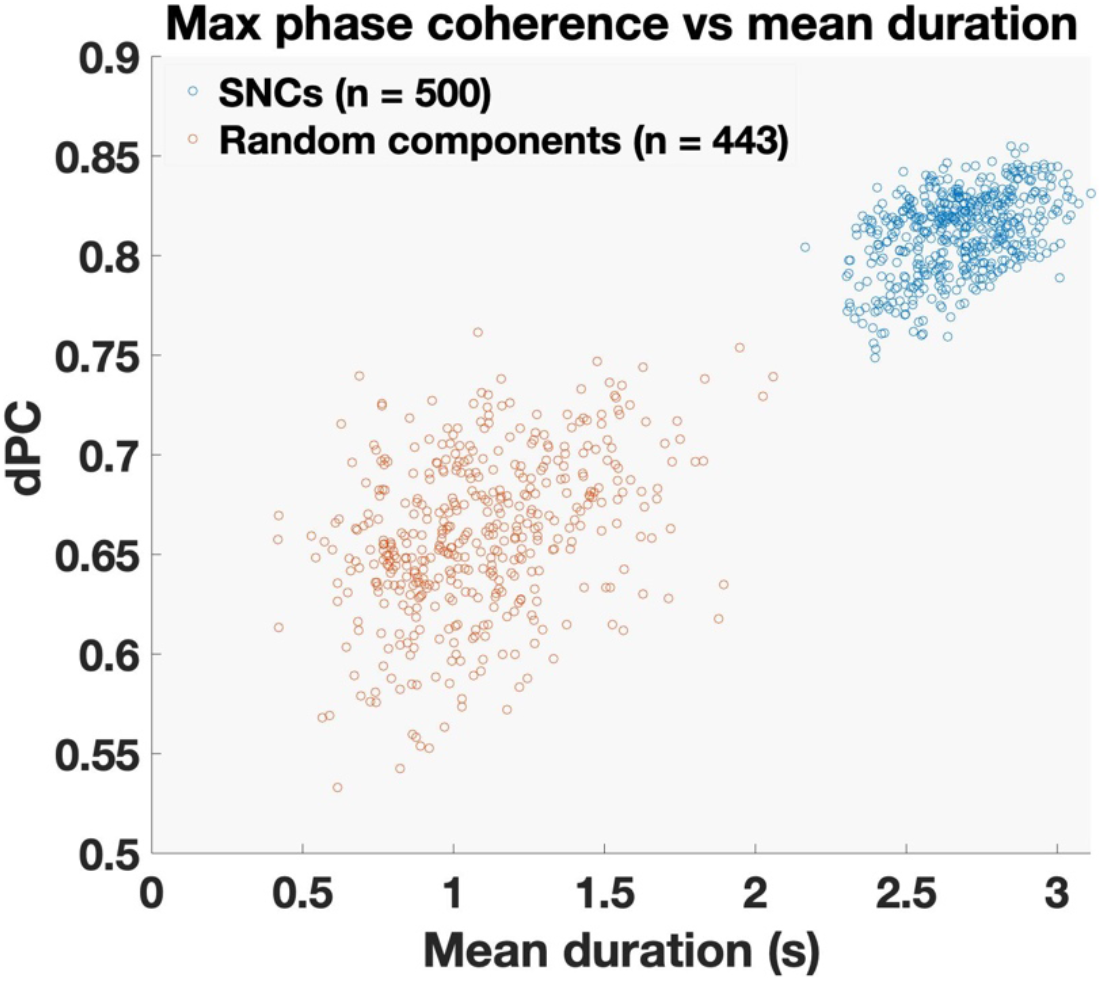
Mean duration of integration versus peak in phase coherence for empirically derived SNCs and randomly generated components (SNCs (n=500) marked in blue, randomly generated components (n=443) in red.

**Figure S16.**
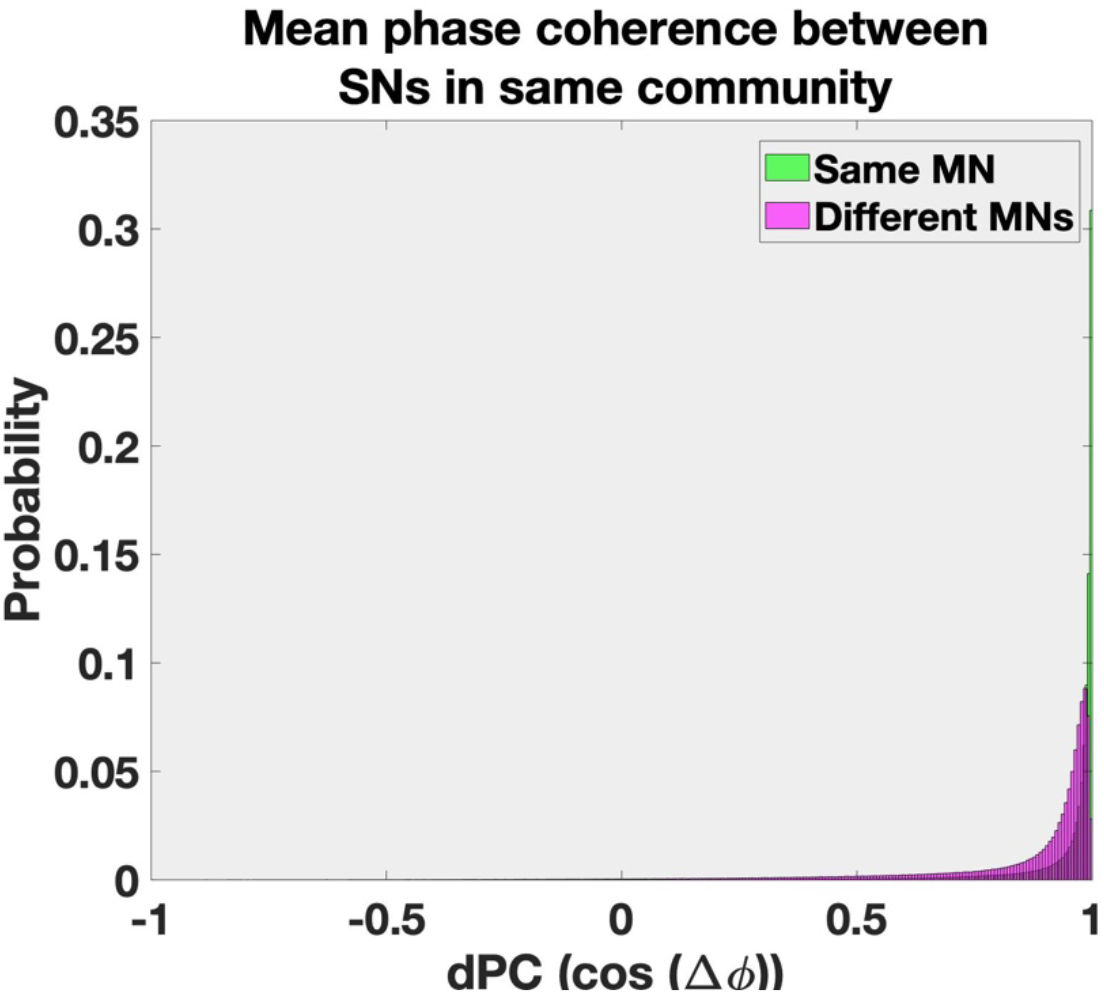
The mean degree of phase coherence between integrated subnetworks (SNs) that either resided in the same meta-network (MN) (green) or in different meta-networks (purple). Here the full distribution before averaging across MNs, communities, time-points and subjects is depicted.

**Figure S17.**
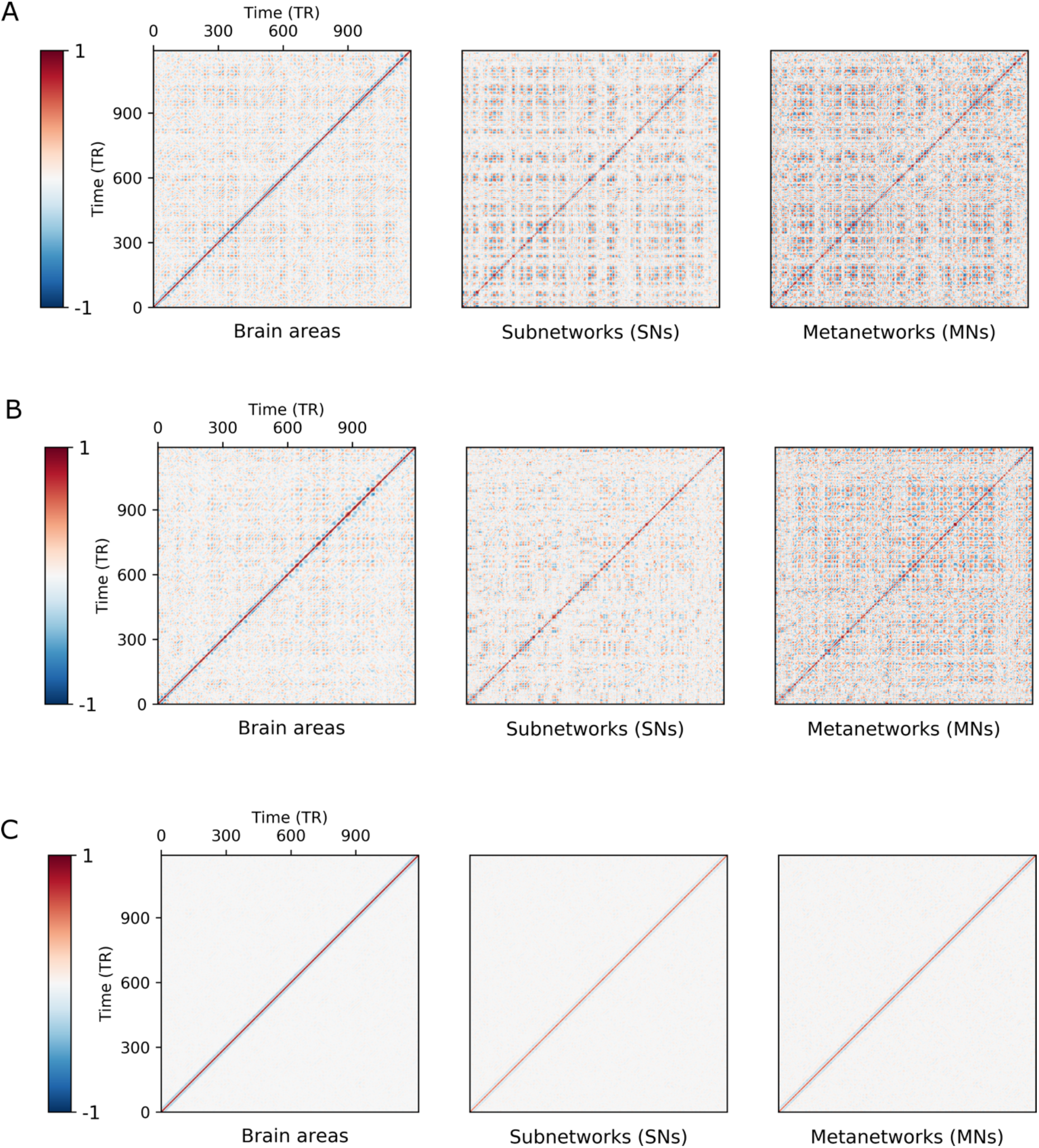
Quasi-cyclic recurrence across all time-points in the acquisition for two individual subjects (A and B) and an averaged across all subjects (C) spanning the entire fMRI resting-state experiment (1200 time-points). Since each subject have their own internal timing only the diagonal expresses strong correlation on the group level, i.e. the time-point to time-point change.

**Figure S18.**
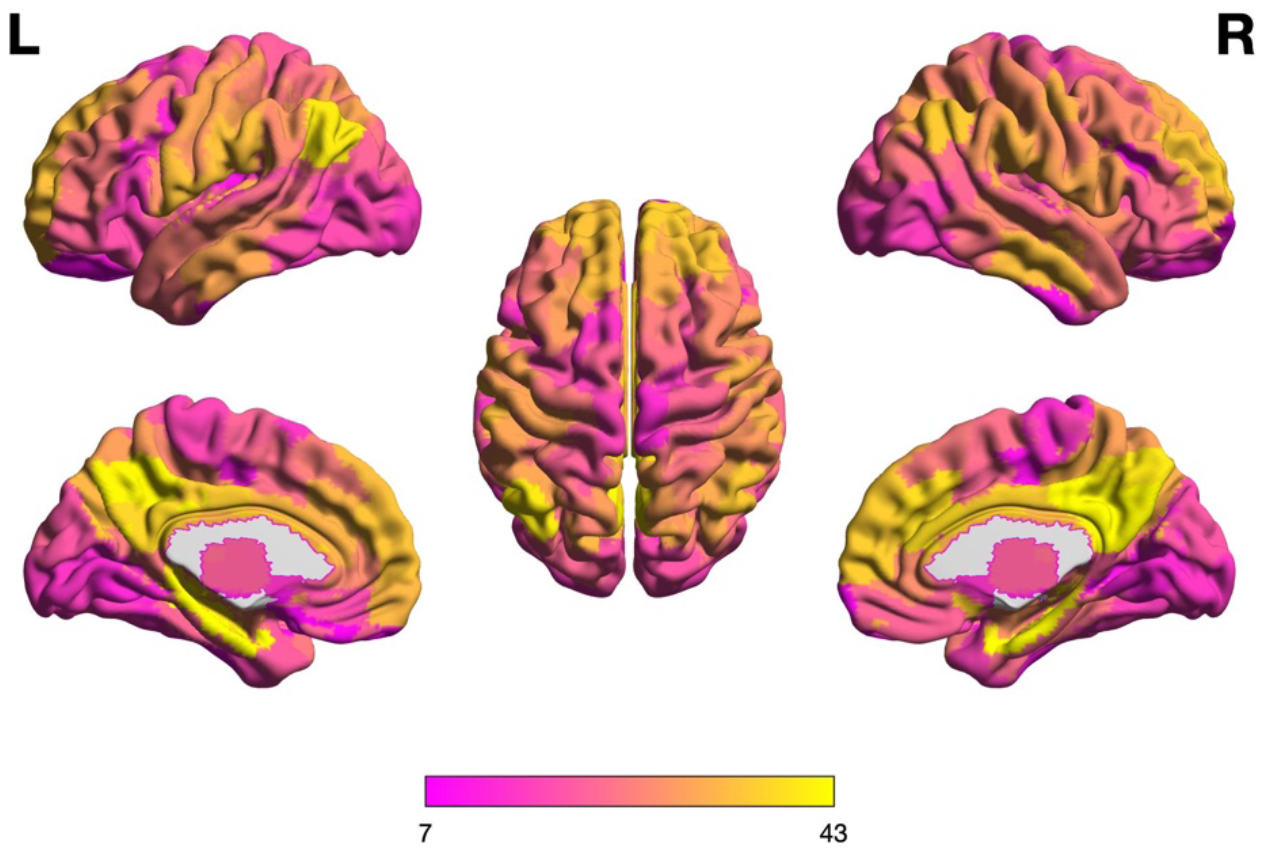
Spatial diversity (p_div_) across all brain areas. Color scale: p_div_*100. Bilaterally the precuneus/posterior cingulate cortex and para-hippocampal gyri stands out (max P_div_). In general, the weights were approximately symmetric across hemispheres.

**Figure S19.**
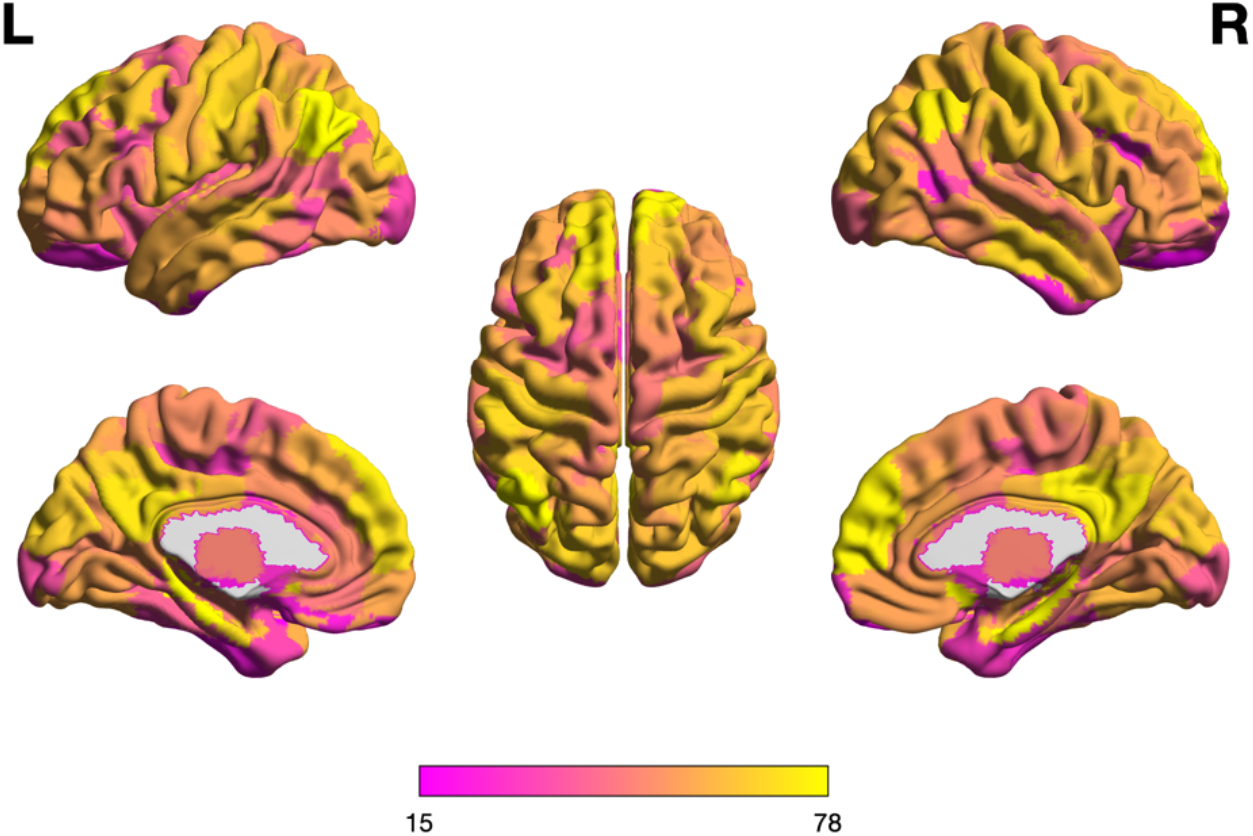
Temporal representation (p_int_) across all areas in the brain. Color scale: p_int_*100. Bilaterally precuneus/posterior cingulate cortex, para-hippocampal gyri, parts of the medial prefrontal cortex and parietal cortex have the highest weights. Similar to p_div_, the weights were approximately symmetric across hemispheres.

**Figure S20.**
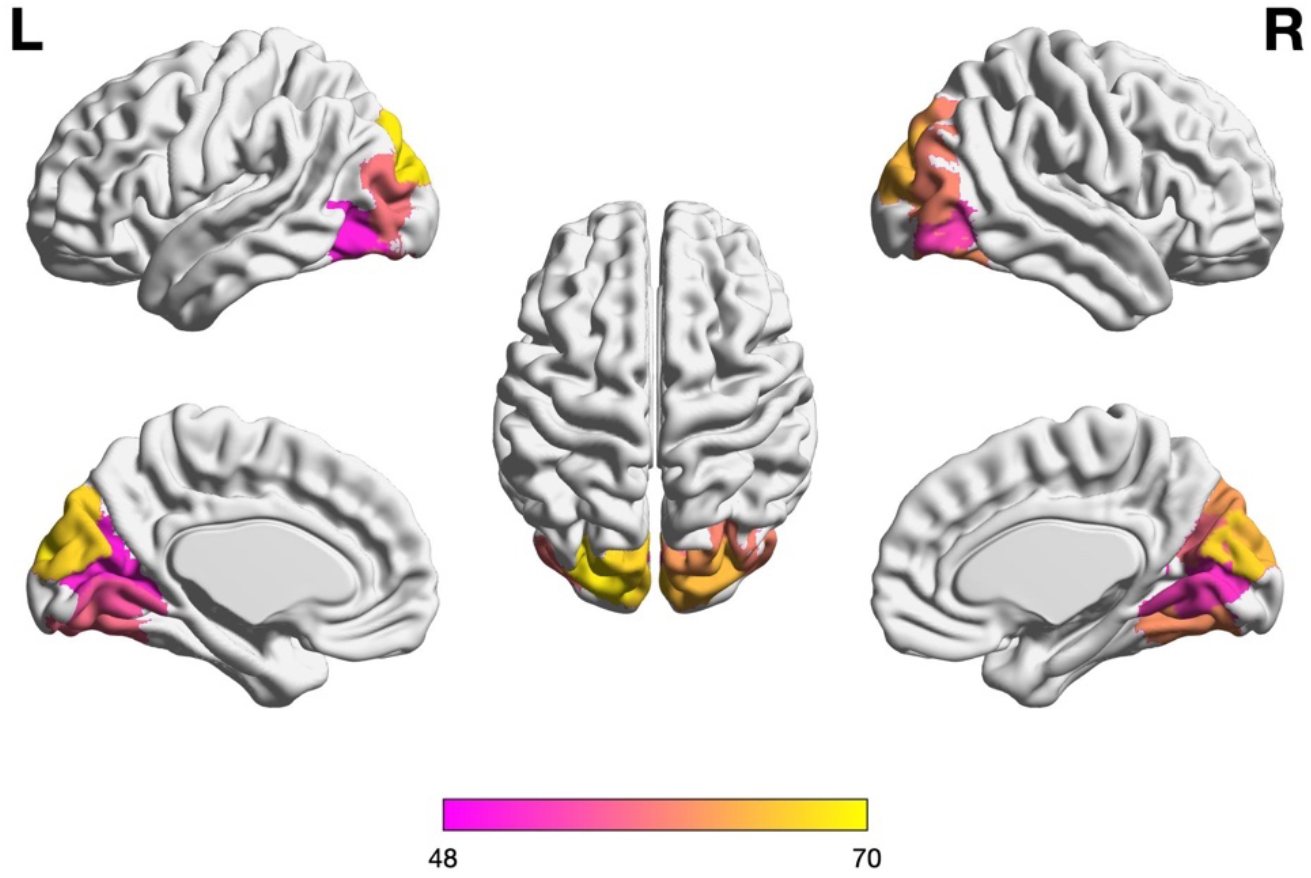
Modular part of the visual network (according to Schaefer 7 parcellation). The band of modular areas aligns perpendicular to the cytoarchitectonic division of primary and secondary visual areas. The weights (Pint *100) were distributed in a symmetric fashion across hemispheres.

**Figure S21.**
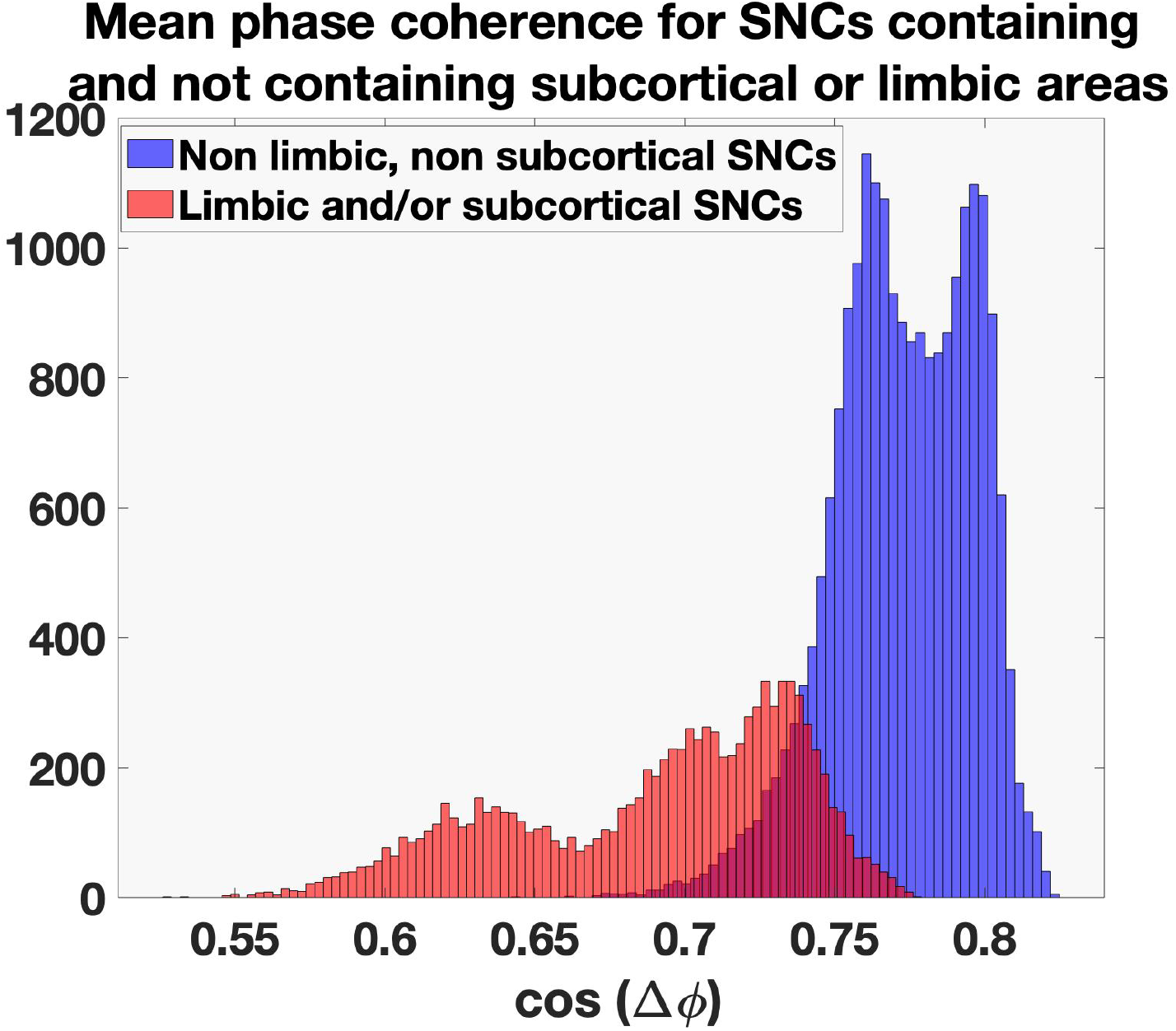
Difference in internal phase coherence for SNCs that contained subcortical and/or limbic areas, either exclusively or mixed with cortical areas (marked in red) compared to SNCs that only contained cortical areas (blue). Clearly the SNCs with subcortical/limbic areas were on average less internally coherent at time-points of integration than SNCs that consisted of cortical areas only.

**Figure S22.**
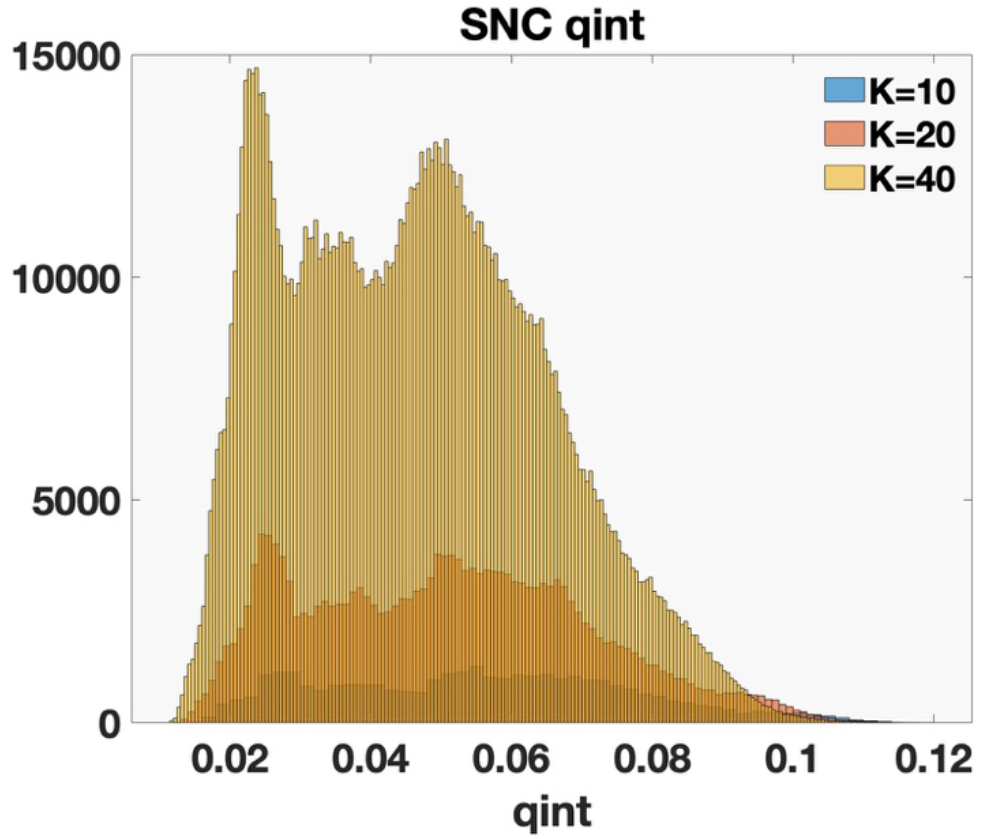
Distribution of SNC q_int_ values for different settings of the expanding factor K = 10 (blue, n = 31976), K = 20 (red, n = 191683) K = 40 (yellow, n = 1257511). See also Suppl. Table S1.

**Figure S23.**
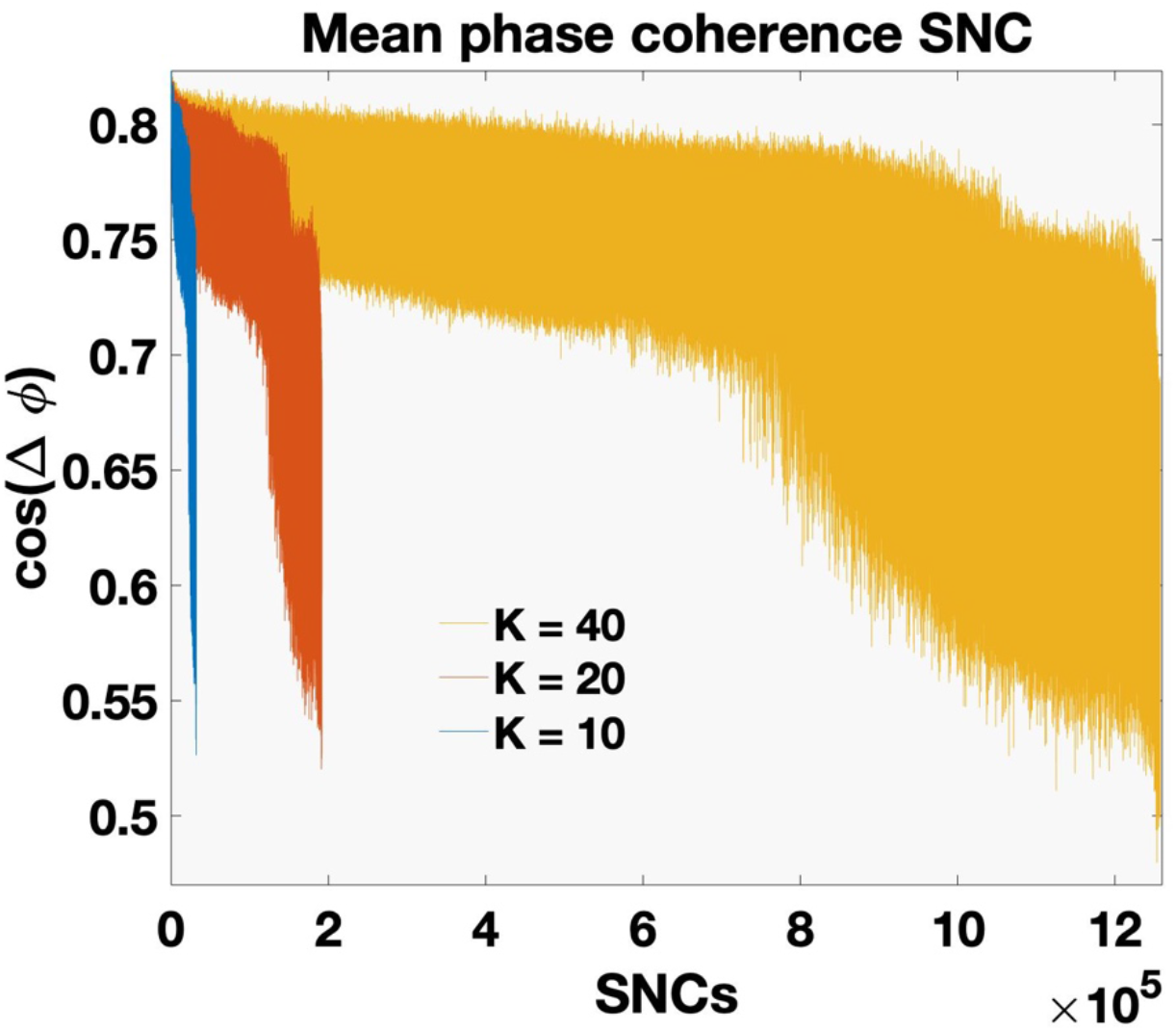
Mean phase coherence (dPC i.e. *cos*(Δ *ϕ*)) of SNCs for different settings of K. The proportion of SNCs with low mean phase coherence (dPC < 0.7), increased with larger K (K = 10 → 0.15, K = 20 → 0.18, K = 40 → 0.20). However, the absolute number was close to exponentially larger due to the total number of SNCs sampled with the doubling of K.

**Figure S24.**
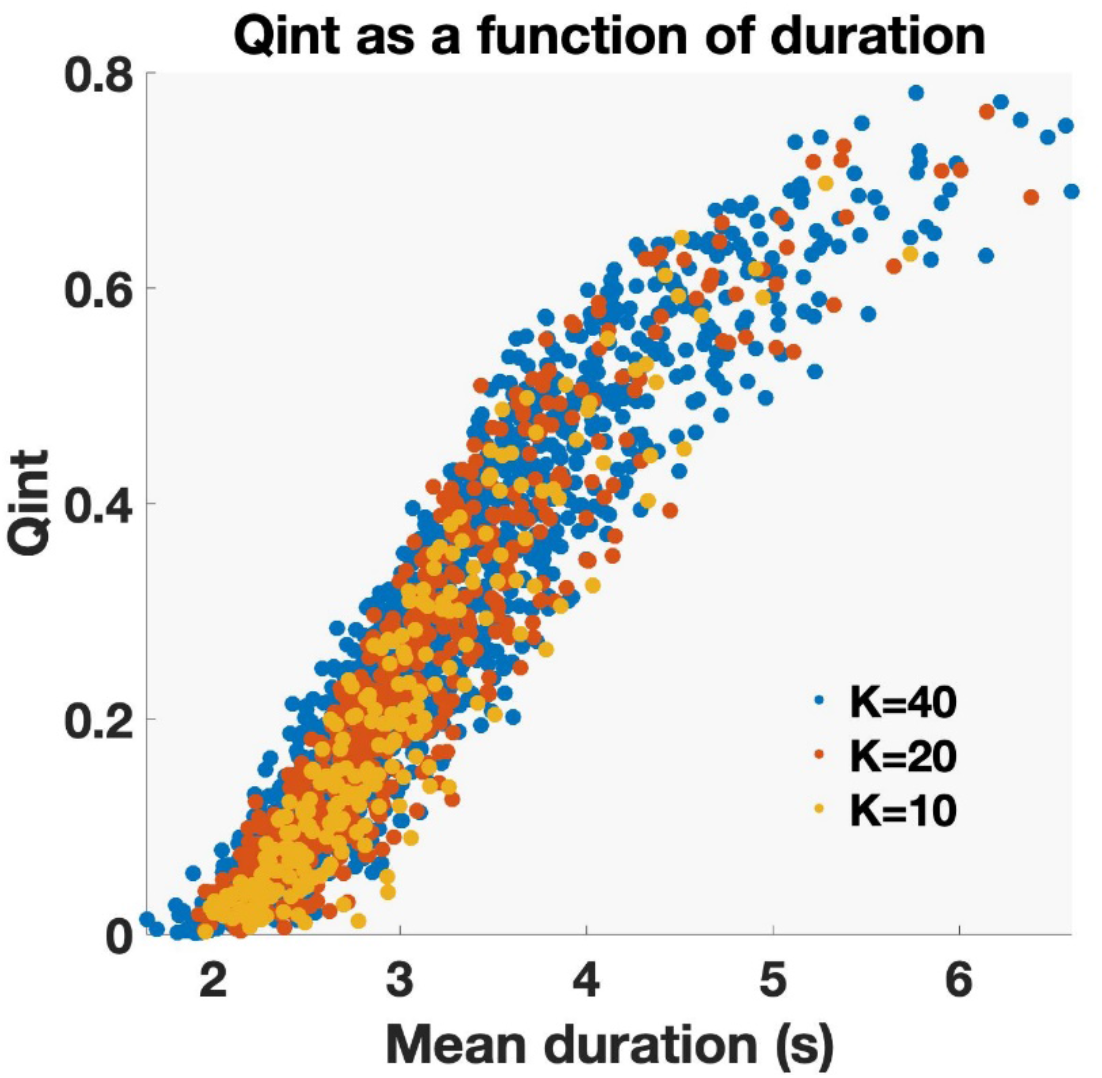
The relationship between q_int_ and duration was stable across choices of K.

**Figure S25.**
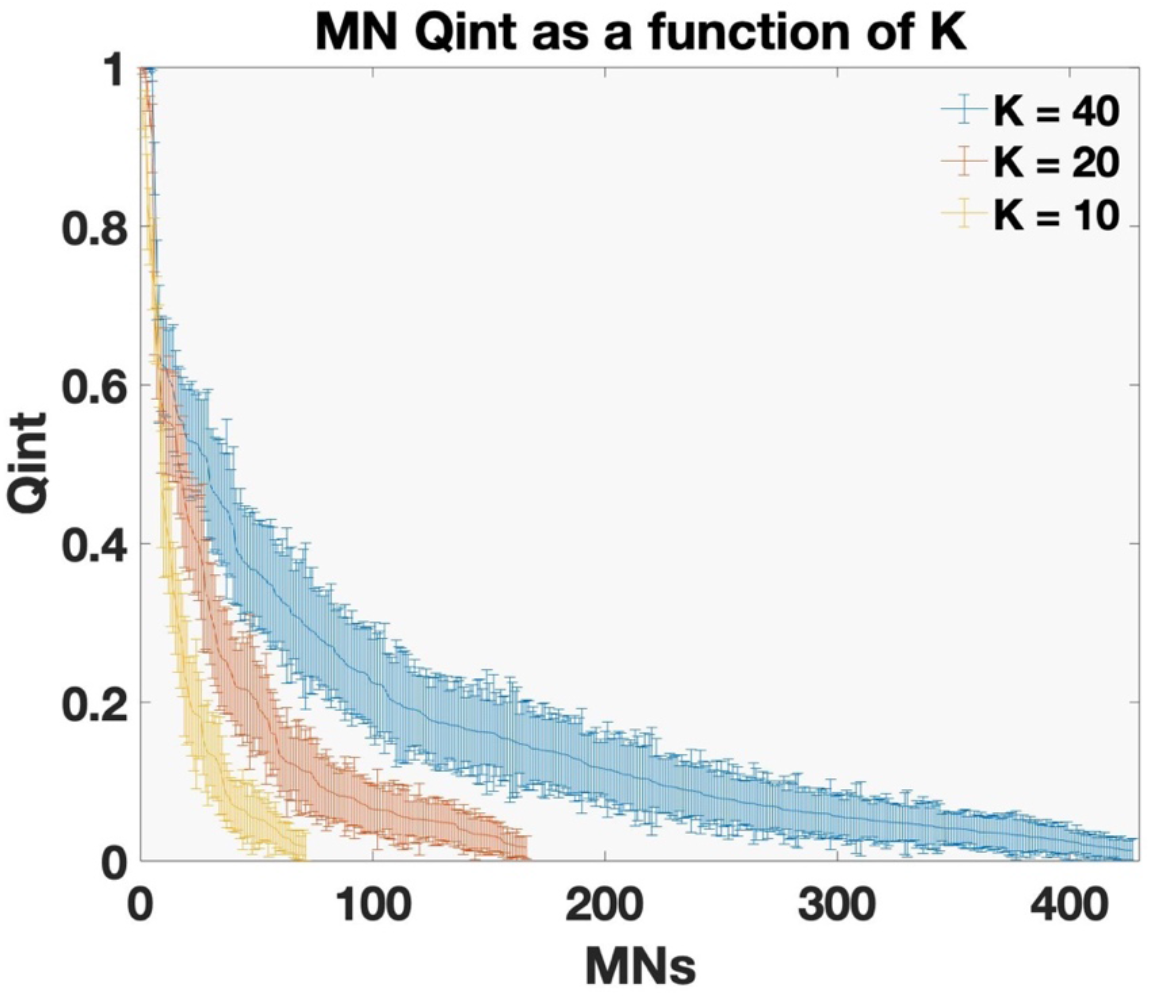
Mean q_int_ for MNs (standard deviations as error bars) for increasing K. Number of MNs with the highest q_int_ remained stable. The small decrease in mean q_int_ with increased K was not statistically significant (p > 0.5) (K = 10 mean MN q_int_ = 0.20 (SD = 0.04), K = 20 mean MN q_int_ = 0.18 (SD = 0.04), K = 40 mean MN q_int_ = 0.17 (SD = 0.04).

**Figure S26.**
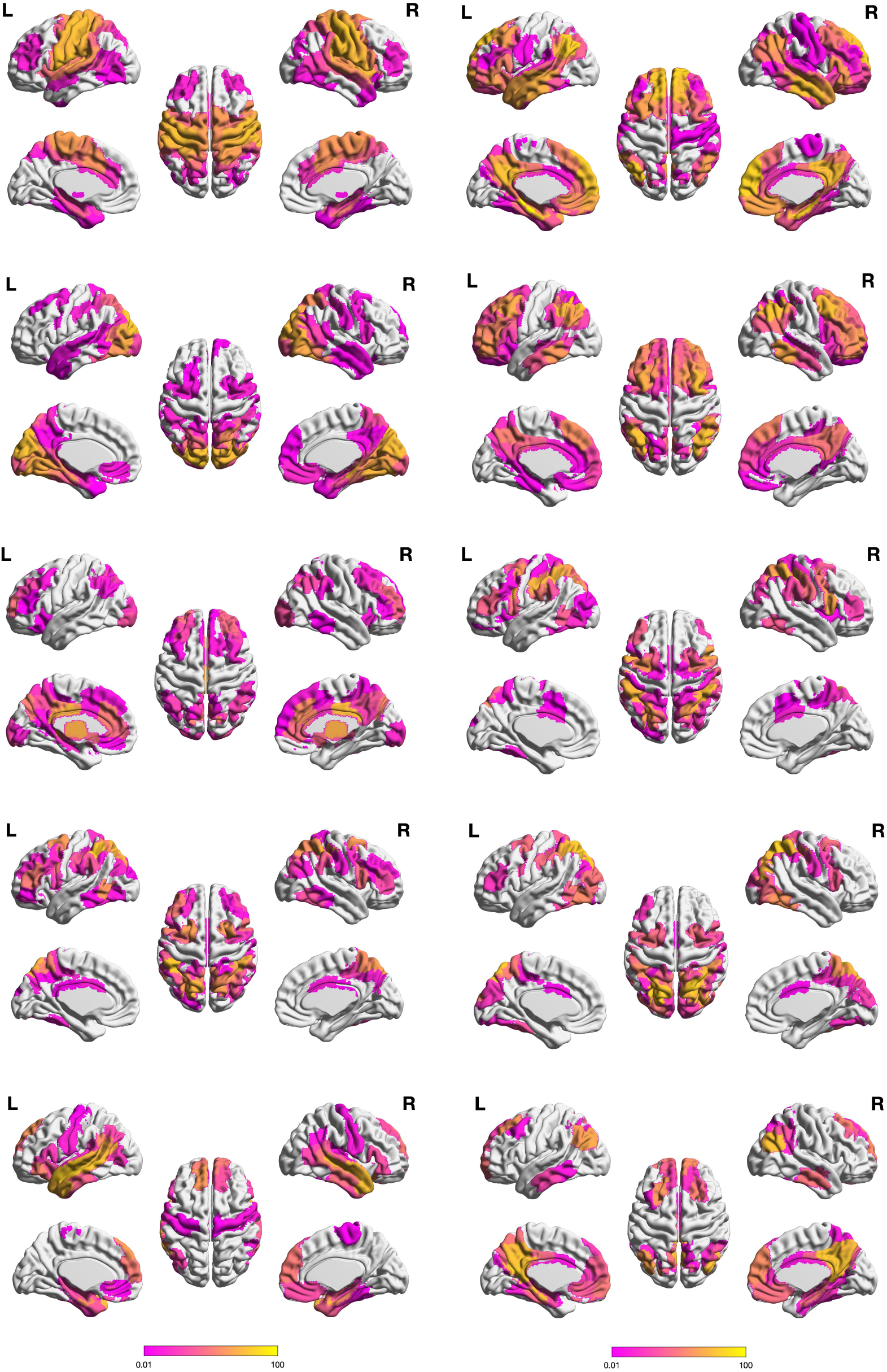
Top ten MNs (K = 20)

**Figure S27.**
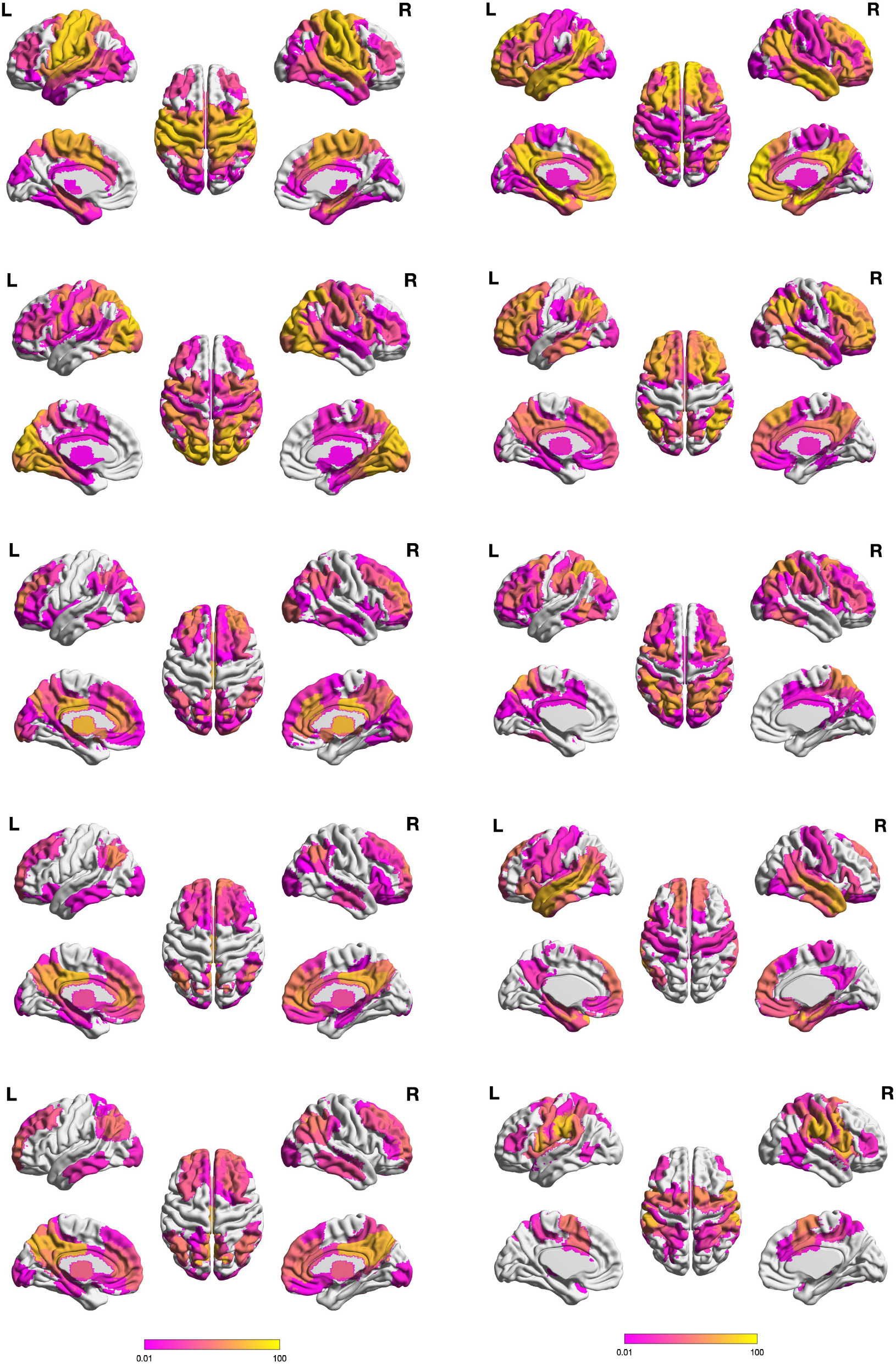
Top ten MNs (K = 40).

**Figure S28.**
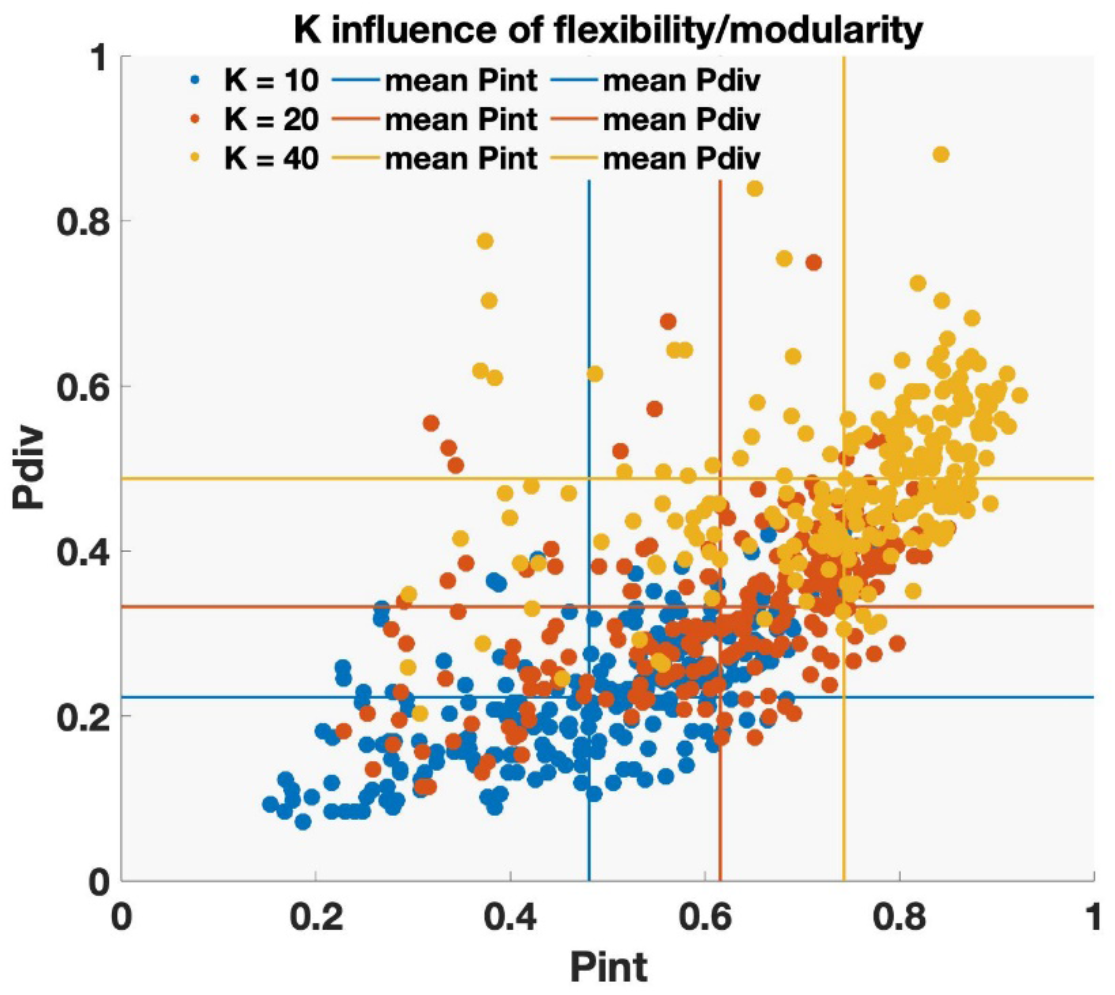
Increase in K result in more SNCs which moves the distribution up and to the right towards higher values in p_div_ and p_int_. Given the increase in number of SNCs with low phase coherence (dPC < 0.7) (Suppl. Figure S23), thresholding at dPC > 0.70 (i.e. < 45°) could be a strategy to prevent overestimation of p_div_ and p_int_ for large K. This would also help with some of the blurring between MNs seen in Suppl. Figure S27.

**Figure S29.**
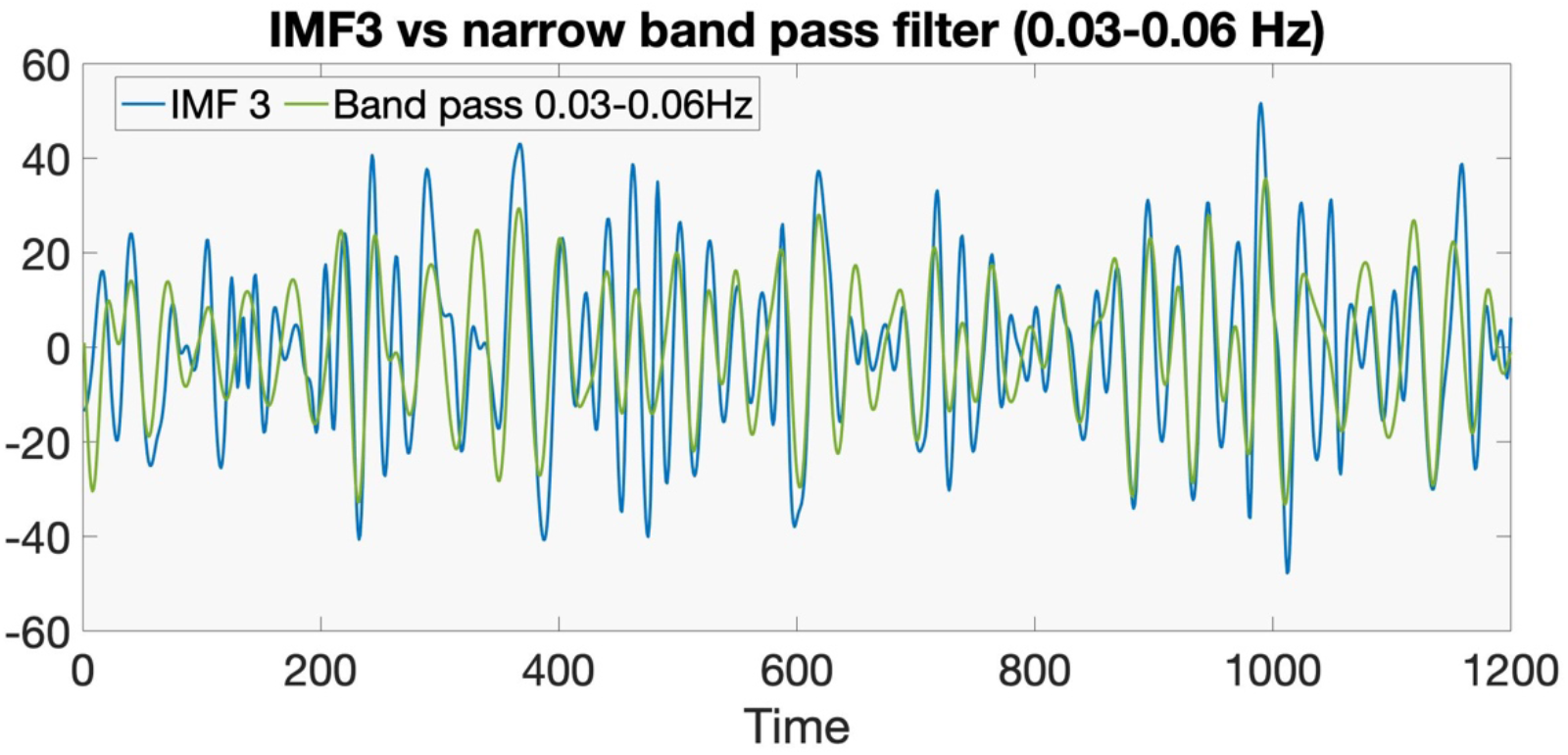
Example of the degree of similarity between the BOLD IMF3 (blue) signal and narrow band-pass filtered signal (0.03-0.06Hz) taken from one random brain area in one subject.

**Figure S30.**
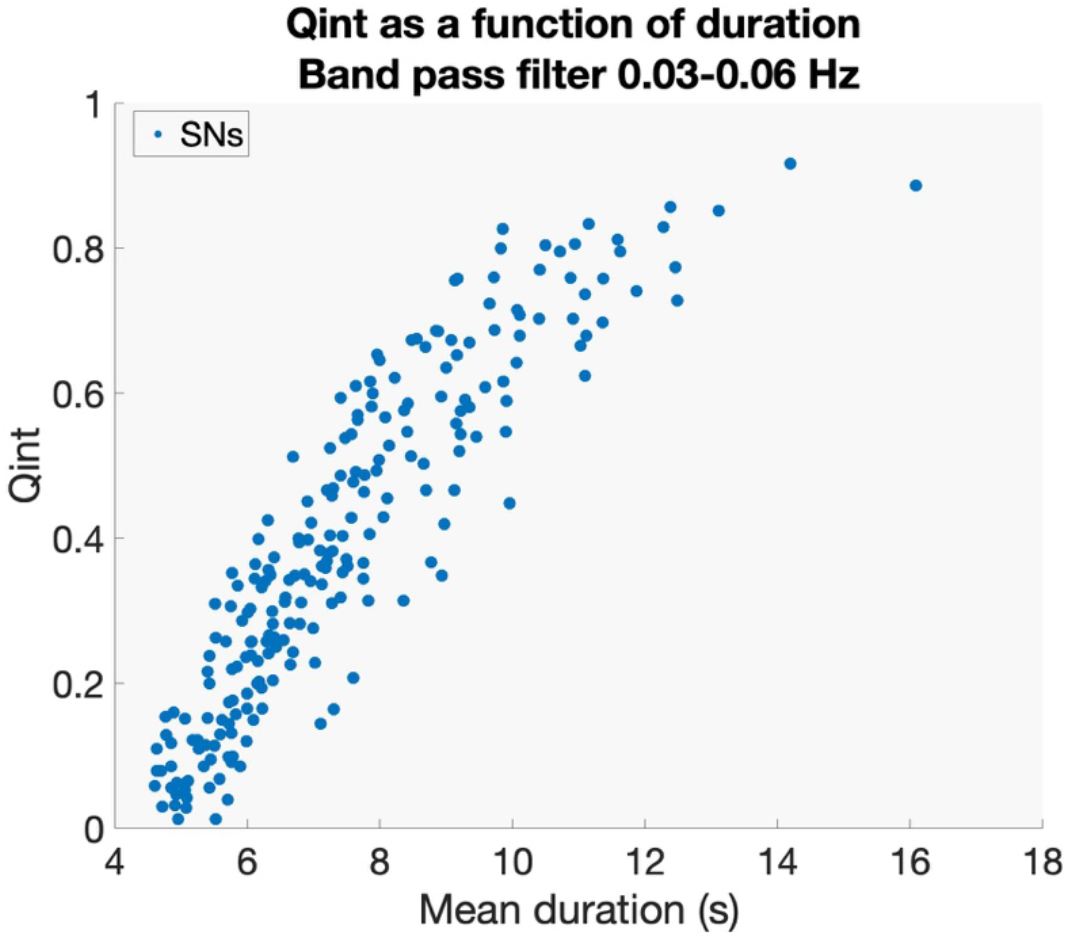
Mean duration of integration as a function of q_int_ for SNs using the band-pass filtered signal. Compare to Suppl. Figure S10 that depicts the same relation computed based on IMF3.

**Figure S31.**
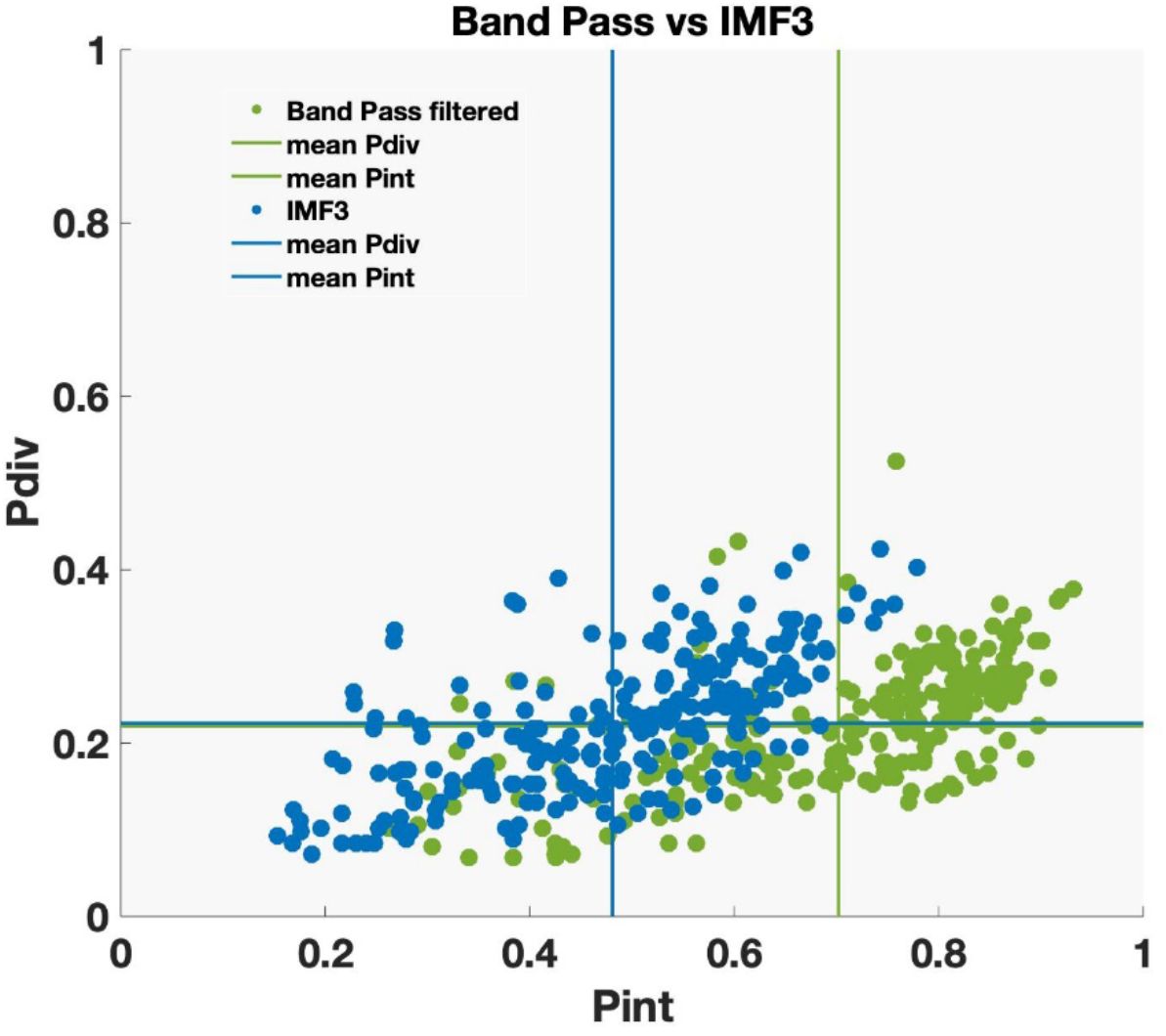
**The** degree of flexibility and modularity for each brain area computed either from the IMF3 time-series (blue dots) or a narrow band-pass filtered signal time-series (green dots). For the band-pass filtering option, we observe a general tendency of increased modularity.

**Figure S32.**
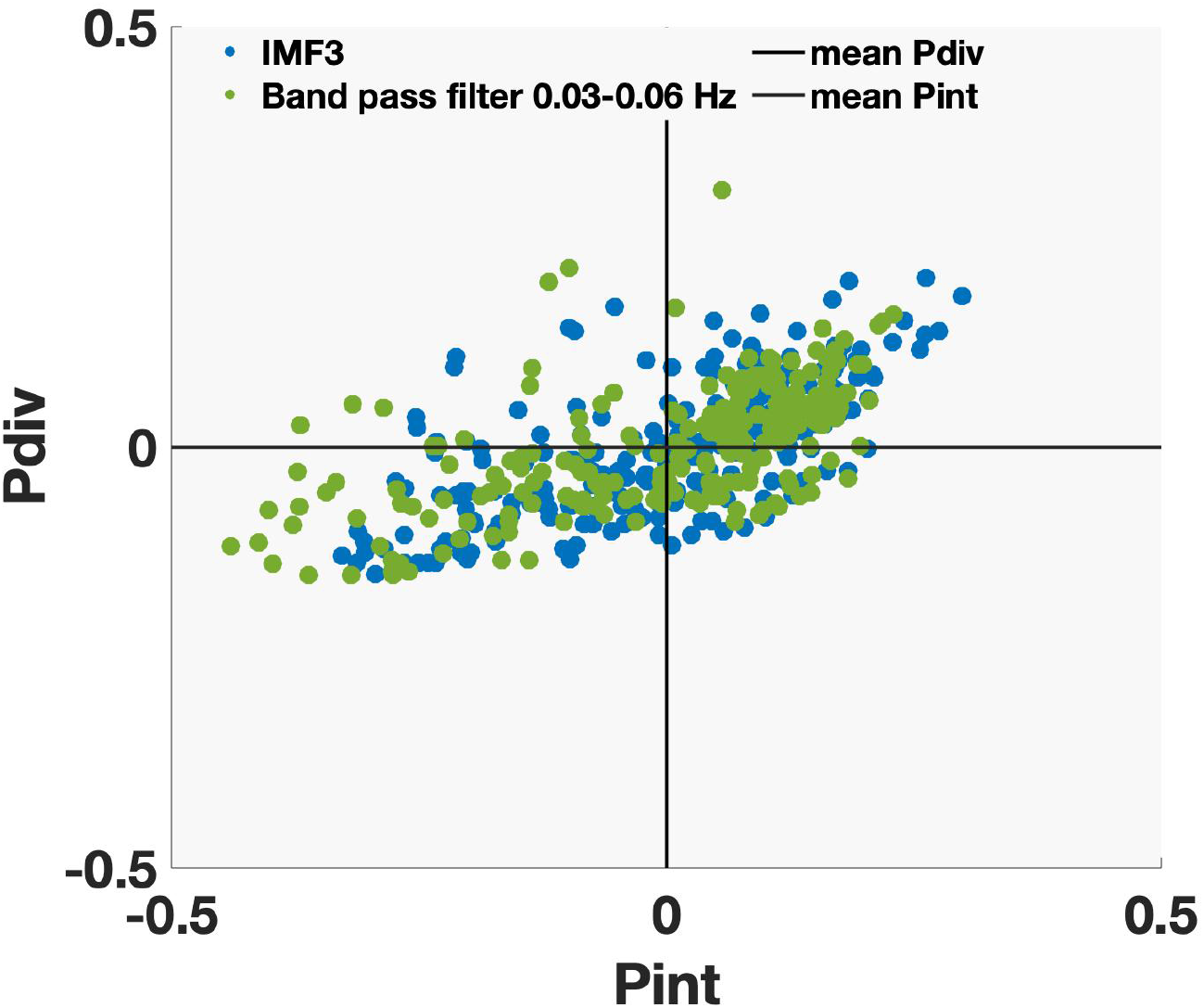
Normalized (demeaned) version of Figure S31. The distribution of areas relative to each other was largely the same for band-bass filtered data and IMF3.

**Figure S33.**
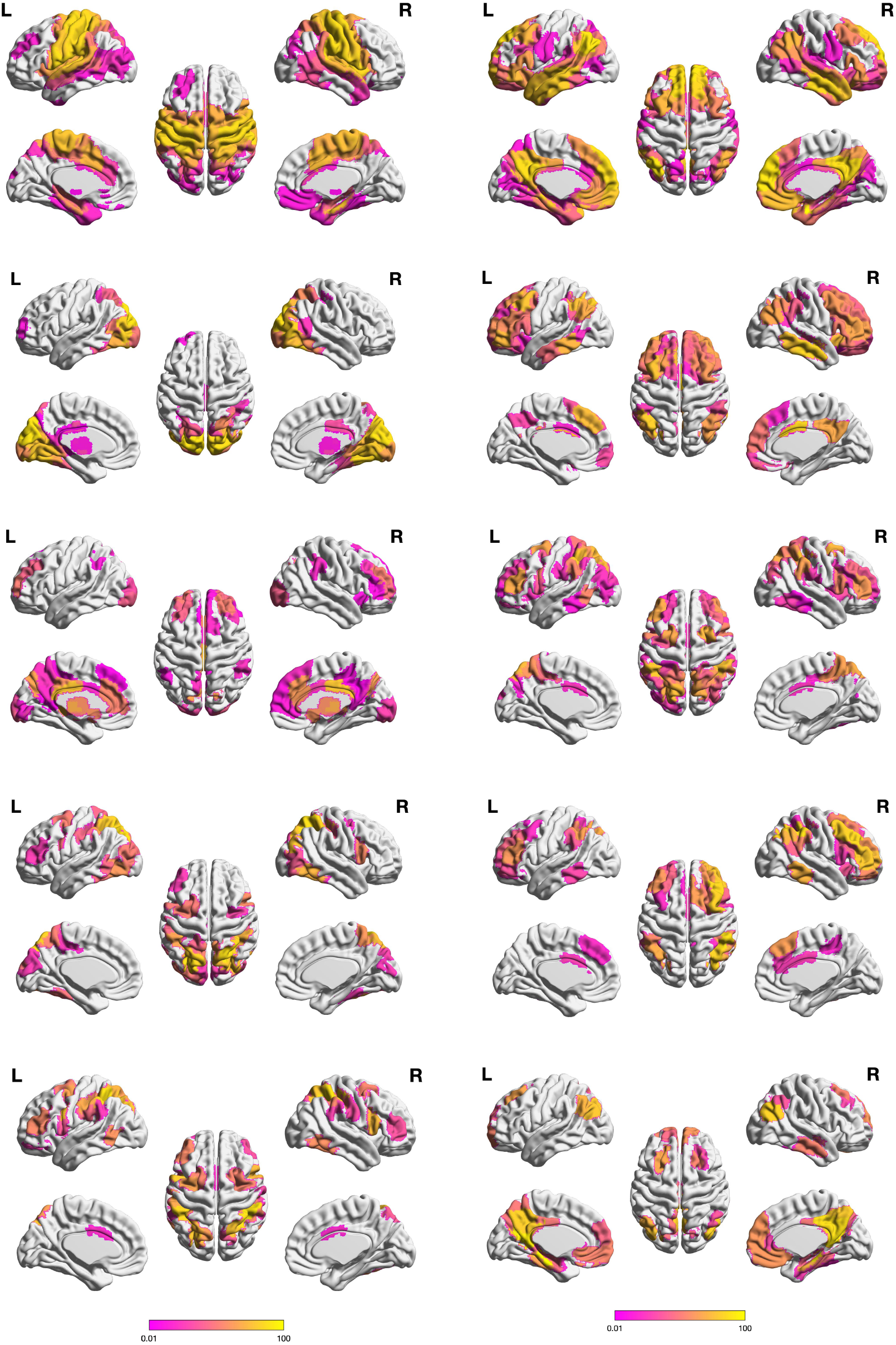
Top ten MNs (using a narrow band-pass filtered signal 0.03-0.06Hz instead of the IMF3 data). Note the increased weights in the maps (brighter yellow) compared to Figure 2.

**Figure S34.**
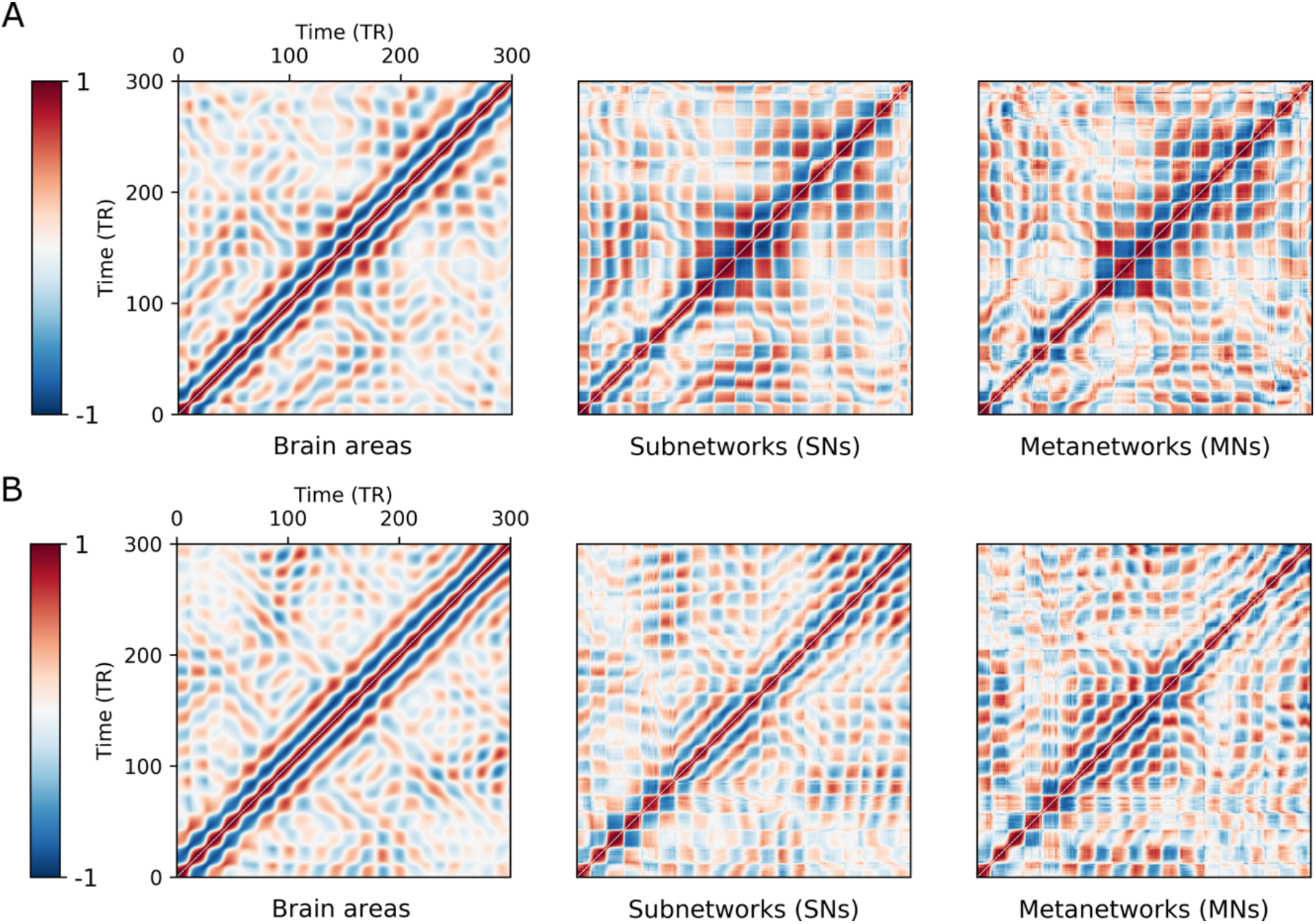
Quasi-cyclic recurrence of state vectors for areas time series, SNs and MNs (narrow band pass filtered signal 0.03-0.06Hz) Example of recurrence plots of time-resolved state vectors across time for two different subjects (panels A and B). The same two subjects are used as in Figure 4 (BOLD IMF3). For increased clarity, each plot is a zoomed-in version focusing on the first 300 time points (out of a total of 1200 time points). Recurrence was more pronounced for band-pass filtered compared to IMF3 based state vectors across granularity levels, for details see Supplementary Text.

**Table S1.** SNC q_int_ summary statistics for choices of the expansion factor K. (K = 10, 20, 40). The range of q_int_ values remain unchanged across different settings of the factor K.

| SNC Q<sub>int</sub> |  |  |  |  |
| --- | --- | --- | --- | --- |
|  | median | iqr | max | min |
| **K=10** | 0.06 | 0.03 | 0.12 | 0.01 |
| **K=20** | 0.05 | 0.03 | 0.12 | 0.01 |
| **K=40** | 0.05 | 0.03 | 0.12 | 0.01 |

**Table S2.**
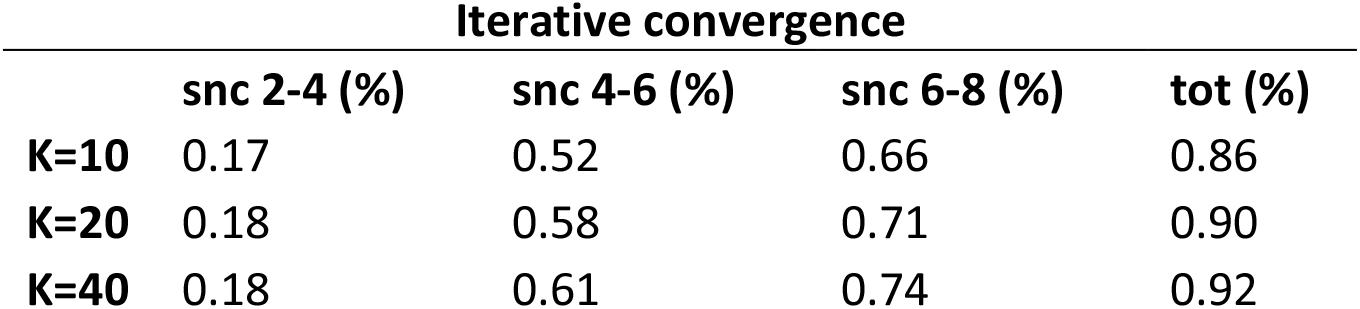
Iterative convergence for different K. Convergence at level n was defined as one minus the proportion between the unique number of SNCs and all SNCs resulting from the seeds at level n-2.

**Table S3.** Number of SNs and MNs for different settings of the expansion factor K.

| Number of Network Units |  |  |  |
| --- | --- | --- | --- |
|  | SNs |  | MNs |
| **K=10** | 240 |  | 71 |
| **K=20** | 578 |  | 166 |
| **K=40** | 1556 |  | 427 |

**Table S4.** Mean duration of SNs as a function of the expansion factor K

| Mean (SD) duration of<br>SNs (seconds) |  |
| --- | --- |
| **K=10** | 2.9 (0.7) |
| **K=20** | 3.0 (0.7) |
| **K=40** | 3.1 (0.8) |

